# Wntless interacts with Notch signaling to balance the generation of neurons and gliocytes in vertebrate dorsal diencephalon

**DOI:** 10.1101/2025.08.06.668881

**Authors:** Shu-Heng Lin, He-Yen Pan, Bo-Tsung Wu, Joe Sakamoto, Chun-Hsiu Wu, Atsuko Shimada, Yasuhiro Kamei, Hiroyuki Takeda, Yung-Shu Kuan

## Abstract

Wnt chaperon Wntless (Wls) mediates the intracellular transport of Wnts and plays important roles in early vertebrate brain development. Spatially restricted induction of Wls denotes the earliest differentiation of non-telencephalic cells in human brain organoids. In zebrafish developing diencephalon, loss-of-Wls reduces the formation of habenula (HA) neurons but how Wls influences HA neurogenesis is unclear and whether Wls regulates gliocyte development is unknown. Here we report that the formations of cholinergic, substance P-ergic or glutamatergic neurons in HA are reduced differentially but the generation of gliocyte-derived choroid plexus (ChP) epithelia is increased in *wls* null mutants. At earlier stage, three-dimensional gene expression analyses revealed that while *neurog1* expressions in HA progenitor zones are reduced, the expressions of Notch downstream effector *her6* are increased and expanded into HA progenitor zones in *wls* mutants. Over-expressing Her6 in *neurog1*-positive cells reduced *neurog1* expressions in HA progenitor zones. These results indicate that Wls restricts the expressions of Notch effector *her6* to promote the specification of *neurog1* proneurons and demotes the generation of ChP epithelia in zebrafish embryonic dorsal diencephalon.

## Background

The developing dorsal diencephalon (or epithalamus) in vertebrate brain contains two major neuronal clusters, the habenular nuclei (HA) plus pineal organ (P), and one gliocyte (glial cell) containing cluster, the choroid plexus (ChP) (Liu, Zhou et al., 2018). In zebrafish, HA are situated bilaterally to the pineal organ and extend their axons to the interpeduncular nucleus (IPN) to construct one of the conserved forebrain to midbrain conduction systems which modulates aversive response and reward processing in vertebrate brain. Defects in HA-IPN circuitry are implicated in a variety of behavior and neurological disorders such as elevated anxiety, depression or schizophrenia (Beretta, Dross et al., 2012, Bianco & Wilson, 2009, Fore & Yaksi, 2019, Kinoshita & Okamoto, 2023, Roberson & Halpern, 2018) (Figure 1A). The ChP comprises a polarized epithelial cell layer (ChPe) derived from glial cells and blood vessels derived from mesodermal endothelial cells. In humans and rodents, ChP develop at four sites in the roof of the neural tube shortly after its closure, in the order fourth, lateral and third ventricles (Hunter & Dymecki, 2007, Kompanikova & Bryja, 2022, Lun, Monuki et al., 2015, MacDonald, Lu et al., 2021). In zebrafish, the third ventricle ChP in dorsal diencephalon is located right anterior to the pineal organ and can be clearly identified in larvae at 3-4 days post-fertilization (Garcia-Lecea, Kondrychyn et al., 2008, Henson, Parupalli et al., 2014) (Figure 1A). The functions of ChP include producing cerebrospinal fluid (CSF), forming the essential blood-CSF barriers which render mechanical support, providing a route for some nutrients, and removing by-products of metabolism and synaptic activity (Kaur, Rathnasamy et al., 2016, Kompanikova & Bryja, 2022, Kratzer, Ek et al., 2020, Lun et al., 2015). Despite their importance, little is known about how these essential structures develop.

**Figure 1.**
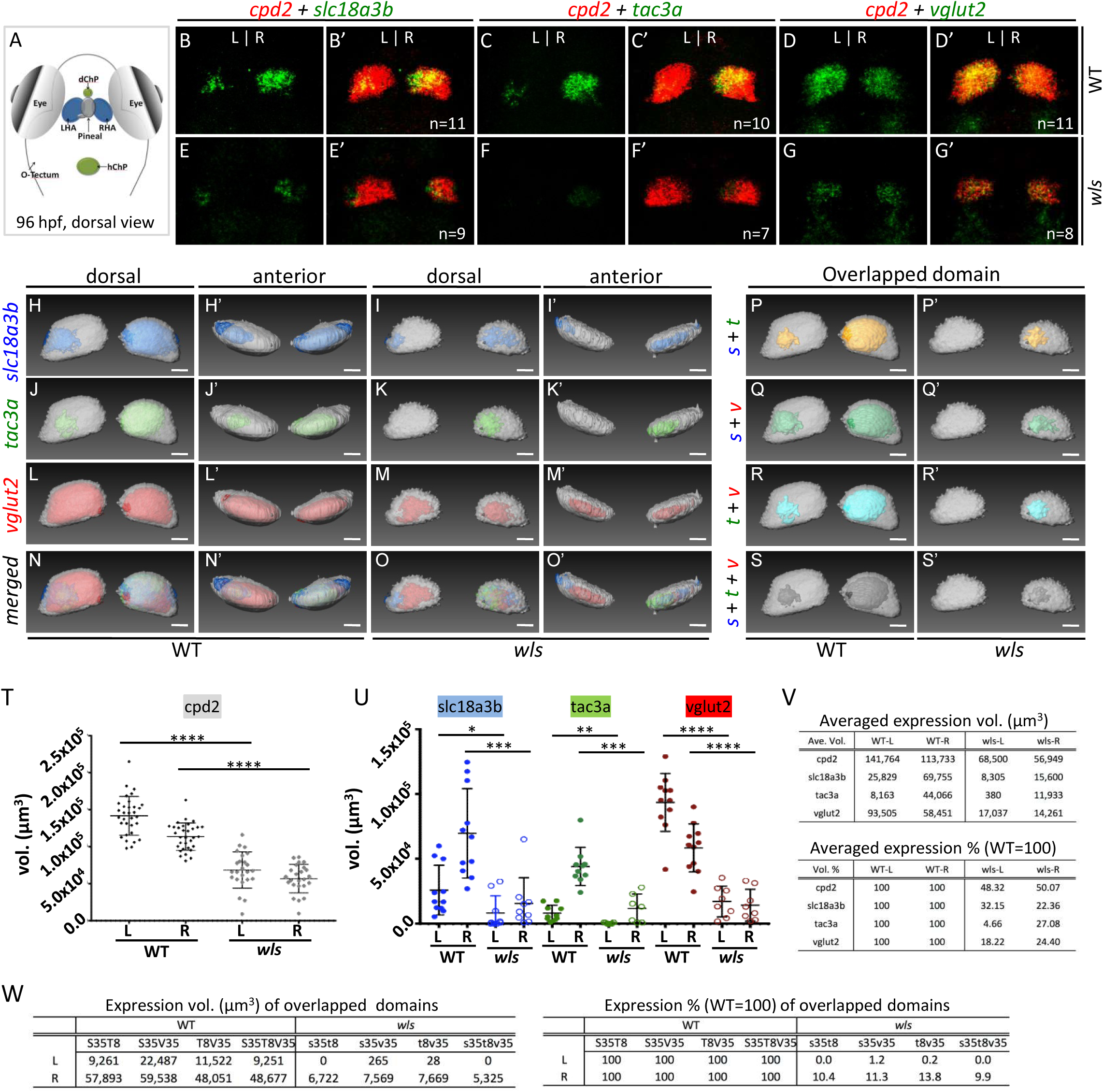
Habenular neuronal fates are changed in *wls* mutants. (A) The schematic cartoon indicating the locations of diencephalic or hindbrain ChPs (dChP or hChP in green), pineal (in gray), left or right HA (LHA or RHA in blue), eyes and optic tectum (O-Tectum). (B-G’) Dorsal views of representative Z-projection images from *slc18a3b* + *cpd2* (B, B’, E, E’), *tac3a* + *cpd2* (C, C’, F, F’), or *vglut2* + *cpd2* (D, D’, G, G’) double mRNA *in situ* labeling in HA of 96h WT (B-D’) or *wls* (E-G’) larva. (H-O’) Dorsal or anterior views of representative snapshots from virtual 3D map showing aligned and averaged Z-stacks of *slc18a2b* + *cpd2* genes (H to I’, in blue and grey), *tac3a* + *cpd2* genes (J to K’, in green and grey) or *vglut2* + *cpd2* genes (L to M’, in red and grey) in WT (H-H’, J-J’, L-L’) or *wls* mutants (I-I’, K-K’, M-M’). (N-O’) Snapshots from merged virtual 3D map of all four genes. Scale bar = 30μm. (P-S’) Snapshots showing overlapped domains for *slc18a3b* + *tac3a* (*s*+*t*; P, P’), *slc18a3b* + *vglut2* (*s*+*v*; Q, Q’), *tac3a* + *vglut2* (*t*+*v*; R, R’), or *slc18a3b* + *tac3a* + *vglut2* (*s*+*t*+*v*; S, S’) gene transcripts from WT (P-S) or *wls* (P’-S’) larvae. (T-V) Averaged expression volumes (Vol.) or percentage (%) of all four genes in WT or *wls* larvae. (W) Expression Vol. or % of overlapped domains in WT or *wls* larvae. Confidence *p* value (*p*) is denoted by “*”. “****” means *P*<0.0001; “***” means *P*<0.001; “**” means *P*<0.01; “*” means *P*<0.05 in t-test. (See Supplemental Figure 1 and Table1 for numeric data)

Wnt and Notch signaling molecules each play their own roles in regulating the generation of HA and ChPs during vertebrate brain development. In mice, Wnt1 and Wnt component Tcf7l2 are involved in HA neuronal differentiation and HA neuronal wiring whereas WNT5a, Wls, beta-Catenin and Notch1 play essential roles in ChP epithelia morphogenesis and proliferation (Carpenter, Rao et al., 2010, Company, Moreno-Cerda et al., 2021, Hunter & Dymecki, 2007, Kaiser, Jang et al., 2021, Langford, O’Leary et al., 2020, Lee, Yoon et al., 2017, Parichha, Suresh et al., 2022). In zebrafish, Wnt components such as *axin1*, *tcf7l2*, *wls*, and Notch activator *mindbomb* (*mib*) modulate the left-right patterning and neurogenesis of HA (Aizawa, Goto et al., 2007, Beretta, Dross et al., 2013, Carl, Bianco et al., 2007, Husken, Stickney et al., 2014, Kuan, Roberson et al., 2015, Roberson & Halpern, 2017). In comparison, no Wnt component has been studied for a role in ChPe development in zebrafish but studies showed that Shh is required for the synchronized outgrowth of ChPe and fenestrated vasculature, depletion of either *delta a* (*dla*) *delta d* (*dld*) or *notch1b* increases ChPe growth, and depletion of *mib* causes defects in specifying ChPe (Bill, Balciunas et al., 2008, Huang, Ketova et al., 2009). Nevertheless, a clear picture of how regional Wnt or Notch activities are established and whether any regulatory mechanism exists to integrate the generation of HA neurons and ChP epithelia is unknown.

Wntless (Wls) encodes a transmembrane protein which is responsible for the secretions of some members of Wnt proteins (WNTs) (Banziger, Soldini et al., 2006, Bartscherer, Pelte et al., 2006, Ching, Hang et al., 2008, Wolf & Boutros, 2023, Wu, Wen et al., 2015). Functional analyses indicated that WLS physically interact with WNTs and escort WNTs from ER, Golgi to plasma membrane where they hand off WNTs to their extracellular carriers such as secreted FZD-related proteins and WNT inhibitory factor 1 (Das, Yu et al., 2012, de Almeida Magalhaes, Liu et al., 2024, Wolf & Boutros, 2023). Study of human brain organoids indicated that spatially restricted induction of WLS marks the earliest emergence of non-telencephalic brain regions, and study with Wnt1-Cre-mediated WLS knockout showed that WLS plays a role in the generation of ChPe in mice but complete WLS knockout caused axis formation defects and early death of the affected animals before ChP formation (Carpenter et al., 2010, Fu, Jiang et al., 2009, Jain, Gut et al., 2025). In zebrafish, maternal deposited *wls* transcripts and Wls proteins can help null mutants to bypass the early axis developmental defects and further studies indicate that Wls modulates the development of HA by controlling the expression of *neurog1* in proneurons in the developing diencephalon (Kuan et al., 2015, Wu et al., 2015). Wls also has been shown to interact with Fgf signaling to modulate the development of jaw cartilages through fine-turning the expression gradient of *fgf3* transcripts (Wu et al., 2015). However, whether Wls involves in the development of zebrafish ChP and how Wls modulates the development of HA neurons and their progenitors are still unknown.

Here through three-dimensional (3D) gene expression analyses we report that the formations of cholinergic, substance P-ergic and glutamatergic HA neurons at 96 hpf (hours post-fertilization) are differentially reduced in *wls* mutants. Conversely, *clusterin* (*clu*) gene expression domains in the ChPe are expanded in *wls* mutants at 72 and 96 hpf, suggesting that loss-of-Wls caused an expansion of gliocyte-lineages in diencephalic ChP. We also found that the reduction of *neurog1* signals in *wls* mutants is caused by failed accumulation of *neurog1* transcripts at 48 hpf and that some diencephalic *neurog1*-positive cells are HA neuronal progenitors. Interestingly, 3D gene expression analyses showed that while *neurog1* expressions are reduced, the expressions of Notch effector *her6* are increased and expanded into HA progenitor zones in *wls* mutants at 48 hpf. Ectopically over-expressing Her6 in *neurog1*-positive cells caused partial reduction of *neurog1* expressions in HA progenitor zones. These results indicate that Wls restricts the expressions of Notch effector *her6* to promote the specification of *neurog1* proneurons but Wls demotes the generation of ChPe during the development of embryonic dorsal diencephalon.

## Results

### Habenular neuronal fates are changed in *wls* mutants

Prior study indicated that loss of Wls activity resulted in complete loss of ventral habenular neurons because the expressions of two HA markers, the *vachtb* (*slc18a3b*) and *ano2* were either greatly (*vachtb*) or completely (*ano2*) gone from the chromogenic ISH staining experiments (Kuan et al., 2015). However, it is unclear whether each neuronal lineage showed equal reduction rate in the absence of Wls. To better demonstrate the spatial distribution of different neuronal lineages and the spectrum of gene expression changes in HA of wild-type (WT, including *wls*/+) or *wls* larvae, double fluorescent mRNA in situ hybridization (FISH) stainings were performed with larvae staged at 96 hours post-fertilization (hpf) (Wang, Pan et al., 2021). Three antisense probes of *vachtb* (*slc18a3b*), *tac3a* or *vglut2a* (*slc17a6b*) genes were adopted to mark the cholinergic, substance-Pergic or glutamatergic HA neurons, respectively (Biran, Palevitch et al., 2012, Hong, Santhakumar et al., 2013, Thisse, 2005). Another antisense probe of *cpd2* gene was adopted to mark the majority of HA cells, and to align all the confocal Z-stacks acquired from the WT or *wls* mutant samples (Wang et al., 2021). The representative Z-projection images from double FISH of *vachtb* + *cpd2*, *tac3a* + *cpd2*, or *vglut2* + *cpd2* were showed in Figure 1B to 1G’. As can be seen, the staining of all four HA markers were apparently reduced at different levels in the *wls* mutants (Figure 1E-G’). Two three-dimensional (3D) four-gene expression maps were generated to show the aligned and averaged gene expression domains of *vachtb* (*slc18a3b*), *tac3a*, *vglut2* and *cpd2* in the HA of WT (32 Z-stacks aligned) or *wls* mutants (24 Z-stacks aligned)(Figure 1H to S’, Supplemental Figure 1 and Supplemental Table 1A-B). In Figure 1H to S’, the differential expression changes of each HA markers or their overlapped regions in *wls* mutants are clearly shown along both the X-Y (1H to 1O) and X-Z (1H’ to 1O’) axes. Quantification analyses of each gene expression volumes (vol., in µm^3^) indicated that while the averaged expressions of *cpd2* in both LHA and RHA in *wls* mutants remained around 50% (48.32% and 50.07%) to their WT siblings, the expressions of *vachtb* (*slc18a3b*), *tac3a* or *vglut2* each in LHA and RHA in *wls* mutants remained 32.15% and 22.36%; 4.66% and 27.08% or 18.22% and 24.4% respectively to their WT siblings (Figure 1T to W, Supplemental Table 1C-E). These discoveries suggest that there are different levels of cell-lineage alternations of each these three types of HA neurons. In addition, while the left-right (L-R) ratio of averaged *cpd2* expressions in *wls* samples was highly similar to their WT siblings (WT or *wls* =124.6% or 120.2%), there are apparent changes of L-R ratios for the expressions of *vachtb* (WT or *wls*= 37.0% or 53.2%), *tac3a* (WT or *wls* = 18.5% or 3.2 %) and *vglut2* (WT or *wls* =159.9% or 119.5%)(Supplemental Table 1F). These results also demonstrated that 3D gene expression analyses can reveal differences of L-R gene expression ratios between WT and *wls* mutants that were overlooked by prior report utilizing chromogenic gene expression analyses (Kuan et al., 2015). The changes of HA neuronal cell fates in *wls* mutants suggested that either cell proliferation rate or cell-fate decision, or both factors are affected upon loss of Wls activity. However, whether cell fates in other diencephalic domains are changed has never been examined before.

#### Expression domains of *clu* in choroid plexus are expanded in *wls* mutants

Because complete WLS knockout caused early death of the affected animals before ChP formation (Carpenter et al., 2010, Fu et al., 2009, Jain et al., 2025), we examined whether loss-of-Wls in zebrafish affects the development of diencephalic ChP (dChP). Utilizing *clusterin* (*clu*) trancripts to mark ChP epithelila (ChPe) at 72 and 96 hpf (Jiao, Dai et al., 2011), FISH staining were performed along with *gng8*, another HA marker, to distinguish the *wls* mutants from their WT siblings (Kuan et al., 2015). Our results demonstrated that the expressions of *clu* were increased in both the diencephalic and hindbrain ChPs (dChP and hChP) (Figure 2A-F). Quantification of *clu* and *gng8* expressions in HA of WT and *wls* mutant samples indicated that the increases of *clu* and the decreases of *gng8* expressions in *wls* mutants at 72 and 96 hpf are statistically significant (Figure 2G-H, Supplemental Table 2A-D). Apparent increases of *clu* expressions in the dChP at both 72 and 96 hpf in the *wls* mutants are better demonstrated in the 3D virtual maps generated using *gng8* signals for image alignment. (Figure 2I-L, Supplemental Figure 2 and Supplemental Table 2E-H). These results are very interesting because they showed a kind of ChP developmental defects that is opposite to those above-mentioned defects resulted from individual Wnt signaling knockout or gain-of-function animals (Carpenter et al., 2010, Kaiser et al., 2021, Langford et al., 2020, Parichha et al., 2022). These results also suggest that Wls may play a role in promoting neuronal cell fate specification because specification of ChP epithelia from neuroepithelial cells seems to require the repression of neural cell fate (Lun et al., 2015).

**Figure 2.**
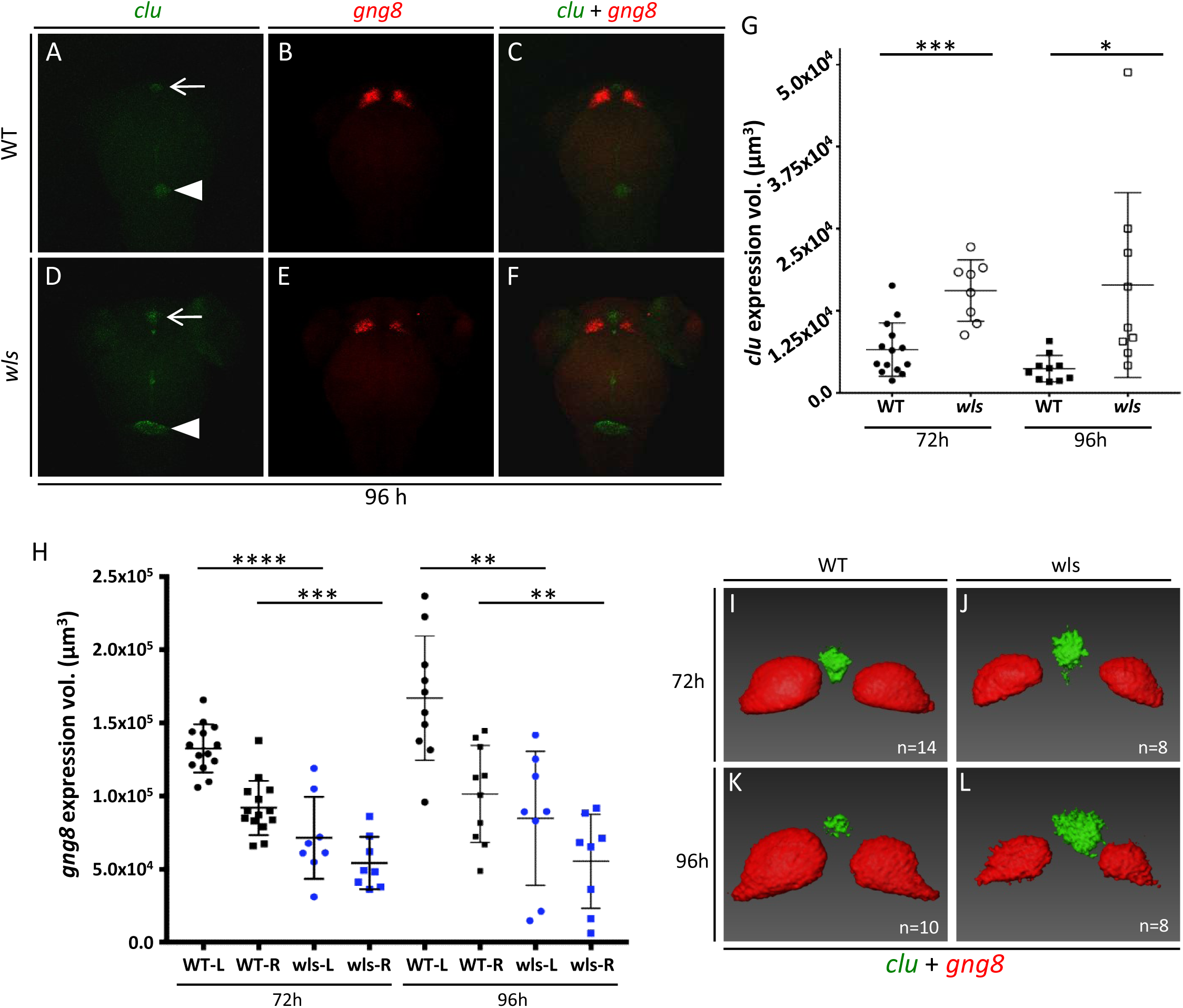
Expression domains of *clu* in choroid plexus are expanded in *wls* mutants. (A-F) Representative images showing *clu* (green, labeling choroid plexus) or *gng8* (red, labeling HA) expressions in WT (A-C) or *wls* mutants (D-F) at 96 hours post-fertilization (h). ChP = choroid plexus, arrows indicate the 3^rd^ ventricle ChP, arrow heads indicate the 4^th^ ventricle ChP. (G-H) Statistical charts of the *clu* (G) or *gng8* (H) expression volumes (vol.) of WT or *wls* mutant samples at 72h or 96h. Confidence *p* value (*p*) is denoted by “*”. “****” means P< 0.0001; “***” means P< 0.001; “**” means P< 0.01 and “*” means P<0.05 in t-test. (I-L) Merged virtual 3D expression maps of *clu* and *gng8* from samples of 72h (I-J) and 96h (K-L). (See Supplemental Figure 3 and Supplemental Table 2 for details)

#### Diencephalic *neurog1-*positive neural progenitors developed into HA neurons

Prior studies reported that neuronal progenitors located ventral to the developing pineal contribute to HA neurons, *neurog1* expressing in the HA progenitor zones in *wls* mutants is decreased, and combined loss of *neurog1* and *neurod4* caused HA neurogenesis defects (Concha, Russell et al., 2003, Halluin, Madelaine et al., 2016, Kuan et al., 2015). However, whether reduction of *neurog1* expression in HA progenitor zone in *wls* mutants is reflecting a decrease of *neurog1* transcripts or a decrease of *neurog1*-expressing proneurons and whether HA neurons can derive from the *neurog1*-positive proneurons has never been experimentally demonstrated before. To verify the location and the extent of *neurog1* expression reduction in HA progenitor zones in *wls* mutant embryos, FISH staining utilizing mixed *neurog1* plus *fgf8* (distinguish WT or *wls* mutants) antisense probes and anti-EGFP (for labeling pineal complex) antibody labeling were performed, then double-colored confocal Z-stacks were collected. Consistent with the previous results, the expressions of *neurog1* in both left and right sub-ventricular zones (SVZ, the HA progenitor zones) were reduced apparently (Figure 3A and 3D, Supplemental Figure 4A-F). The reductions of *neurog1* expressions in left and right HA progenitor zones in *wls* embryos are better visualized in the virtual 3D maps, while the averaged *neurog1* expressions in the middle domain along the midline were highly similar between the WT and *wls* samples (Figure 3B-C, 3E-F, Supplemental Figure 3). Quantification of *neurog1* expressions in the left and right HA progenitor zones indicated that the differences between WT siblings and the *wls* mutants are statistically significant (Figure 3G, Supplemental Table 3C-D). Because the development of pineal complex (anti-EGFP labeled) in *wls* is comparable to that in their WT siblings (Kuan et al., 2015), we aligned and merged all the Z-stacks of WT (21 stacks) and *wls* (14 stacks) embryos using EGFP channel (35 Z-stacks) to generate one virtual 3D map for visualizing the averaged *neurog1* expressions simultaneously (Figure 3H-I). Apparent differences and locations of *neurog1* expressions in the HA progenitor zones are clearly seen in this combined 3D model as well. In addition, we introduced *neurog1*:*egfp* transgene into the mutant carriers (*wls*/+ males and females) and found that the EGFP expressions driven by the endogenous *neurog1* promoter were no different in the WT siblings and *wls* mutants at 24 and 48 hpf (Supplemental Figure 4G-R). Therefore, the reduction of *neurog1* signals reflects a decrease of *neurog1* transcripts in the HA progenitor zone. Because *neurog1*:*egfp* generated EGFP signals are present in HA at 72 and 96 hpf despite that the endogenous *neurog1* mRNAs were not detectable in HA at all (Figure 3J-K, 3L-M), it suggested that the EGFP-positive HA neurons may derive from the *neurog1*-positive progenitors. To confirm this hypothesis, we adopted two cell-labeling techniques, the photo-convertible fluorescent protein Kaede and the infrared laser-evoked gene operator (IR-LEGO), to trace the developmental destination of *neurog1*-positive cells in the HA progenitor zones (Figure 3O) (Ando, Hama et al., 2002, Deguchi, Itoh et al., 2009, Kamei, Suzuki et al., 2009, Shimada, Kawanishi et al., 2013). Kaede-mediated cell-tracing demonstrated that some of the *neurog1*-positive cells in the HA progenitor zones indeed became HA neurons (Supplemental Figure 4S-X), and IR-LEGO-mediated cell-tracing demonstrated that some of the *neurog1*-positive cells in the HA progenitor zones would became HA neurons (Figure 3P-S). All these results indicate that Wls activities are required to specify the *neurog1*-expressing cells rather than to produce the cells which express *neurog1* in the HA progenitor zones, and dorsal diencephalic *neurog1-*positive proneurons are HA neuronal progenitors.

**Figure 3.**
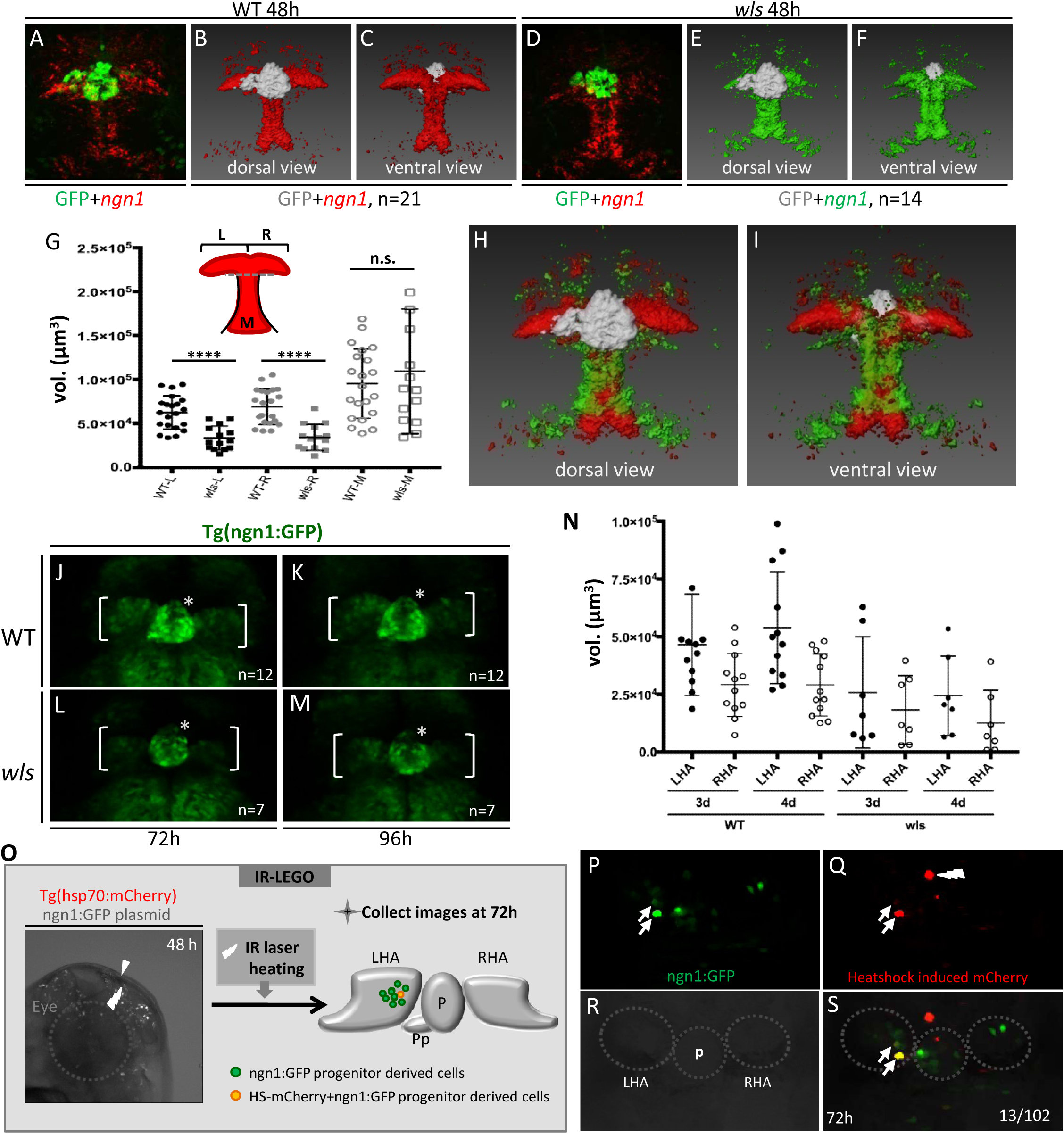
Diencephalic *ngn1-*positive progenitors developed into HA neurons. (A-I) Images showing *ngn1*-expressing cells labeled by antisense *ngn1* mRNA probes (in red) in WT (A) or *wls* (D) embryos at 48h. *foxD3*:GFP signal is utilized to locate the pineal and parapineal (in green). Aligned and averaged virtual 3D *ngn1* expression maps for WT (B, C) or *wls* (E, F) embryos. (G) Statistical chart for the volumes (vol.) of *ngn1* expression domains in left (L), right (R) or middle (M) diencephalic regions. Confidence *p* value (*p*) is denoted by “*”. “****” means P< 0.0001, “n.s.” means no statistical difference in t-test. (H-I) Merged virtual 3D *ngn1* expression maps from B and E (H) or C and F (I). (J-N) Images showing *ngn1* -positive cells (promoter-driven GFP) contribute to HA neurons at 72 (J, L) or 96 hpf (K, M) in the WT (J-K) or *wls* mutant (L-M) embryos. “*” indicates pineal; brackets indicate the left and right HA. (N) Statistical chart for the vol. of GFP-positive cells in LHA or RHA. (O) Diagram showing how IR-LEGO (Infrared-Laser Evoked Gene Over-expression) was conducted to label *ngn1*-positive progenitors in dorsal diencephalon at 48h. “P” indicates pineal; “Pp” indicates parapineal. Lighting sign indicates the laser entry site. Dotted circle indicates the left eye. (P-S) Images showing *ngn1*-expressing cells (P) labeled by heatshock-induced mCherry (Q) developed into HA neurons at 72 h (S). Dotted circles indicate the pineal (P), LHA or RHA. (Also see Supplementary Figure 4, 5 and Table 3 and 4)

#### Loss-of-Wls caused expansion of *her6* expression domains in dorsal diencephalon

Because the neuronal fates were changed in the remaining HA in *wls* mutants at 96 hpf (Figure 1) and the cells that should express *neurog1* in the HA progenitor zone were still present in the *wls* mutant embryos at 48 hpf despite *neurog1* transcripts were low or absent in those cells (Supplemental Figure 4G-R), these suggest that those cells might adopt different cell fates in the *wls* mutants. Previous studies indicated that *neurog1* plays roles downstream to *flh*, *her6* or *pax6a* in the neurogenesis of pineal, thalamus or HA, respectively during zebrafish embryonic brain development (Cau & Wilson, 2003, Halluin et al., 2016, Scholpp, Delogu et al., 2009). Because the expressions of *flh* and *pax6a* were shown to be normal in *wls* mutants and the ChP developmental defect in *wls* mutants resembles the defect in Notch signaling perturbed larvae, we decided to examine the expression of *her6* in *wls* mutants (Bill et al., 2008, Kuan et al., 2015). The results showed that *her6* expressions were markedly increased in *wls* mutants at 48 hpf, suggesting it plays a role in Wls-mediated specification of *neurog1* progenitors or a role in ChPe development. (Figure 4A-F, Supplemental Figure 5A-H and Supplemental Table 5A-B). Quantification of *her6* expression domains in the left and right hemispheres demonstrated that the differences between WT siblings and *wls* mutants are statistically significant (Figure 4G-I, Supplemental Table 5C-D). The positional relationships between the *neurog1* and *her6* expression domains were clearly demonstrated in the virtual 3D maps in Figure 4J-M, in which the *neurog1* and *her6* are expressed in adjacent but separate domains in the developing dorsal diencephalon (Figure 4J-K). In comparison, in the virtual 3D map combing image stacks from both WT and mutant samples, the *her6* expression domains are larger and expanded into the normal *neurog1* expression domains in the left and right hemispheres in *wls* mutants (Figure 4L-M). Because prior report indicated that presence or absence of *her6* promote Ascl1-mediated GABAergic or *neurog1*-mediated glutamatergic neuronal generation respectively during thalamus development (Scholpp et al., 2009), our observations therefore strongly suggest a model that Wls is required either to promote *neurog1*-mediated *her6* expression suppression or to de-repress *her6*-mediated *neurog1* expression suppression during the development of HA neurons.

**Figure 4.**
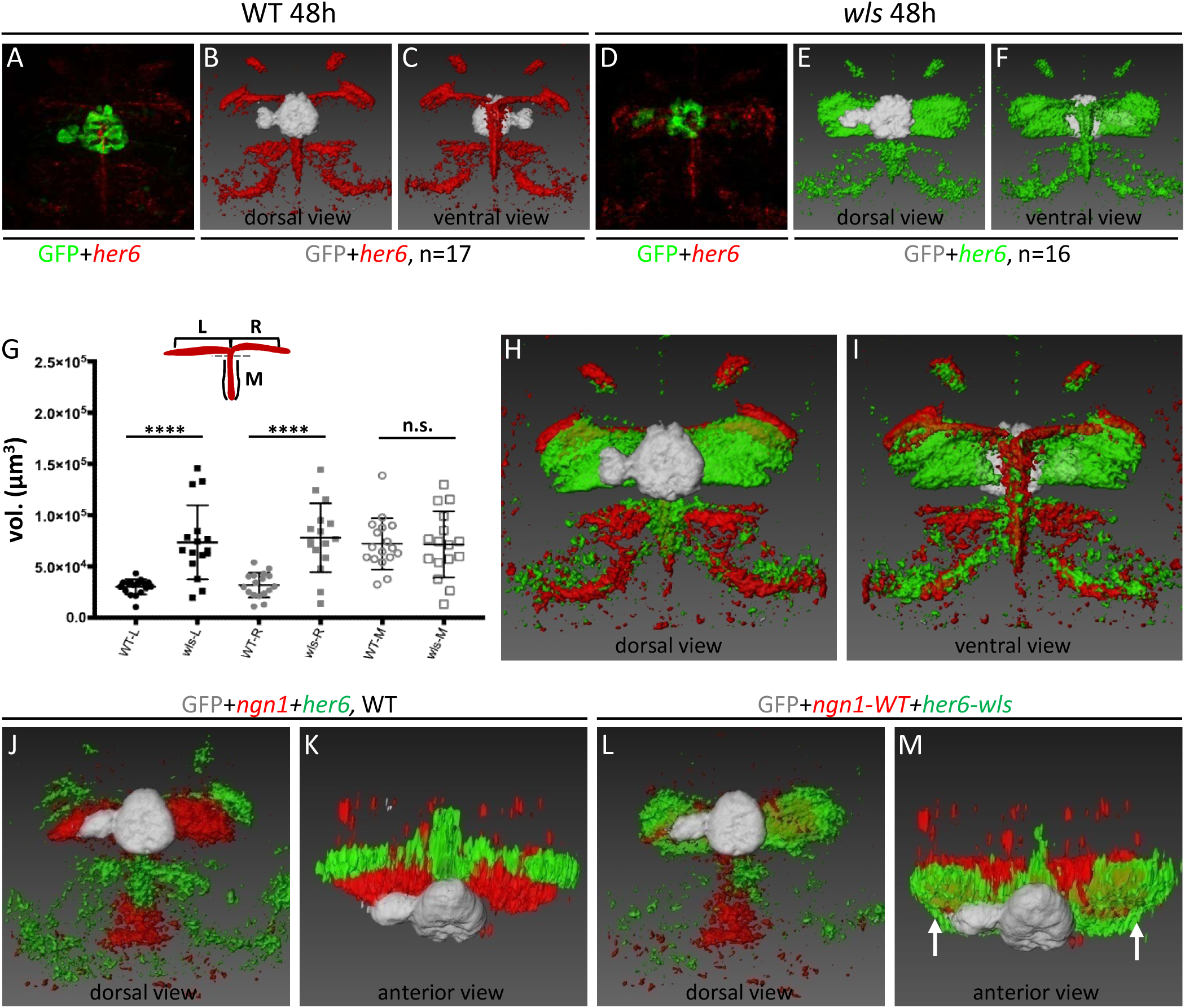
Loss-of-Wls causes expansion of *her6* expression domains in dorsal diencephalon. (A-F) Images showing *her6*-expressing cells labeled by antisense *her6* mRNA probes in WT (A) or *wls* (D) embryos at 48h. *foxD3*:GFP transgenic line was utilized to locate the pineal and parapineal in each sample. Aligned and averaged virtual 3D *ngn1* expression maps (B, C, E, F) for WT (B-C) or *wls* (E-F) embryos were generated for samples from 48h. (G) Statistical chart for the volumes (vol.) of *her6* expression domains in left (L), right (R) or middle (M) diencephalic regions (see the insert drawing). (H-I) Merged virtual 3D *her6* expression maps from B and E (H) or C and F (I). (J-M) Merged virtual 3D *ngn1* plus *her6* expression maps aligned using Pineal (GFP positive) of 48h embryos from WT groups (J-K) or from WT (for ngn1 expression) plus *wls* (for *her6* expression) groups. Arrows in (M) indicate the expansion of *her6* expression domains into the HA progenitor zones. (See Supplemental Figure 6 and Supplemental Table 5, 6 for details)

#### Over-expressing *her6* suppresses *neurog1* expressions in the HA progenitor zones

In order to understand the regulatory relationships between *neurog1* and *her6*, we generated *neurog1*:*her6* plasmid which can express full-length *her6* mRNA ectopically under the control of *neurog1* promoter (Figure 5A). Transiently inducing *her6* expressions in the *neurog1* expressing cells through plasmid injections caused apparent reduction of *neurog1* expressions in the HA progenitor zones (Figure 5B-C, E-J, Supplemental Figure 6). Quantification of *neurog1* expression volumes in control and *neurog1*:*her6* plasmid-injected samples demonstrated that the differences in left and right HA progenitor zones are statistically significant whereas the *neurog1* expressions in the middle domains are not significant (Figure 5D, Supplemental Table 7). This result strongly supports our notion that Wls-mediated suppression of *her6* expression plays a role in specifying some of the *neurog1*-positive neurons during the development of HA. Because there is no increase of apoptosis (Kuan et al., 2015) and there are incorrect cell fate specifications in the HA progenitor zones in the *wls* mutants (Figure 1, 3, 4), those indicate that the neuronal generation and specification defects in HA at 96 hpf (Figure 1) are resulted from incorrect cell specification at earlier developmental stages and slightly reduced cell proliferation (Figure 3 and 4, Supplemental Figure 7). This notion is consistent with the previous statement that, as a protein controlling the secretions of diverse Wnt molecules, Wls plays distinct roles in regulating the generation and specification of HA neurons and its laterality (Kuan et al., 2015).

**Figure 5.**
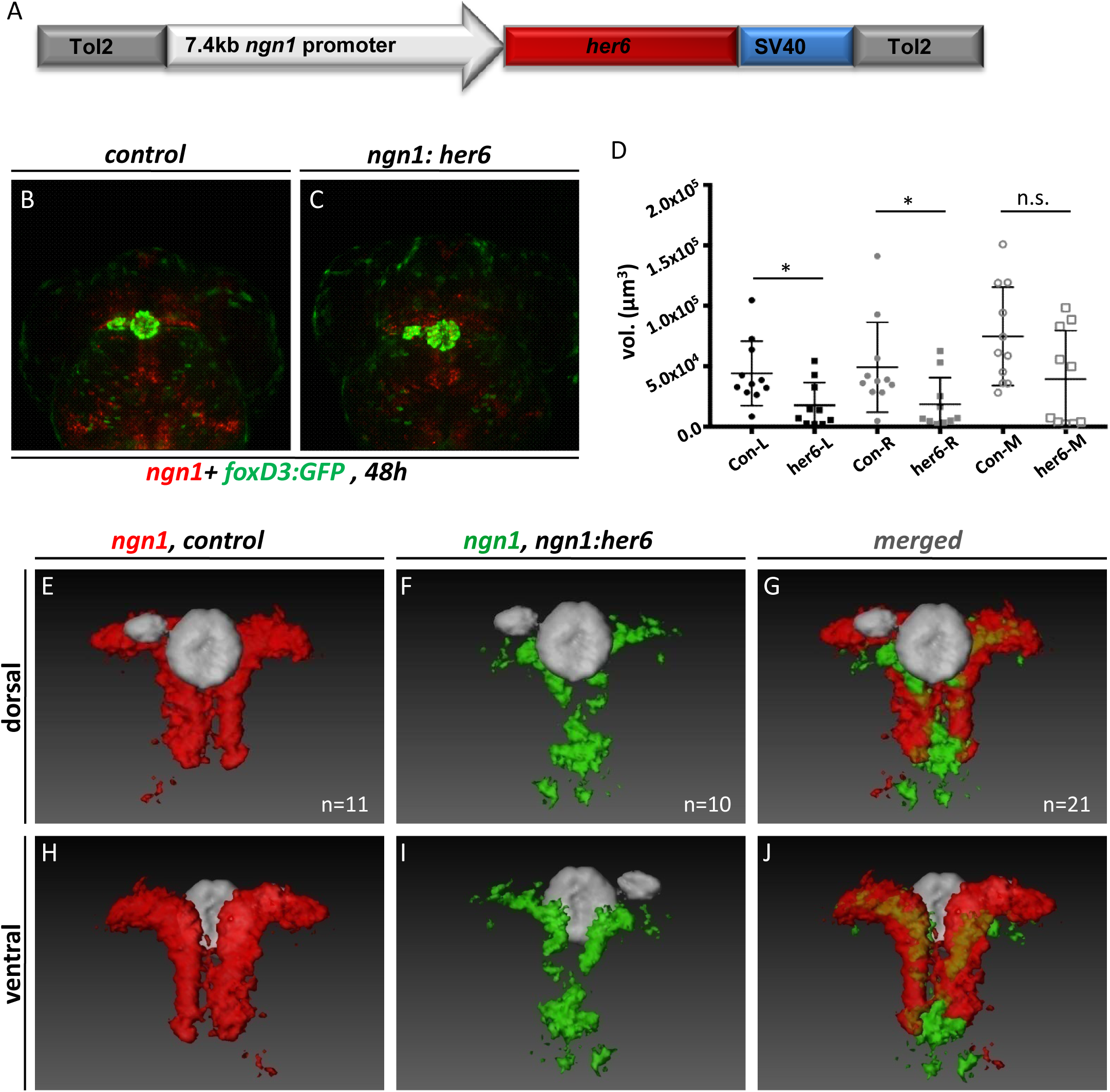
Over-expressing *her6* suppresses *ngn1* expressions in HA progenitor zones. (A) Schematic drawing of the *ngn1* promoter-controlled *her6*-expressing plasmid. (B-C) Images showing *ngn1*-expressing cells labeled by antisense *ngn1* mRNA probes in WT (B) or *wls* (C) embryos at 48h. *foxD3*:GFP transgenic line was utilized to locate the pineal and parapineal (in grey) in each sample. (D) Statistical chart for the volumes (vol.) of *ngn1* expression domains in left (L), right (R) or middle (M) diencephalic region. Confidence *p* value (*p*) is denoted by “*”. “*” means P< 0.05, “n.s.” means no statistical difference in t-test. (E-J) Images showing averaged expression vol. of *ngn1* in samples from injection control group (E, H) or *ngn1*:*her6* plasmid-injected group (F, I). Merged images (G-J) from (E-F) and (H-I) utilizing pineal-signal-guided image alignment. (See Supplemental Figure 7 and Supplemental Table 7 for details)

## Discussion

In this report, we have unraveled the regulatory relationships between *wls*, *her6* and *neurog1* that have never been demonstrated before during the generation and specification of HA neurons in developing vertebrate diencephalon. In addition, we found that the interaction between Wls and Notch signaling guides the development of HA neurons and ChP gliocytes in opposite directions. Utilizing FISH and 3D gene expression analyses, we found that loss of Wls activity caused uneven reductions of cholinergic, substance-Pergic or glutamatergic neurons in the developing HA, despite that an overall reduction of neurons is 50% in both left and right HA (Figure 1T, 1U and 1V). Our observation that a steady and continuous cell proliferation reduction exists in the *wls* mutant embryos (Supplemental Figure 7) may explain that the evenly reduced generation of *cpd2* neurons in the L-R HA is resulted from reduced cell proliferation. Nonetheless, the uneven reductions of different types of neurons in the HA of *wls* mutants suggest that cell fate changes during the development of HA progenitors may account for the alterations of neuronal ratios when Wls activities are depleted. These cell fate alterations are apparently not caused by abnormal cell migration because none of the four tested HA neuronal markers showed any relocation outside the main *cpd2* expression domains. This conclusion is consistent to what was reported before using the other HA markers such as *f-spondin*, *nrp1a*, *lov* and *ron* (Kuan et al., 2015). In addition, through introducing *neurog1*:*egfp* transgene into *wls* mutants we are able to clarify that reduced ISH signals from *neurog1* mRNA staining in the *wls* mutants is caused by decreased accumulation of *neurog1* mRNAs, indicating that Wls activity is required to promote the expression of *neurog1* in the HA progenitor zone. This is important for our subsequent experiments which proofed the first time that some *neurog1*-positive cells in the HA progenitor zones indeed are HA neuronal progenitors (Figure 3O-S and Supplemental Figure 4S-X).

One very interesting observation is that we found the sizes of both diencephalic and hindbrain ChPs are significantly increased in the *wls* mutants (Figure 2). Because this ChP phenotype resembles to the developmental defects seen in the Notch signaling-perturbed larvae (Bill et al., 2008, Hunter & Dymecki, 2007), we hypothesized that loss-of-Wls might perturb the Notch signaling activities which led to ChP size increase in 72 and 96 hpf *wls* mutants. Subsequently we found another interesting discovery that the expressions of *her6*, the Notch-dependant transcription factor, were apparently increased in the dorsal diencephalon of *wls* mutants at 48 hpf (Figure 4). This result may reflect the cause of the ChP size increases seen in the Notch signaling-perturbed embryos (Bill et al., 2008, Hunter & Dymecki, 2007). Because either increasing (*dld* or *dla* morphants, or Notch1 constitutively activation) or reducing (DAPT-treatment, *notch1b* morphants) Notch signaling activities could cause size increases of ChP, it suggests that fine-tuning Notch receptor activities by antagonizing each other may play a key role in ChP development. Therefore, deciphering the relationships between the specificity of each Notch component on *her6* expressions and the role of *her6* expression on ChP development will be the keys to reveal the molecular mechanisms of how Wls interacts with Notch signaling component to simultaneously control the development of HA neurons and glial-derived ChP epithelial cells. Because ectopically expressing Her6 can suppress the expressions of *neurog1* in the HA progenitors (Figure 5), this indicates that Wls-mediated suppression of *her6* expression is required for specifying the *neurog1*-positive progenitors of HA neurons and may be required to down-regulate the generation of glial cells. In addition, because previous studies indicated that the expressions of *pax6a* were normal in the *wls* mutants and *neurog1* plays downstream to *pax6a* in promoting the neurogenesis of HA (Halluin et al., 2016, Kuan et al., 2015), suggesting that *pax6a*-mediated HA development may account for the remaining *neurog1*-positive HA progenitors and neurons seen in the *wls* mutants (see our model below).

In order to sum up a model of how *wls*, *her6* and *neurog1* interact with each others during the development of HA and ChP in the embryonic dorsal diencephalon, we evaluated their expression positional relationships in 3D at 48 hpf. Using the pineal organ (P, marked by EGFP) as the image aligning landmark from three WT samples each also labeled with either *her6*, *neurog1* or *wls* antisense mRNA probes (Figure 6A-C), we found that *wls* expressing domains are situated ventral to the *her6* expressing domains near the ventricular zone where most of the neural stem cells are generated. And, the *her6* expressing domains are situated ventral to the *neurog1* expressing domains where the HA progenitors are generated. The spatial relationships between *wls*, *her6*, *neurog1* and HA along with the fact that *pax6a* expression is not affected by loss-of-Wls all pointed out the existences of Wls-dependant and Wls-independent formation of *neurog1*-positive HA progenitors that later on may rely on Wls activities to proliferate and differentiate into mature HA neurons (Kuan et al., 2015). Because loss of Wls caused an expression elevation of Notch signaling component *her6* in the developing dorsal diencephalon, a model therefore is emerged that Wls-mediated Wnt secretions interact with Notch signaling to govern the proper generation of ChPe cells and HA neurons simultaneously, suggesting that Wls plays a key and early role in neuronal-glial cell fate determination during the development of both ChPe and HA. This model of us well echoes the facts that the generation of ChPe from neuroepithelial cells requires the repression of neural cell fate and misexpression of Hes protein family member significantly increased the population of glial cells at the expense of neurons [9, 51]. Additionally, because some Wnt signaling molecules such as Wnt1, WNT5A and beta-catenin are playing positive roles in the ChP development in mice (Kaiser et al., 2021, Langford et al., 2020, Parichha et al., 2022), it suggests that Wls-dependant Wnt secretions exist to either repress the elevation of Notch signaling directly or indirectly by repressing the other signaling (another Wnt?) which normally keeps the Notch signaling in check.

**Figure 6.**
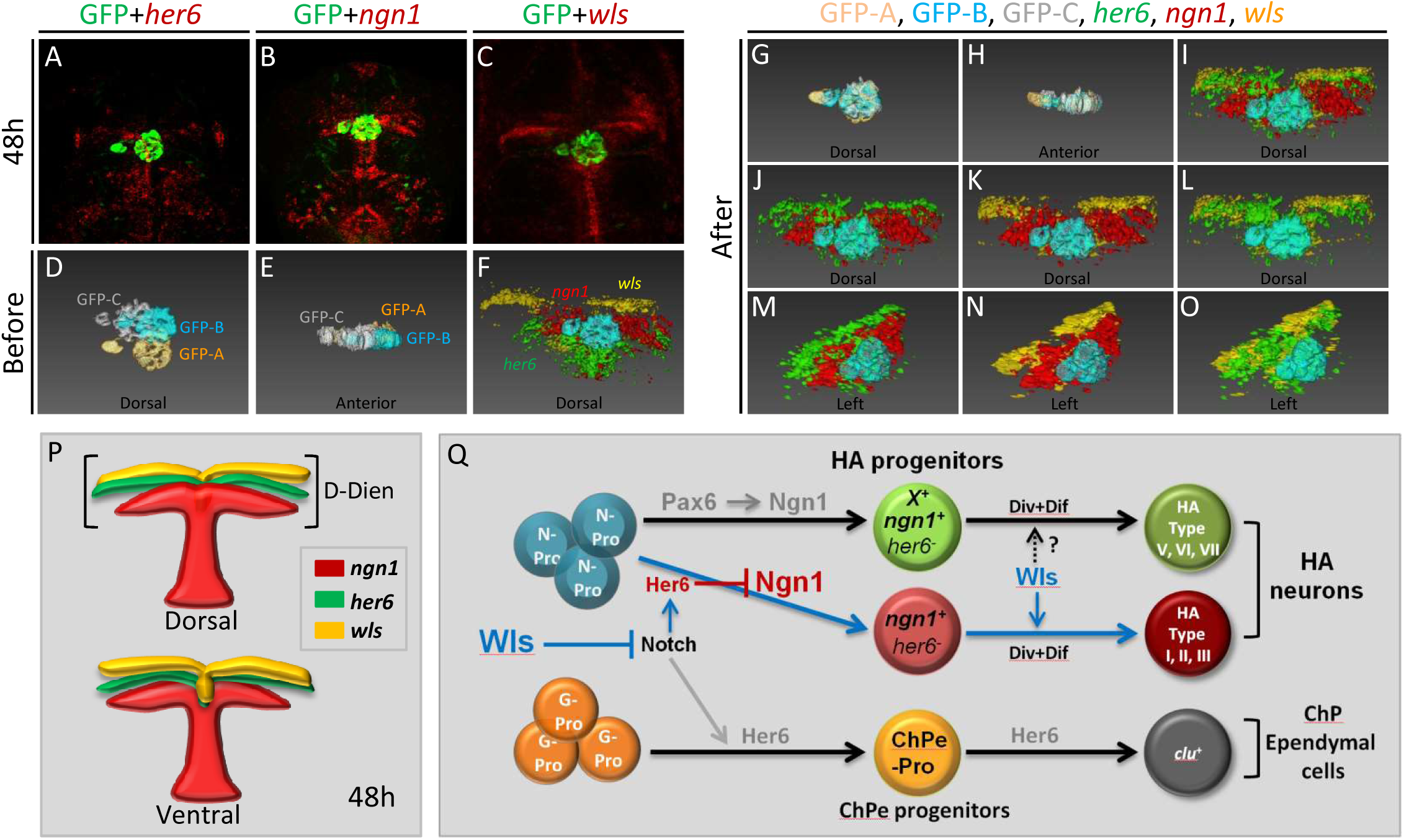
Wls interacts with Notch signaling to control early neural-glial cell fates and promotes the specification and proliferation of HA progenitors. (A-C) Representative Z-projection snapshots showing the expressions of *her6* (A), *ngn1* (B) and *wls* (C) transcripts in the Tg(Foxd3:GFP) embryos at 48 hours post-fertilization (48h). Anti-GFP labeling indicated the pineal and parapineal in these samples. Dorsal means dorsal view, ventral means ventral view. (D-F) Snapshots generated in Amira software showing fragmented and thresholded isosurface views for GFP (in D, E) or GFP plus *her6*, *ngn1* and *wls* signals (in F) from the image stacks of (A), (B) and (C), before image alignment. GFP-A, GFP-B, GFP-C represent the pineal and parapineal from figure 6A, 6B and 6C, respectively. (G-O) Snapshots generated in Amira software showing fragmented and thresholded surface views for images of (A), (B) and (C), after image alignment. (P) Models of the expression relationship between *her6*, *ngn1* and *wls* in the dorsal diencephalon (D-Dien) near the third ventricle in zebrafish embryonic brain at 48h. (Q) Summary of our model for the roles of Wls and Notch on early neural or glial cell fate determination and how Wls guides the specification and proliferation of HA progenitors during zebrafish embryonic brain development.

### Conclusions

Our results have revealed a novel regulatory relationship between Wls and Notch signaling on balancing the generation of neurons and gliocytes during the development of HA and ChPe in dorsal diencephalon. This well echoes to the recent discovery that a spatially restricted induction of Wls denotes the earliest differentiation of non-telencephalic cells in human brain organoids (Jain et al., 2025). Because the early developmental defects seen in mouse mutants have hindered functional studies of Wls after axis-formation (Carpenter et al., 2010, Fu et al., 2009, Wu et al., 2015), our report therefore has demonstrated that Zebrafish loss-of-Wls embryos or larvae carrying maternally deposited *wls* proteins and transcripts are excellent conditional *wls* mutants to decipher the function of Wls on the development of vertebrate brain or the pathogenesis of brain defects.

## Methods

### Fish Maintenance and Sample Collection

AB (wildtype) strain zebrafish were obtained from Taiwan Zebrafish Core Facility (TZCF) in Academic Sinica. The lines of *wls*/+ (*fh252* allele) and Tg(*foxD3*: *GFP*) were obtained from Drs. C.B. Moens (Fred Hutch Cancer Center, Seattle, WA, U.S.A.) and M. E. Halpern (Carnegie Inst. for Science, Baltimore, MD, U.S.A.) (Barth, Miklosi et al., 2005, Kuan et al., 2015). ZFIN fish IDs for AB, *wls*/+ and Tg(*foxD3*:*GFP*) are ZDB-GENO-960809-7, ZDB-ALT-071019-5 and ZDB-TGCONSTRCT-070117-95, respectively. The two fish lines of Tg(*neurog1*:*egfp*) and Tg(*hsp70*:*mCherry*-*CMV*:*gfp*-*nanos*3’UTR) were generated based on the articles by Blader et al. and Evans et al., 2005 (Blader, Lam et al., 2004, Evans, Yamamoto et al., 2005). All fishes were maintained and bred according to standard procedures described on ZFIN (http://zfin.org). All larva were collected from natural spawning and reared/staged in 0.003% PTU (Sigma P7629). Except for those utilized in live images, staged embryos and larva will be fixed at 36, 48, 72, 84 or 96 hours post-fertilization (hpf) in 4% paraformaldehyde (PFA, Sigma P6148) in PBS overnight at 4°C then stored in 100% MeOH (Merck 1070182511) at −20°C until processed for RNA *in situ* hybridization or immunohistochemical (IHC) staining. All animal procedures were performed in accordance with the protocol approved by the Institutional Animal Care and Use Committee (IACUC) at Academia Sinica and National Taiwan University (Protocol #20-12-1618 and #113-00145).

#### Transgene constructions and DNA microinjection

The *neurog1*:EGFP transgenic plasmid was generated based on the article by Blader et al. (Blader et al., 2004). The gene name “*neurog1*” was previously called “*ngn1*” in the article by Blader et al.. Basically we utilized a modified Tol2 vector (obtained from Dr. Koichi Kawakami, NIG, Shizuoka, Japan) to generate the final plasmid containing a 7.4kb upstream fragment of *neurog1* gene locus 5’ to the *egfp*-SV40 expression cassette inside the Tol2 arms (primer pair A A C T C G A G A C T A G T T T T T G G A C T G T T A T C a n d A A G G A T C C T G T T G ATAACCTGGAGAAATT) (Kawakami, Takeda et al., 2004, Urasaki, Morvan et al., 2006). Tg(*hsp70*:*mCherry*-CMV : *GFP*-*nanos*3’UTR) transgenic construct was generated by inserting P C R a m p l i f i e d 1 . 5 K b *h s p 7 0* p r o m o t e r f r a g m e n t (p r i m e r p a i r T T G T C G A C A C G T G A A T G G T T T G T G A G C A C a n d CCGGATCCGATTGATTTCAAGAAACTGCA) (Evans et al., 2005), *mCherry* fragment (p r i m e r p a i r A C G G A T C C A A T G G T G A G C A A G G G C G A G a n d T T A T C G A TCCCGGGTTACTTGTACAGCTCGTCCATG) and enzyme digested CMV:*GFPnaos*3’UTR fragment (SalI at 5’ and XbaI at the 3’ ends) into the Tol2 vector. The pME-*mCherry* and pCS2-CMV:*egfp-naos3’UTR* plasmids were obtained from Taiwan Zebrafish Core Facility (TZCF) and Dr. Marnie Halpern (Embryology Dept., Carnegie Institute for Science, Baltimore, USA), respectively. The *neurog1*:*Kaede* transgenic construct was generated by replacing the *egfp* of *neurog1*:*egfp* with PCR amplified DNA fragment of pKaede-S1 plasmid (MBL international corporation, Woburn, MA, USA). The *neurog1*:*her6* transgenic construct was generated by replacing the *egfp* fragment with PCR amplified DNA fragment of *her6* ORF plus its 3’UTR (ZFIN: ZDB-TSCRIPT-090929-15349) from cDNAs generated from 48 hpf total RNAs. DNA microinjections were performed as described before (Kawakami et al., 2004, Urasaki et al., 2006). In this report, we injected 25-40pg of plasmids plus 25pg of transposase mRNAs (generated from pCS-TP obtained from Dr. Koichi Kawakami, NIG, Shizuoka, Japan) into each embryos at 1-2 cell stages.

#### Fluorescent mRNA *in situ* hybridization and immunohistochemical staining

Digoxigenin (DIG, Roche 11277073910)-labeled *cpd2, gng8, neurog1*, *her6*, *fgf3* or *wls* probes, or fluorescein (FLU, Roche 11685619910)-labeled *vachtb* (*slc18a3b*), *tac3a, vglut2a* or *clu* RNA probes were generated according to previously described procedures (Wang et al., 2021). We introduced *fgf3* antisense probes along with the *neurog1* antisense probes to perform the FISH staining with 48 hpf samples because reduction of *fgf3* staining signals in the developing pharyngeal pouches was an evident feature of *wls* mutants at 36 and 48 hpf (Supplemental Figure 4A-F) (Wu et al., 2015). ZFIN (Zebrafish Information Network) Gene IDs for *cpd2, vachtb*, *tac3a, vglut2a, gng8, neurog1*, *her6*, *fgf3*, *wls* or *clu* are ZDB-GENE-030903-1, ZDB-GENE-040426-1410, ZDB-GENE-090313-253, ZDB-GENE-030616-554, ZDB-GENE-050522-312, ZDB-GENE-990415-174, ZDB-GENE-980526-144, ZDB-GENE-980526-178, ZDB-GENE-040426-2161, ZDB-GENE-040426-1774, respectively. The two-color whole mount fluorescent RNA *in situ* hybridization (FISH) was carried out as previously described by Wang et al. with minor modifications (Wang et al., 2021). One-color FISH plus one-color immunohistochemical staining (IHC) was conducted basically similar to the two-color FISH except that 1 to 200 dilutions of rabbit anti-EGPF (Torrey Pines TP401, USA) or anti-Wls antibodies were co-incubated with anti-DIG-POD or anti-FLU-POD antibodies (Roche 11207733910 and Roche 11426346910), followed by Tyrimide substrate stainining (Perkin Elmer NEL753001 Kit) for 60 minutes then followed by incubating with 1 to 200 dilutions of goat anti-Rabbit IgG Alexa-488 or goat anti-Rabbit IgG Alexa-568 (ThermoFisher/Invitrogen A-11008 or A-11011) (Wu et al., 2015).

#### Image Acquisition, Processing, visualization and quantification

FISH or FISH plus IHC staining samples were incubated overnight in 50% glycerol containing 2 % PFA then imaged with Leica TCS SP5 or SP8 (IR-LEGO data) confocal systems (Leica GmbH, Germany) using a 10x (Figure 3) or 20X (Figure 1-2, 4-7) air objectives with 0.3 or 0.5 N.A. Each samples were scanned only once with 2x (Figure 4-7), 3x (Figure 1-2) or 4x (Figure 3) zooming factors, and recorded as 512 x 512 pixels and Z-step of 3.0 or 4.0 μm (Figure 3 or Figure 1-2, 4-7) for 24 (Figure 1), 31 (Figure 2), 27 (Figure 3), 34 (Figure 4-7) steps (final 25, 32, 28 or 35 frames) at 200 Hz. The resulted voxel size of our images are 0.76 x 0.76 x 3.0 (Figure 1), 0.76 x 0.76 x 4.0 (Figure 2) or 0.76 x 0.76 x 3.0 (Figure 3-7) μm^3^. For presentation, Z-projection images were first obtained using “3D projection” function in Leica’s LAS AF software then they were directly exported as TIF files then some of these images were cropped to proper size using Photoshop CS5 (Adobe Inc., USA).

Rigid image registrations, Correlation Factors (CFs, image similarities) or gene expression volumes (vol.) were calculated as previously described utilizing Amira 5.6 software (FEI, OR, USA) (Wang et al., 2021). Intensity threshold values (0-255) adopted for gene expression vol. calculation and image visualization (Iso-surface views) were listed in each related supplemental figure legends (Supplemental Figure 1, 2, 3, 5, 6). Signals from each samples were taken once only to avoid the bleaching effects that were obviously affecting the vol. calculations for the signals emitted from FLU-conjugated Tyrimides (green channel) (Wang et al., 2021). Images exhibiting different viewing angles of the virtual 3D maps were exported as JPG files from viewer window using the Snapshot function placed in Amira software, then JPG images were cropped to proper size using Photoshop CS5 (Adobe Inc., USA). 3D proliferating cell counting was conducted manually utilizing the Cell Counter Plug-In of Fiji-ImageJ (Supplemental Figure 7) (Schindelin, Arganda-Carreras et al., 2012).

#### Infrared laser-mediated or Kaede-mediated cell tracing

Infrared laser-evoked gene operator (IR-LEGO) was conducted following previously reported protocols with some modifications (Deguchi et al., 2009, Kamei et al., 2009, Shimada et al., 2013). Basically utilizing transient expression of *neurog1*:*egfp* in the *hsp70*:mCherry transgenic embryos through plasmid injection (25 pg/egg), we first identified the *neurog1*-positive cells (EGFP-positive) in the HA progenitor zones at 48 hpf. Then we labeled 1-2 cells by laser-mediated heatshock induction (IR-LEGO) of mCherry expressions under an 20x objective lens (NA=0.75, laser power received is 18-21.8mW) installed at the inverted microscope IX80 (Olympus, Japan) and trace where the EGFP-mCherry double-positive cells would be at 72 hpf by taking image Z-stacks of HA regions with Leica SP8 confocal microscope system utilizing 20x objective. The Kaede-mediated cell-tracing experiments were conducted based on published methods (Gfrerer, Dougherty et al., 2013). One to two nanoliters (nl) of Tol2-*neurog1*: *kaede* plasmid (25 pg/egg) was injected into AB eggs at the one-cell stage. Injected embryos exhibiting mosaic green Kaede signals in the HA progenitor zones were quickly harvested and anesthetize in egg water with 0.015% Tricaine for 5 min then mounted with 1 % low melting agarose gel (with 0.015% Tricaine). Mounted 36hpf embryos then were subjected to photo-conversion by UV laser (<405 nm) exposure for 200 msec. Z-stack confocal images of treated embryos were taken right before and 12 or 24 hours after UV exposure with Leica SP5 confocal microscope system (Leica GmbH, Germany).

#### Statistical Analysis

Statistics for gene expression volume averaging, standard deviation (SD) calculation and heatmap generation were performed utilizing Prism 7 software (Graphpad, CA, USA). Unpaired Student’s t-test (two tailed) was utilized to determine significance at the *P* value that is smaller than 0.05. Confidence for *P* value (*p*) is denoted by “*” that equals to *p*< 0.05 and “**” that equals to *p*< 0.01.

#### Availability of data and materials

The materials generated or datasets used during the current study are available from the corresponding author on reasonable request (Material Transfer Agreements may be required.).

## Authors’ contributions

S-H Lin and Y-S Kuan designed the experiments. S-H Lin, H-Y Pan, B-T Wu, C-H Wu, J Sakamoto, A Shimada and Y-S Kuan performed the experiments. Y Kamei, H Takeda and Y-S Kuan provided the reagents. S-H Lin, H-Y Pan and Y-S Kuan analyzed the data. Y-S Kuan wrote the manuscript with the inputs from Y Kamei and H Takeda.

## Acknowledgements

We thank Taiwan Zebrafish Core Facility funded by National Science and Technology Council, (NSTC, Taiwan, R.O.C.) Grant #113-2740-B-400-001 for fish line maintenance and reagents. We also thank Drs. Sheng-Ping L. Hwang, Ya-Hui Chou (ICOB, Academia Sinica, Taiwan) and Bon-Chu Chung (IMB, Academia Sinica, Taiwan) for critical discussion; Drs. C.B. Moens (Fred Hutch Cancer Center, Seattle, WA, U.S.A.), M. E. Halpern (Carnegie Inst. for Science, Baltimore, MD, U.S.A.) and Koichi Kawakami, (NIG, Shizuoka, Japan) for reagents; Ms. Chie Kinoshita (NIBB, Okazaki, Japan), Ms. Misako Saida (NIBB, Okazaki, Japan), Mr. Bo-Hung Lin (ICOB, Academia Sinica, Taiwan) and Dr. Sheng-Wei Lin (IBC, Academia Sinica, Taiwan) for technical supports.

