## Supplemental Figures and Tables for "Wntless interacts with Notch signaling to balance the generation of neurons and gliocytes in vertebrate dorsal diencephalon"

**Supplemental Figure 1. Correlation factors from image registration results for *cpd2* mRNA-labeled samples.**

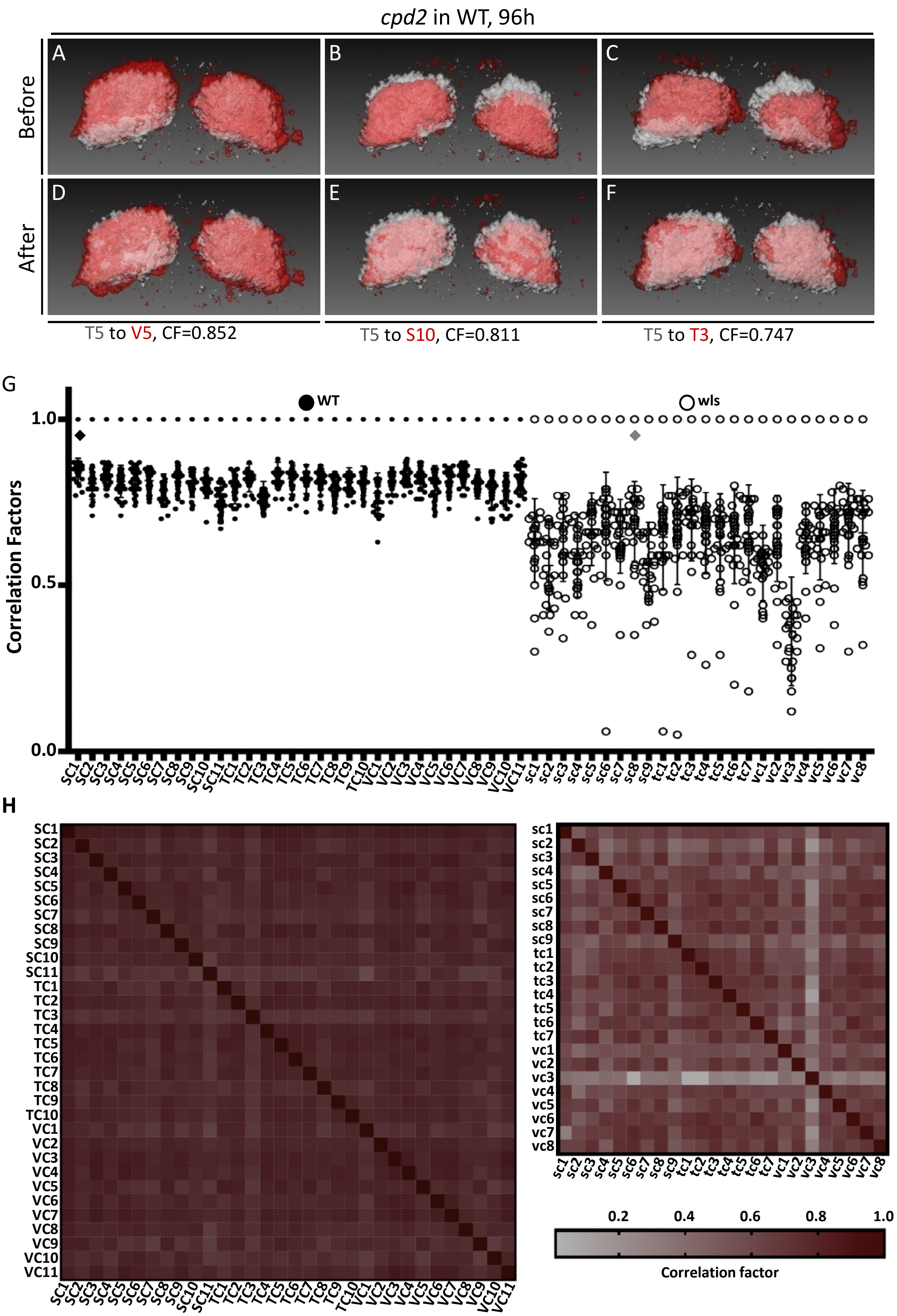

**Supplemental Figure 2. Correlation factors from image registration results for *gng8* mRNA-labeled samples.**

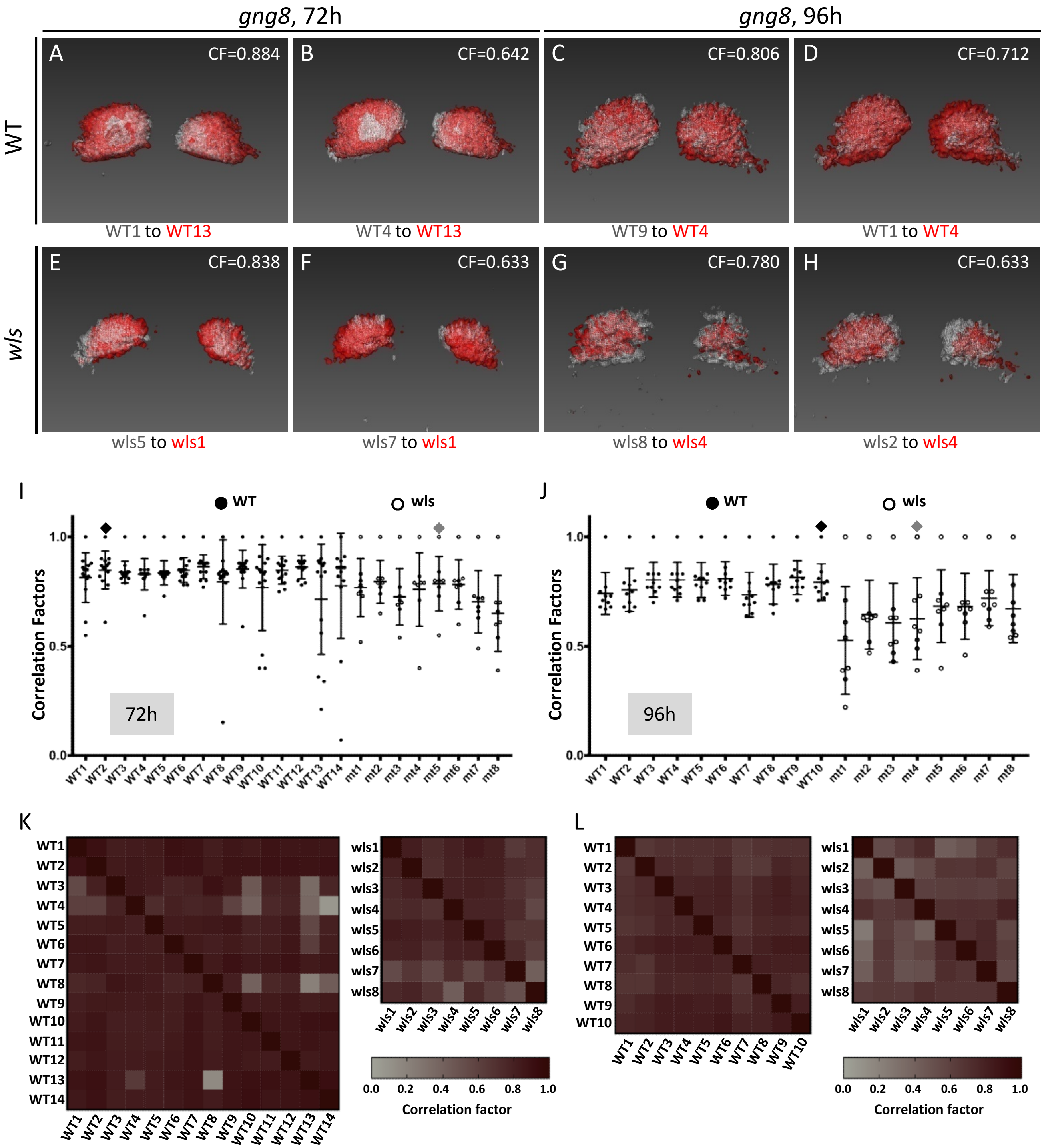

**Supplemental Figure 3. Correlation factors from image registration results for *ngn1* mRNA-labeled samples.**

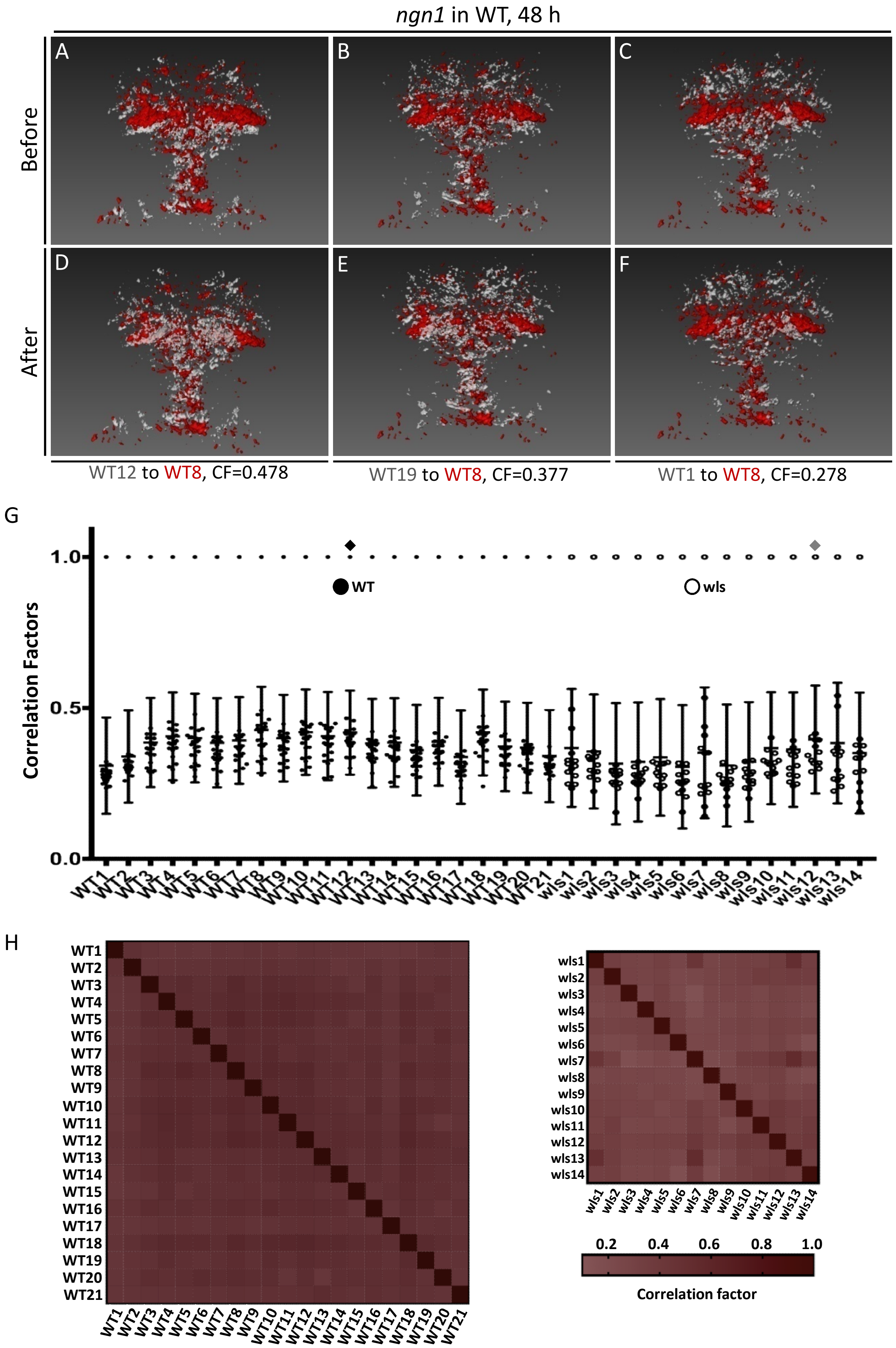

**Supplemental Figure 4. *ngn1*-positive progenitors developed into HA neurons.**

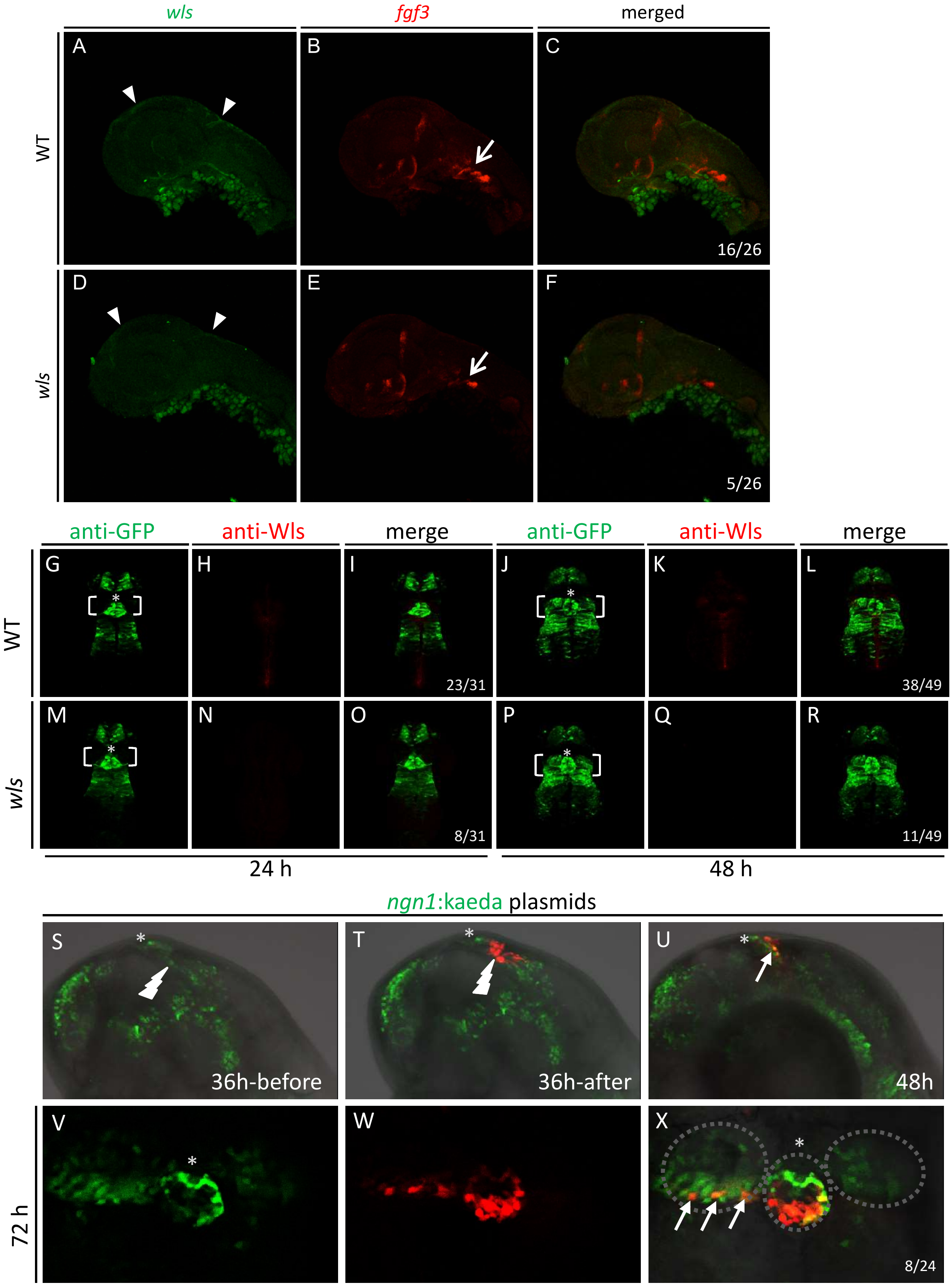

**Supplemental Figure 5. Correlation factors from image registration results for *her6* mRNA-labeled samples.**

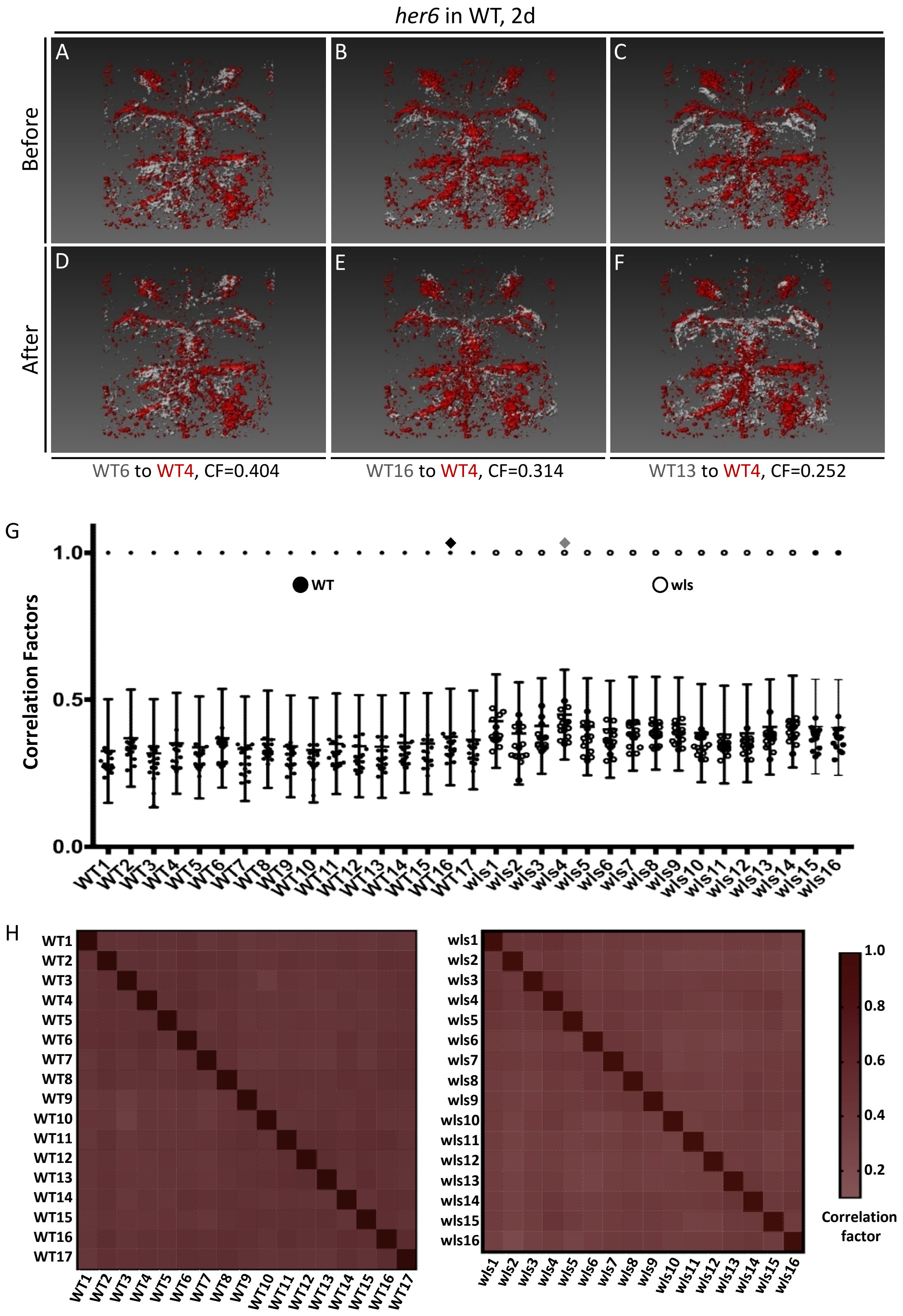

**Supplemental Figure 6. Correlation factors from image registration results for foxD3:GFP-labeled samples.**

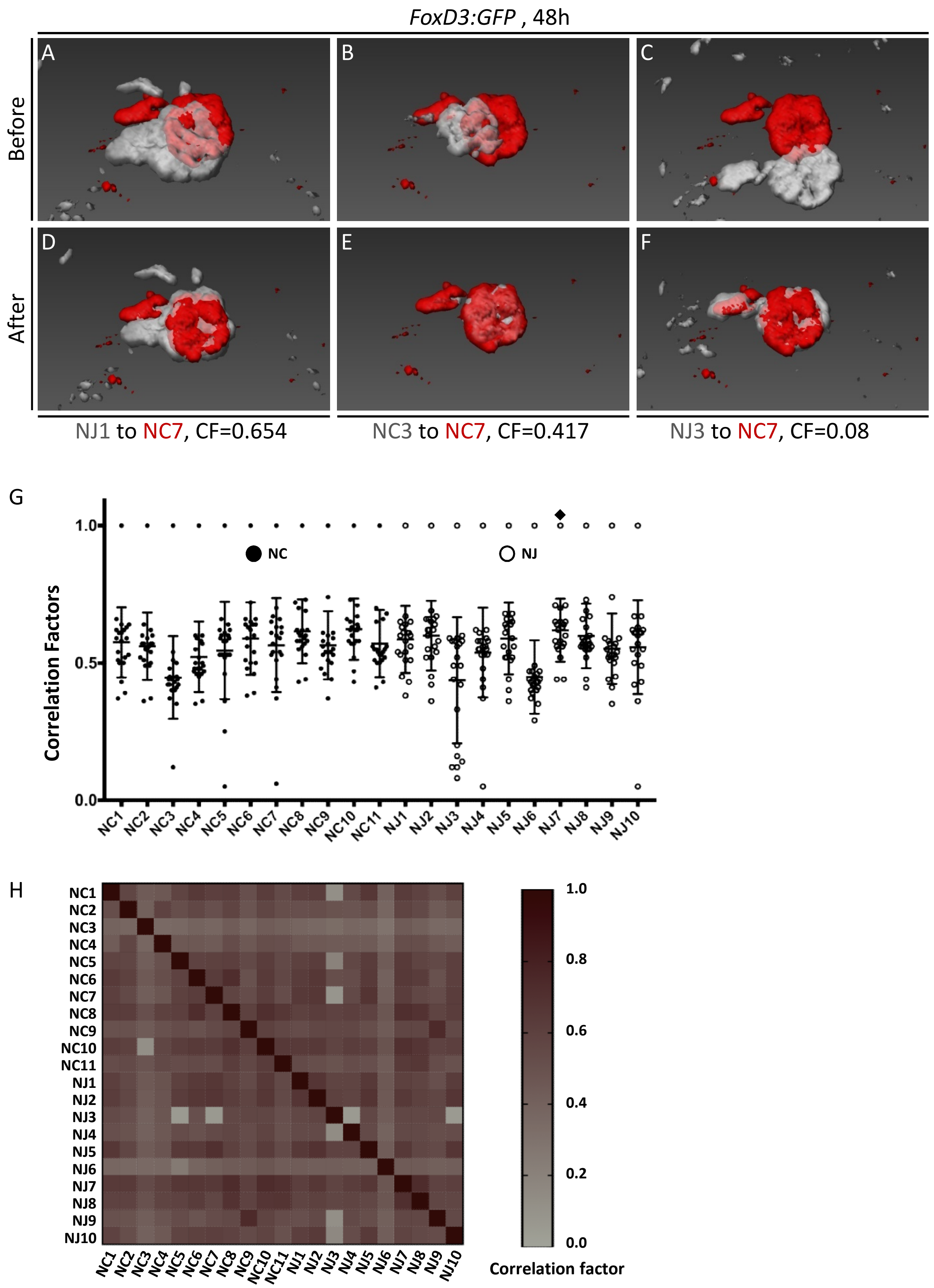

**Supplemental Figure 7. Cell proliferations in HA progenitor zone are decreased in *wls* mutants.**

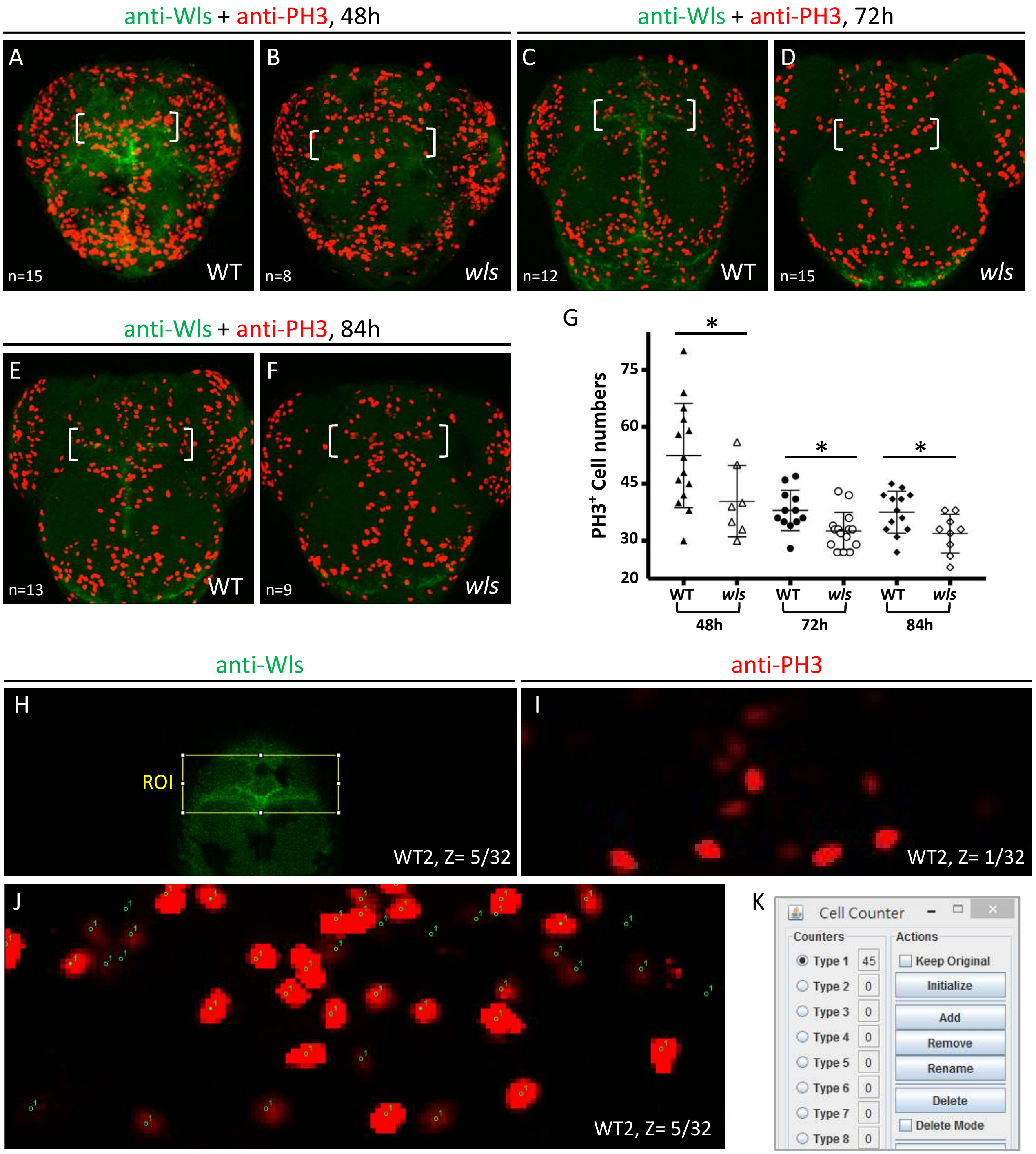


L

| | WT1 | WT2 | WT3 | WT4 | WT5 | WT6 | WT7 | WT8 | WT9 | WT10 | WT11 | WT12 | WT13 | WT14 | WT15 | Ave. |
|---|---|---|---|---|---|---|---|---|---|---|---|---|---|---|---|---|
| 48h | 58 | 45 | 62 | 65 | 52 | 69 | 30 | 80 | 38 | 32 | 42 | 59 | 40 | 46 | 48 | 51.1 |
| 72h | 35 | 38 | 47 | 41 | 38 | 34 | 28 | 46 | 36 | 36 | 35 | 42 | | | | 38.0 |
| 84h | 42 | 36 | 35 | 31 | 41 | 38 | 42 | 27 | 33 | 41 | 44 | 33 | 45 | | | 37.5 |

M

| | WT1 | WT2 | WT3 | WT4 | WT5 | WT6 | WT7 | WT8 | WT9 | WT10 | WT11 | WT12 | WT13 | WT14 | WT15 | Ave. |
|---|---|---|---|---|---|---|---|---|---|---|---|---|---|---|---|---|
| 48h | 56 | 33 | 50 | 35 | 30 | 40 | 39 | | | | | | | | | 40.4 |
| 72h | 33 | 27 | 33 | 34 | 36 | 33 | 42 | 29 | 27 | 31 | 33 | 32 | 43 | 29 | 29 | 32.6 |
| 84h | 32 | 35 | 33 | 29 | 33 | 23 | 26 | 38 | 38 | | | | | | | 31.9 |

**Supplemental Figure 1. Correlation factors from image registration results for *cpd2* mRNA-labeled samples.** (A-F) Representative snapshots from one-to-one 3D image registration tests for the *cpd2* expression domains in habenula nuclei (HA) in 96 hpf (96h) wildtype (WT) larva, before (A-C) or after (D-F) registration. All are dorsal views. (G) Statistical chart of correlation factors (CFs) from each Z-stacks of *cpd2* after one-to-one registration. Diamonds mark the selected reference Z-stacks in WT or *wls* mutant (wls) groups (n=32 and n=24, respectively). (H) Cluster analyses (heatmaps) of the CFs from one-to-one registration between the Z-stacks of *cpd2* gene. Left or the top-right panels are analysis results for WT or *wls* groups. Bottom-right panel is the color code for the two heatmaps. (See Supplemental Table 1 for numerical values)

**Supplemental Figure 2. Correlation factors from image registration results for *gng8* mRNA-labeled samples.** (A-H) Representative snapshots from one-to-one 3D image registration for the *gng8* expression domains in the habenular nuciei (HA) in 72h or 96h wildtype (WT, A-D) or *wls* mutant (*wls*, E-H) larva after registration. Intensity threshold I=80 for A to D and I=40 for E to H. All images are dorsal views. (I-J) Statistical charts of image correlation factors (CFs) from each Z-stacks of *gng8* after one-to-one registration. (K-L) Cluster analyses (heatmaps) of the CFs from one-to-one registration between the Z-stacks of *gng8* gene (72h group in K, 96h group in L). Left or the top-right panels are analysis results for WT or *wls* groups, respectively. Bottom-right panel is the color codes for the heatmaps (See Supplemental Table 2 for numerical values).

**Supplemental Figure 3. Correlation factors from image registration results for *ngn1* mRNA-labeled samples.** (A-F) Representative snapshots from one-to-one 3D image registration tests for the *ngn1* expression domains in diencephalic sub-ventricular zones in 2 dpf (2d) embryos, before (A-C) or after (D-F) registration. All are dorsal views. (G) Statistical chart of correlation factors (CFs) from each Z-stacks of *ngn1* after one-to-one registration. Diamonds mark the selected reference Z-stacks in wildtype (WT) or *wls* mutant (wls) groups(n=21 and n=14, respectively). (H) Cluster analyses (heatmaps) of the CFs from one-to-one registration between the Z-stacks of *ngn1* gene. Left or the top-right panels are analysis results for WT or wls groups. Bottom-right panel is the color codes for the heatmaps (See Supplementary Table 3 for numerical values).

**Supplemental Figure 4. *ngn1*-positive progenitors developed into HA neurons.** (A-F) Representative snapshots (lateral views) from *wls* (in green) and *fgf3* (in red) FISH labeling in the dorsal brain (*wls*, indicated by arrow heads) and pharyngeal pouches (*fgf3*, indicated by arrows) of 36h WT (A to C) or *wls* (D to F) embryos. Total sample size is 26, and the proportion of WT or *wls* samples is indicated at the bottom right corners in (C) and (F). (G-R) Images showing *ngn1*-expressing cells (*ngn1*:GFP, labeled by anti-GFP) looked normal in the WT (G-L) or *wls* mutants (M-R) at 24 (G-I, M-O) or 48h (J-L,P-R). "*" indicates pineal; brackets indicate the left and right HA. See Supplemental Table 4 for numeric data. (S-X) Images showing *ngn1*:kaeda injected embryo before (S) or after color version (green to red) (T) by blue light laser at 36h (lateral views). (U-X) Image showing the converted Kaeda-expressing cells (in red) in (T) developed into HA neurons at 48h (U) (lateral view) and 72 h (V-X) (dorsal views).

**Supplemental Figure 5. Correlation factors from image registration results for *her6* mRNA-labeled samples.** (A-F) Representative snapshots from one-to-one 3D image registration tests for the *her6* expression domains in diencephalic sub-ventricular zones in 48 hpf (48h) embryos, before (A-C) or after (D-F) registration. All are dorsal views. (G) Statistical chart of correlation factors (CFs) from each Z-stacks of *her6* after one-to-one registration. Diamonds mark the selected reference Z-stacks in wildtype (WT) or *wls* mutant (wls) groups(n=17 and n=16, respectively). (H) Cluster analyses (heatmaps) of the CFs from one-to-one registration between the Z-stacks of *her6* gene. Left or the top-right panels are analysis results for WT or *wls* groups. Bottom-right panel is the color codes for the heatmaps (See Supplementary Table 5, 6 for numerical values).

**Supplemental Figure 6. Correlation factors from image registration results for foxD3:GFP-labeled samples.** (A-F) Representative snapshots from one-to-one 3D image registration tests for the GFP expression domains in pineal and parapineal in embryos at 48 hours post-fertilization (48h), before (A-C) or after (D-F) registration. All are dorsal views. (G) Statistical chart of correlation factors (CFs) from each Z-stacks from control and *her6*-plasmid injected samples after one-to-one registration. Black diamond marks the selected reference Z-stack. Total sample size is 21. (H) Cluster analyses (heatmaps) of the CFs from one-to-one registration between the Z-stacks of GFP signals (mark pineals). (See Supplementary Table 7 for numerical values)

**Supplemental Figure 7. Cell proliferations in HA progenitor zone are decreased in *wls* mutants.** (A-F) Images showing proliferating cells labeled by anti-PH3 (red) in the WT (A, C, E, anti-Wls-positive) or *wls* mutant embryos (B, D, F) at 48 (A-B), 72 (C-D) or 84 (E-F) hours (h) post-fertilization. Brackets indicate the regions of the HA progenitor zone (P-zone) in dorsal diencephalion (D-Dien). (G) Statistical chart for the counts of anti-PH3-positive cells in 3D HA P-zone. Confidence *p* value (*p*) is denoted by “*”. “*” means $P<0.05$ in t-test. See the numeric data in (L-M). (H) A snapshot showing the fifth slice from a confocal Z-stack (Z=5/32) of a WT embryo (WT2 here) labeled by anti-Wls at 48h. Yellow rectangle indicates the region of interest (ROI) for cropping the confocal Z-stack to isolate the 3D HA P-zone in D-Dien. (I-J) Snapshots of the first and fifth slices showing the proliferating cells labeled with anti-PH3 in the ROI from (H) and the cell counting result. Some cells were marked by green dots (“1” means “type 1”.) but were out of focus here would be clearly showed in other Z-slices. (K) Total cell counts (45 for WT2) in the Cell Counter control window of Fiji-ImageJ. (L-M) Tables showing the cell counts from WT (L) or *wls* mutant (M) embryos at 48, 72 or 84h.

**Supplemental Table 1. Computed correlation factors (CFs) from reciprocal one-to-one registration for each Z-stacks of *cpd2*, and the expression volumes of all four HA genes used in figure 1**. (A-B) CFs for WT (A, SC1 to VC 11) or *wls* mutant (B, sc1 to vc9) groups. (C-E) Volumes (vol.) of *slc18a3b* (S or s), *tac3a* (T or t), *vglut2* (V or v)(top 2 panels in C-E) or *cpd2* (C or c) (bottom 2 panels in C-E) expression domain ($\mu m^3$) from samples of WT ($1^{st}$ and $3^{rd}$ panels in C-E) or *wls* mutants ($2^{nd}$ and $4^{th}$ panels in C-E). (F) Gene expression vol. ratios. L is left, R is right HA domain.Intensity threshold (I) = 35, 8, 35 and 10 for S, T, V and C gene expression vol. calculation.

(A)

| Name | SC1 | SC2 | SC3 | SC4 | SC5 | SC6 | SC7 | SC8 | SC9 | SC10 | SC11 | TC1 | TC2 | TC3 | TC4 | TC5 | TC6 |
|---|---|---|---|---|---|---|---|---|---|---|---|---|---|---|---|---|---|
| SC1 | 1.00 | 0.86 | 0.86 | 0.85 | 0.85 | 0.85 | 0.80 | 0.87 | 0.83 | 0.85 | 0.77 | 0.84 | 0.87 | 0.79 | 0.88 | 0.87 | 0.86 |
| SC2 | 0.85 | 1.00 | 0.81 | 0.83 | 0.79 | 0.80 | 0.74 | 0.83 | 0.79 | 0.79 | 0.71 | 0.81 | 0.82 | 0.74 | 0.82 | 0.85 | 0.84 |
| SC3 | 0.85 | 0.81 | 1.00 | 0.82 | 0.85 | 0.83 | 0.80 | 0.83 | 0.83 | 0.80 | 0.79 | 0.79 | 0.83 | 0.77 | 0.84 | 0.83 | 0.82 |
| SC4 | 0.85 | 0.83 | 0.82 | 1.00 | 0.79 | 0.80 | 0.75 | 0.83 | 0.78 | 0.80 | 0.71 | 0.83 | 0.83 | 0.74 | 0.83 | 0.84 | 0.84 |
| SC5 | 0.85 | 0.79 | 0.86 | 0.79 | 1.00 | 0.85 | 0.81 | 0.81 | 0.84 | 0.81 | 0.79 | 0.76 | 0.82 | 0.77 | 0.84 | 0.79 | 0.82 |
| SC6 | 0.84 | 0.80 | 0.82 | 0.80 | 0.85 | 1.00 | 0.80 | 0.80 | 0.83 | 0.82 | 0.74 | 0.76 | 0.81 | 0.78 | 0.84 | 0.80 | 0.82 |
| SC7 | 0.80 | 0.74 | 0.80 | 0.75 | 0.81 | 0.80 | 1.00 | 0.77 | 0.83 | 0.80 | 0.75 | 0.74 | 0.77 | 0.76 | 0.80 | 0.76 | 0.76 |
| SC8 | 0.87 | 0.83 | 0.84 | 0.84 | 0.81 | 0.81 | 0.78 | 1.00 | 0.79 | 0.82 | 0.74 | 0.83 | 0.84 | 0.75 | 0.85 | 0.86 | 0.83 |
| SC9 | 0.83 | 0.79 | 0.83 | 0.78 | 0.84 | 0.83 | 0.83 | 0.79 | 1.00 | 0.81 | 0.78 | 0.77 | 0.80 | 0.79 | 0.81 | 0.79 | 0.81 |
| SC10 | 0.85 | 0.79 | 0.80 | 0.80 | 0.80 | 0.82 | 0.80 | 0.82 | 0.81 | 1.00 | 0.69 | 0.80 | 0.81 | 0.77 | 0.83 | 0.81 | 0.82 |
| SC11 | 0.77 | 0.71 | 0.79 | 0.71 | 0.79 | 0.75 | 0.75 | 0.73 | 0.78 | 0.69 | 1.00 | 0.70 | 0.74 | 0.71 | 0.76 | 0.73 | 0.72 |
| TC1 | 0.83 | 0.81 | 0.79 | 0.84 | 0.76 | 0.77 | 0.74 | 0.83 | 0.77 | 0.80 | 0.71 | 1.00 | 0.82 | 0.73 | 0.81 | 0.85 | 0.81 |
| TC2 | 0.86 | 0.82 | 0.83 | 0.83 | 0.82 | 0.81 | 0.77 | 0.84 | 0.80 | 0.81 | 0.74 | 0.82 | 1.00 | 0.77 | 0.85 | 0.84 | 0.83 |
| TC3 | 0.78 | 0.74 | 0.77 | 0.74 | 0.77 | 0.78 | 0.76 | 0.75 | 0.79 | 0.77 | 0.71 | 0.73 | 0.76 | 1.00 | 0.76 | 0.74 | 0.76 |
| TC4 | 0.87 | 0.82 | 0.85 | 0.84 | 0.84 | 0.84 | 0.80 | 0.86 | 0.81 | 0.84 | 0.76 | 0.81 | 0.85 | 0.76 | 1.00 | 0.84 | 0.84 |
| TC5 | 0.86 | 0.85 | 0.83 | 0.85 | 0.79 | 0.80 | 0.76 | 0.86 | 0.79 | 0.81 | 0.74 | 0.85 | 0.84 | 0.75 | 0.84 | 1.00 | 0.84 |
| TC6 | 0.86 | 0.84 | 0.83 | 0.84 | 0.82 | 0.83 | 0.76 | 0.83 | 0.81 | 0.82 | 0.72 | 0.81 | 0.83 | 0.76 | 0.84 | 0.84 | 1.00 |
| TC7 | 0.86 | 0.81 | 0.83 | 0.83 | 0.80 | 0.79 | 0.77 | 0.85 | 0.79 | 0.81 | 0.73 | 0.83 | 0.84 | 0.74 | 0.84 | 0.84 | 0.82 |
| TC8 | 0.82 | 0.75 | 0.83 | 0.76 | 0.84 | 0.81 | 0.78 | 0.78 | 0.80 | 0.78 | 0.77 | 0.74 | 0.80 | 0.75 | 0.81 | 0.77 | 0.77 |
| TC9 | 0.84 | 0.77 | 0.83 | 0.81 | 0.80 | 0.79 | 0.79 | 0.83 | 0.80 | 0.79 | 0.77 | 0.81 | 0.82 | 0.74 | 0.83 | 0.83 | 0.80 |
| TC10 | 0.85 | 0.81 | 0.83 | 0.82 | 0.80 | 0.79 | 0.77 | 0.84 | 0.79 | 0.81 | 0.75 | 0.82 | 0.83 | 0.74 | 0.83 | 0.84 | 0.83 |
| VC1 | 0.82 | 0.80 | 0.77 | 0.82 | 0.74 | 0.76 | 0.71 | 0.81 | 0.73 | 0.78 | 0.67 | 0.82 | 0.82 | 0.71 | 0.80 | 0.82 | 0.80 |
| VC2 | 0.85 | 0.81 | 0.84 | 0.81 | 0.82 | 0.80 | 0.79 | 0.83 | 0.82 | 0.80 | 0.77 | 0.79 | 0.82 | 0.77 | 0.83 | 0.82 | 0.80 |
| VC3 | 0.86 | 0.82 | 0.87 | 0.82 | 0.86 | 0.85 | 0.81 | 0.83 | 0.85 | 0.81 | 0.81 | 0.80 | 0.84 | 0.79 | 0.85 | 0.83 | 0.84 |
| VC4 | 0.85 | 0.79 | 0.87 | 0.81 | 0.87 | 0.84 | 0.82 | 0.83 | 0.84 | 0.79 | 0.81 | 0.78 | 0.82 | 0.76 | 0.85 | 0.82 | 0.83 |
| VC5 | 0.85 | 0.81 | 0.82 | 0.84 | 0.78 | 0.76 | 0.74 | 0.85 | 0.75 | 0.79 | 0.73 | 0.85 | 0.84 | 0.72 | 0.83 | 0.85 | 0.81 |
| VC6 | 0.84 | 0.79 | 0.85 | 0.79 | 0.84 | 0.81 | 0.79 | 0.80 | 0.83 | 0.78 | 0.80 | 0.76 | 0.81 | 0.77 | 0.81 | 0.80 | 0.81 |
| VC7 | 0.87 | 0.82 | 0.87 | 0.82 | 0.86 | 0.84 | 0.81 | 0.84 | 0.83 | 0.82 | 0.79 | 0.80 | 0.84 | 0.77 | 0.85 | 0.83 | 0.84 |
| VC8 | 0.84 | 0.80 | 0.81 | 0.81 | 0.82 | 0.83 | 0.79 | 0.81 | 0.81 | 0.83 | 0.70 | 0.78 | 0.81 | 0.77 | 0.83 | 0.80 | 0.83 |
| VC9 | 0.84 | 0.84 | 0.79 | 0.82 | 0.75 | 0.77 | 0.70 | 0.84 | 0.74 | 0.78 | 0.69 | 0.81 | 0.83 | 0.73 | 0.81 | 0.84 | 0.81 |
| VC10 | 0.81 | 0.77 | 0.80 | 0.75 | 0.83 | 0.84 | 0.80 | 0.76 | 0.83 | 0.80 | 0.71 | 0.73 | 0.77 | 0.78 | 0.79 | 0.76 | 0.80 |
| VC11 | 0.84 | 0.79 | 0.86 | 0.79 | 0.87 | 0.84 | 0.81 | 0.80 | 0.85 | 0.79 | 0.82 | 0.76 | 0.82 | 0.78 | 0.83 | 0.80 | 0.81 |

| Name | TC7 | TC8 | TC9 | TC10 | VC1 | VC2 | VC3 | VC4 | VC5 | VC6 | VC7 | VC8 | VC9 | VC10 | VC11 | Average |
|---|---|---|---|---|---|---|---|---|---|---|---|---|---|---|---|---|
| SC1 | 0.86 | 0.82 | 0.84 | 0.85 | 0.82 | 0.85 | 0.87 | 0.85 | 0.85 | 0.84 | 0.87 | 0.84 | 0.84 | 0.81 | 0.84 | 0.84 |
| SC2 | 0.81 | 0.75 | 0.77 | 0.81 | 0.80 | 0.81 | 0.82 | 0.79 | 0.81 | 0.79 | 0.82 | 0.80 | 0.84 | 0.77 | 0.79 | 0.80 |
| SC3 | 0.82 | 0.83 | 0.83 | 0.83 | 0.77 | 0.84 | 0.87 | 0.86 | 0.82 | 0.85 | 0.87 | 0.81 | 0.79 | 0.80 | 0.86 | 0.82 |
| SC4 | 0.83 | 0.76 | 0.81 | 0.82 | 0.81 | 0.81 | 0.82 | 0.81 | 0.84 | 0.78 | 0.82 | 0.81 | 0.82 | 0.75 | 0.78 | 0.80 |
| SC5 | 0.81 | 0.84 | 0.81 | 0.80 | 0.75 | 0.82 | 0.86 | 0.87 | 0.78 | 0.84 | 0.86 | 0.82 | 0.76 | 0.83 | 0.87 | 0.82 |
| SC6 | 0.79 | 0.80 | 0.79 | 0.79 | 0.75 | 0.80 | 0.85 | 0.83 | 0.76 | 0.80 | 0.84 | 0.83 | 0.76 | 0.83 | 0.83 | 0.81 |
| SC7 | 0.77 | 0.78 | 0.79 | 0.77 | 0.71 | 0.79 | 0.80 | 0.82 | 0.74 | 0.79 | 0.81 | 0.79 | 0.70 | 0.79 | 0.80 | 0.78 |
| SC8 | 0.85 | 0.78 | 0.83 | 0.84 | 0.81 | 0.83 | 0.83 | 0.83 | 0.85 | 0.81 | 0.84 | 0.81 | 0.84 | 0.76 | 0.80 | 0.82 |
| SC9 | 0.79 | 0.80 | 0.79 | 0.79 | 0.73 | 0.82 | 0.85 | 0.83 | 0.75 | 0.83 | 0.83 | 0.81 | 0.74 | 0.83 | 0.85 | 0.80 |
| SC10 | 0.81 | 0.78 | 0.79 | 0.81 | 0.78 | 0.80 | 0.81 | 0.79 | 0.79 | 0.78 | 0.82 | 0.83 | 0.78 | 0.80 | 0.79 | 0.80 |
| SC11 | 0.73 | 0.76 | 0.77 | 0.75 | 0.63 | 0.77 | 0.80 | 0.81 | 0.73 | 0.80 | 0.79 | 0.70 | 0.69 | 0.71 | 0.82 | 0.74 |
| TC1 | 0.83 | 0.75 | 0.82 | 0.82 | 0.82 | 0.80 | 0.80 | 0.78 | 0.84 | 0.77 | 0.80 | 0.78 | 0.81 | 0.73 | 0.76 | 0.79 |
| TC2 | 0.84 | 0.80 | 0.82 | 0.83 | 0.82 | 0.82 | 0.84 | 0.82 | 0.84 | 0.81 | 0.84 | 0.81 | 0.83 | 0.77 | 0.82 | 0.82 |
| TC3 | 0.74 | 0.74 | 0.74 | 0.74 | 0.71 | 0.77 | 0.78 | 0.76 | 0.72 | 0.77 | 0.77 | 0.77 | 0.73 | 0.78 | 0.78 | 0.75 |
| TC4 | 0.84 | 0.81 | 0.83 | 0.83 | 0.81 | 0.83 | 0.85 | 0.85 | 0.83 | 0.81 | 0.85 | 0.83 | 0.81 | 0.79 | 0.83 | 0.83 |
| TC5 | 0.84 | 0.77 | 0.83 | 0.84 | 0.82 | 0.82 | 0.83 | 0.82 | 0.85 | 0.80 | 0.83 | 0.80 | 0.84 | 0.76 | 0.80 | 0.82 |
| TC6 | 0.82 | 0.77 | 0.81 | 0.83 | 0.80 | 0.80 | 0.84 | 0.83 | 0.81 | 0.81 | 0.83 | 0.83 | 0.81 | 0.80 | 0.81 | 0.81 |
| TC7 | 1.00 | 0.79 | 0.83 | 0.83 | 0.82 | 0.81 | 0.82 | 0.82 | 0.85 | 0.80 | 0.83 | 0.80 | 0.81 | 0.75 | 0.80 | 0.81 |
| TC8 | 0.79 | 1.00 | 0.77 | 0.77 | 0.72 | 0.80 | 0.83 | 0.82 | 0.77 | 0.82 | 0.81 | 0.78 | 0.73 | 0.79 | 0.83 | 0.79 |
| TC9 | 0.83 | 0.77 | 1.00 | 0.83 | 0.78 | 0.81 | 0.81 | 0.84 | 0.83 | 0.80 | 0.83 | 0.79 | 0.78 | 0.74 | 0.81 | 0.80 |
| TC10 | 0.83 | 0.77 | 0.83 | 1.00 | 0.80 | 0.81 | 0.83 | 0.83 | 0.83 | 0.81 | 0.84 | 0.78 | 0.80 | 0.76 | 0.81 | 0.81 |
| VC1 | 0.82 | 0.72 | 0.78 | 0.80 | 1.00 | 0.75 | 0.77 | 0.76 | 0.82 | 0.73 | 0.77 | 0.77 | 0.80 | 0.70 | 0.74 | 0.77 |
| VC2 | 0.81 | 0.80 | 0.81 | 0.81 | 0.75 | 1.00 | 0.84 | 0.83 | 0.82 | 0.83 | 0.83 | 0.79 | 0.79 | 0.78 | 0.83 | 0.81 |
| VC3 | 0.82 | 0.83 | 0.81 | 0.83 | 0.78 | 0.84 | 1.00 | 0.86 | 0.80 | 0.86 | 0.86 | 0.81 | 0.80 | 0.82 | 0.88 | 0.83 |
| VC4 | 0.82 | 0.82 | 0.84 | 0.83 | 0.76 | 0.83 | 0.87 | 1.00 | 0.81 | 0.84 | 0.86 | 0.82 | 0.76 | 0.80 | 0.86 | 0.82 |
| VC5 | 0.85 | 0.77 | 0.83 | 0.83 | 0.82 | 0.82 | 0.80 | 0.81 | 1.00 | 0.78 | 0.81 | 0.77 | 0.83 | 0.71 | 0.77 | 0.80 |
| VC6 | 0.79 | 0.81 | 0.80 | 0.81 | 0.73 | 0.83 | 0.86 | 0.84 | 0.78 | 1.00 | 0.85 | 0.78 | 0.75 | 0.80 | 0.86 | 0.81 |
| VC7 | 0.83 | 0.81 | 0.83 | 0.84 | 0.77 | 0.83 | 0.86 | 0.87 | 0.81 | 0.85 | 1.00 | 0.82 | 0.79 | 0.81 | 0.86 | 0.83 |
| VC8 | 0.80 | 0.78 | 0.79 | 0.78 | 0.77 | 0.79 | 0.81 | 0.82 | 0.77 | 0.78 | 0.82 | 1.00 | 0.77 | 0.81 | 0.80 | 0.80 |
| VC9 | 0.81 | 0.73 | 0.78 | 0.80 | 0.80 | 0.79 | 0.80 | 0.76 | 0.83 | 0.76 | 0.79 | 0.77 | 1.00 | 0.70 | 0.77 | 0.78 |
| VC10 | 0.75 | 0.79 | 0.75 | 0.76 | 0.70 | 0.77 | 0.82 | 0.80 | 0.71 | 0.80 | 0.81 | 0.81 | 0.70 | 1.00 | 0.82 | 0.78 |
| VC11 | 0.80 | 0.83 | 0.81 | 0.81 | 0.74 | 0.83 | 0.88 | 0.86 | 0.77 | 0.86 | 0.86 | 0.80 | 0.77 | 0.82 | 1.00 | 0.81 |

(B)

| Name | sc1 | sc2 | sc3 | sc4 | sc5 | sc6 | sc7 | sc8 | sc9 | tc1 | tc2 | tc3 | tc4 | tc5 | tc6 | tc7 | vc1 | vc2 | vc3 | vc4 | vc5 | vc6 | vc7 | vc8 | Average |
|---|---|---|---|---|---|---|---|---|---|---|---|---|---|---|---|---|---|---|---|---|---|---|---|---|---|
| sc1 | 1.00 | 0.49 | 0.64 | 0.54 | 0.66 | 0.66 | 0.62 | 0.70 | 0.57 | 0.67 | 0.67 | 0.69 | 0.70 | 0.66 | 0.62 | 0.65 | 0.59 | 0.62 | 0.41 | 0.72 | 0.66 | 0.67 | 0.66 | 0.63 | 0.63 |
| sc2 | 0.50 | 1.00 | 0.68 | 0.41 | 0.52 | 0.63 | 0.50 | 0.63 | 0.39 | 0.48 | 0.48 | 0.62 | 0.53 | 0.55 | 0.44 | 0.69 | 0.44 | 0.68 | 0.18 | 0.48 | 0.64 | 0.49 | 0.59 | 0.63 | 0.53 |
| sc3 | 0.65 | 0.69 | 1.00 | 0.47 | 0.66 | 0.71 | 0.58 | 0.77 | 0.47 | 0.57 | 0.59 | 0.73 | 0.60 | 0.68 | 0.58 | 0.76 | 0.54 | 0.76 | 0.38 | 0.59 | 0.71 | 0.59 | 0.67 | 0.73 | 0.63 |
| sc4 | 0.55 | 0.41 | 0.47 | 1.00 | 0.59 | 0.59 | 0.68 | 0.55 | 0.64 | 0.68 | 0.69 | 0.59 | 0.59 | 0.49 | 0.71 | 0.51 | 0.58 | 0.49 | 0.46 | 0.57 | 0.49 | 0.70 | 0.65 | 0.50 | 0.57 |
| sc5 | 0.67 | 0.52 | 0.66 | 0.60 | 1.00 | 0.75 | 0.63 | 0.72 | 0.62 | 0.67 | 0.74 | 0.72 | 0.65 | 0.69 | 0.71 | 0.72 | 0.56 | 0.65 | 0.27 | 0.62 | 0.66 | 0.71 | 0.72 | 0.73 | 0.65 |
| sc6 | 0.67 | 0.63 | 0.71 | 0.59 | 0.75 | 1.00 | 0.68 | 0.79 | 0.52 | 0.68 | 0.71 | 0.79 | 0.69 | 0.73 | 0.65 | 0.76 | 0.55 | 0.73 | 0.27 | 0.66 | 0.74 | 0.68 | 0.79 | 0.76 | 0.68 |
| sc7 | 0.63 | 0.50 | 0.57 | 0.67 | 0.62 | 0.66 | 1.00 | 0.65 | 0.59 | 0.72 | 0.71 | 0.68 | 0.70 | 0.60 | 0.69 | 0.61 | 0.61 | 0.59 | 0.38 | 0.66 | 0.60 | 0.72 | 0.72 | 0.59 | 0.63 |
| sc8 | 0.70 | 0.63 | 0.77 | 0.54 | 0.72 | 0.79 | 0.65 | 1.00 | 0.53 | 0.66 | 0.69 | 0.79 | 0.70 | 0.75 | 0.66 | 0.76 | 0.57 | 0.71 | 0.37 | 0.69 | 0.74 | 0.68 | 0.76 | 0.76 | 0.68 |
| sc9 | 0.57 | 0.53 | 0.46 | 0.64 | 0.62 | 0.51 | 0.59 | 0.53 | 1.00 | 0.66 | 0.66 | 0.54 | 0.57 | 0.48 | 0.68 | 0.48 | 0.57 | 0.45 | 0.39 | 0.57 | 0.45 | 0.66 | 0.56 | 0.51 | 0.55 |
| tc1 | 0.67 | 0.48 | 0.57 | 0.68 | 0.66 | 0.68 | 0.72 | 0.66 | 0.66 | 1.00 | 0.77 | 0.70 | 0.72 | 0.62 | 0.74 | 0.57 | 0.60 | 0.59 | 0.31 | 0.68 | 0.59 | 0.72 | 0.71 | 0.59 | 0.64 |
| tc2 | 0.68 | 0.48 | 0.59 | 0.69 | 0.74 | 0.71 | 0.72 | 0.69 | 0.66 | 0.77 | 1.00 | 0.72 | 0.72 | 0.65 | 0.77 | 0.63 | 0.59 | 0.62 | 0.43 | 0.67 | 0.64 | 0.78 | 0.75 | 0.64 | 0.67 |
| tc3 | 0.70 | 0.62 | 0.73 | 0.59 | 0.72 | 0.79 | 0.68 | 0.79 | 0.54 | 0.70 | 0.72 | 1.00 | 0.71 | 0.73 | 0.68 | 0.76 | 0.61 | 0.73 | 0.25 | 0.68 | 0.72 | 0.73 | 0.78 | 0.75 | 0.68 |
| tc4 | 0.71 | 0.53 | 0.60 | 0.60 | 0.65 | 0.69 | 0.70 | 0.70 | 0.57 | 0.72 | 0.72 | 0.71 | 1.00 | 0.68 | 0.67 | 0.63 | 0.59 | 0.63 | 0.12 | 0.74 | 0.66 | 0.73 | 0.69 | 0.63 | 0.64 |
| tc5 | 0.67 | 0.55 | 0.68 | 0.49 | 0.69 | 0.73 | 0.60 | 0.75 | 0.49 | 0.62 | 0.65 | 0.73 | 0.68 | 1.00 | 0.62 | 0.70 | 0.53 | 0.72 | 0.30 | 0.62 | 0.71 | 0.65 | 0.70 | 0.72 | 0.63 |
| tc6 | 0.63 | 0.44 | 0.59 | 0.71 | 0.72 | 0.65 | 0.70 | 0.66 | 0.68 | 0.74 | 0.77 | 0.68 | 0.67 | 0.61 | 1.00 | 0.62 | 0.41 | 0.56 | 0.45 | 0.64 | 0.58 | 0.80 | 0.71 | 0.62 | 0.64 |
| tc7 | 0.65 | 0.70 | 0.77 | 0.51 | 0.72 | 0.76 | 0.61 | 0.76 | 0.49 | 0.58 | 0.63 | 0.76 | 0.63 | 0.70 | 0.62 | 1.00 | 0.55 | 0.71 | 0.34 | 0.61 | 0.72 | 0.65 | 0.73 | 0.76 | 0.65 |
| vc1 | 0.58 | 0.43 | 0.53 | 0.57 | 0.55 | 0.53 | 0.62 | 0.56 | 0.56 | 0.59 | 0.58 | 0.59 | 0.58 | 0.51 | 0.56 | 0.54 | 1.00 | 0.50 | 0.45 | 0.59 | 0.50 | 0.63 | 0.58 | 0.52 | 0.55 |
| vc2 | 0.62 | 0.68 | 0.75 | 0.49 | 0.65 | 0.73 | 0.59 | 0.74 | 0.45 | 0.59 | 0.62 | 0.73 | 0.63 | 0.71 | 0.56 | 0.71 | 0.42 | 1.00 | 0.35 | 0.60 | 0.71 | 0.61 | 0.67 | 0.72 | 0.62 |
| vc3 | 0.40 | 0.36 | 0.34 | 0.41 | 0.38 | 0.06 | 0.35 | 0.35 | 0.38 | 0.06 | 0.05 | 0.29 | 0.26 | 0.29 | 0.20 | 0.18 | 0.40 | 0.32 | 1.00 | 0.40 | 0.31 | 0.38 | 0.30 | 0.32 | 0.29 |
| vc4 | 0.72 | 0.41 | 0.59 | 0.57 | 0.61 | 0.66 | 0.66 | 0.68 | 0.57 | 0.69 | 0.67 | 0.68 | 0.74 | 0.61 | 0.64 | 0.60 | 0.60 | 0.60 | 0.41 | 1.00 | 0.64 | 0.69 | 0.65 | 0.61 | 0.62 |
| vc5 | 0.67 | 0.64 | 0.70 | 0.49 | 0.66 | 0.74 | 0.61 | 0.74 | 0.46 | 0.59 | 0.64 | 0.72 | 0.65 | 0.70 | 0.58 | 0.71 | 0.52 | 0.71 | 0.22 | 0.64 | 1.00 | 0.61 | 0.68 | 0.72 | 0.63 |
| vc6 | 0.67 | 0.49 | 0.59 | 0.71 | 0.71 | 0.68 | 0.72 | 0.68 | 0.66 | 0.73 | 0.78 | 0.73 | 0.73 | 0.65 | 0.80 | 0.65 | 0.65 | 0.61 | 0.41 | 0.69 | 0.61 | 1.00 | 0.74 | 0.64 | 0.67 |
| vc7 | 0.30 | 0.59 | 0.67 | 0.65 | 0.72 | 0.78 | 0.72 | 0.76 | 0.56 | 0.71 | 0.75 | 0.78 | 0.69 | 0.70 | 0.70 | 0.73 | 0.59 | 0.68 | 0.22 | 0.65 | 0.68 | 0.74 | 1.00 | 0.71 | 0.66 |
| vc8 | 0.63 | 0.63 | 0.73 | 0.50 | 0.73 | 0.76 | 0.60 | 0.76 | 0.51 | 0.59 | 0.64 | 0.75 | 0.63 | 0.72 | 0.62 | 0.75 | 0.53 | 0.72 | 0.30 | 0.61 | 0.71 | 0.64 | 0.71 | 1.00 | 0.64 |

(C)

| Name | SC1 | SC2 | SC3 | SC4 | SC5 | SC6 | SC7 | SC8 | SC9 | SC10 | SC11 |
|---|---|---|---|---|---|---|---|---|---|---|---|
| L | 23,897 | 24,719 | 59,934 | 11,850 | 9,829 | 5,349 | 16,983 | 50,080 | 12,367 | 52,334 | 16,778 |
| R | 72,261 | 46,879 | 110,346 | 40,754 | 35,498 | 47,989 | 66,908 | 124,642 | 77,652 | 117,450 | 26,924 |

| Name | sc1 | sc2 | sc3 | sc4 | sc5 | sc6 | sc7 | sc8 | sc9 |
|---|---|---|---|---|---|---|---|---|---|
| L | 0 | 0 | 0 | 29,159 | 1,741 | 9,792 | 32,522 | 762 | 772 |
| R | 883 | 1,590 | 6,688 | 65,088 | 15,128 | 20,508 | 16,674 | 9,251 | 4,586 |

| Name | SC1 | SC2 | SC3 | SC4 | SC5 | SC6 | SC7 | SC8 | SC9 | SC10 | SC11 |
|---|---|---|---|---|---|---|---|---|---|---|---|
| L | 160,985 | 145,455 | 154,289 | 131,417 | 152,098 | 131,247 | 107,578 | 146,435 | 128,871 | 105,871 | 214,958 |
| R | 137,273 | 108,039 | 126,711 | 104,581 | 127,971 | 98,237 | 94,105 | 121,722 | 100,408 | 95,185 | 126,292 |

| Name | sc1 | sc2 | sc3 | sc4 | sc5 | sc6 | sc7 | sc8 | sc9 |
|---|---|---|---|---|---|---|---|---|---|
| L | 51,843 | 65,731 | 56,576 | 55,762 | 87,852 | 115,957 | 75,346 | 77,680 | 27,297 |
| R | 47,121 | 46,896 | 55,922 | 29,503 | 84,863 | 68,388 | 63,028 | 68,141 | 27,261 |

(D)

| Name | TC1 | TC2 | TC3 | TC4 | TC5 | TC6 | TC7 | TC8 | TC9 | TC10 |
|---|---|---|---|---|---|---|---|---|---|---|
| L | 2,347 | 17,351 | 1,196 | 4,016 | 11,903 | 6,296 | 11,556 | 4,102 | 17,623 | 5,242 |
| R | 45,894 | 34,072 | 24,069 | 44,659 | 37,443 | 40,364 | 36,765 | 51,817 | 79,855 | 45,723 |

| Name | tc1 | tc2 | tc3 | tc4 | tc5 | tc6 | tc7 |
|---|---|---|---|---|---|---|---|
| L | 0 | 658 | 899 | 0 | 1,101 | 0 | 0 |
| R | 834 | 22,951 | 13,065 | 2,575 | 15,373 | 27,954 | 778 |

| Name | TC1 | TC2 | TC3 | TC4 | TC5 | TC6 | TC7 | TC8 | TC9 | TC10 |
|---|---|---|---|---|---|---|---|---|---|---|
| L | 115,716 | 137,224 | 109,090 | 131,764 | 158,491 | 118,669 | 145,956 | 139,551 | 140,254 | 138,527 |
| R | 100,541 | 105,151 | 80,765 | 112,082 | 125,341 | 97,109 | 113,645 | 127,718 | 126,510 | 112,202 |

| Name | tc1 | tc2 | tc3 | tc4 | tc5 | tc6 | tc7 |
|---|---|---|---|---|---|---|---|
| L | 65,954 | 101,587 | 99,405 | 78,747 | 70,889 | 94,347 | 65,935 |
| R | 59,422 | 78,964 | 79,245 | 65,989 | 71,634 | 75,409 | 48,446 |

(E)

| Name | VC1 | VC2 | VC3 | VC4 | VC5 | VC6 | VC7 | VC8 | VC9 | VC10 | VC11 |
|---|---|---|---|---|---|---|---|---|---|---|---|
| L | 41,837 | 102,673 | 128,994 | 95,291 | 102,761 | 110,513 | 105,165 | 76,173 | 95,140 | 82,202 | 87,803 |
| R | 24,635 | 77,703 | 90,094 | 69,625 | 67,388 | 60,826 | 59,184 | 40,343 | 46,535 | 62,499 | 44,131 |

| Name | vc1 | vc2 | vc3 | vc4 | vc5 | vc6 | vc7 | vc8 |
|---|---|---|---|---|---|---|---|---|
| L | 1,740 | 9,701 | 4,212 | 27,404 | 13,150 | 25,477 | 35,333 | 19,278 |
| R | 1,932 | 8,114 | 2,900 | 33,110 | 13,010 | 31,299 | 5,959 | 17,767 |

| Name | VC1 | VC2 | VC3 | VC4 | VC5 | VC6 | VC7 | VC8 | VC9 | VC10 | VC11 |
|---|---|---|---|---|---|---|---|---|---|---|---|
| L | 112,565 | 153,639 | 168,656 | 171,527 | 181,903 | 156,486 | 153,101 | 99,551 | 148,902 | 97,893 | 177,789 |
| R | 87,125 | 130,311 | 124,143 | 132,614 | 163,085 | 121,091 | 118,443 | 84,325 | 112,850 | 89,350 | 134,529 |

| Name | vc1 | vc2 | vc3 | vc4 | vc5 | vc6 | vc7 | vc8 |
|---|---|---|---|---|---|---|---|---|
| L | 30,204 | 61,410 | 8,991 | 55,413 | 67,352 | 91,096 | 79,972 | 58,655 |
| R | 29,804 | 59,871 | 9,612 | 51,012 | 54,664 | 85,812 | 51,849 | 53,928 |

(F)

| | WT L/R % | wls L/R % |
|---|---|---|
| cpd2 | 124.6 | 120.2 |
| slc18a3b | 37.0 | 53.2 |
| tac3a | 18.5 | 3.2 |
| vglut2 | 159.9 | 119.5 |

**Supplemental Table 2. Computed correlation factors from reciprocal one-to-one registration for each Z-stacks of *gng8*.** (A-D) Correlation factors (CFs) from 72h (A-B) or 96h (C-D) wildtype (WT in A, C) or *wls* mutant (mt in B, D) groups. (E-H) Volumes of *gng8* (HA marker) and *clu* (choroid plexus marker) expression domain (vol., $\mu m^3$) from samples of WT (E, G)or *wls* (wls in F, H) groups. Intensity thresholds "I" is 80 (E, F) or 40 (G, H) for *gng8*, and is 50 (E, F) or 12 (G, H) for *clu* expression vol. calculation.

(A)

| Name | WT1 | WT2 | WT3 | WT4 | WT5 | WT6 | WT7 | WT8 | WT9 | WT10 | WT11 | WT12 | WT13 | WT14 | Average |
|---|---|---|---|---|---|---|---|---|---|---|---|---|---|---|---|
| WT1 | 1.00 | 0.90 | 0.82 | 0.83 | 0.84 | 0.90 | 0.88 | 0.82 | 0.86 | 0.84 | 0.88 | 0.86 | 0.87 | 0.86 | 0.86 |
| WT2 | 0.89 | 1.00 | 0.84 | 0.85 | 0.84 | 0.89 | 0.89 | 0.84 | 0.88 | 0.88 | 0.89 | 0.89 | 0.88 | 0.89 | 0.87 |
| WT3 | 0.55 | 0.80 | 1.00 | 0.85 | 0.83 | 0.78 | 0.80 | 0.89 | 0.84 | 0.46 | 0.79 | 0.80 | 0.34 | 0.77 | 0.73 |
| WT4 | 0.61 | 0.61 | 0.79 | 1.00 | 0.73 | 0.80 | 0.77 | 0.79 | 0.59 | 0.40 | 0.75 | 0.78 | 0.36 | 0.07 | 0.62 |
| WT5 | 0.82 | 0.82 | 0.84 | 0.81 | 1.00 | 0.83 | 0.84 | 0.85 | 0.87 | 0.79 | 0.81 | 0.85 | 0.56 | 0.80 | 0.81 |
| WT6 | 0.86 | 0.86 | 0.81 | 0.83 | 0.82 | 1.00 | 0.84 | 0.81 | 0.84 | 0.80 | 0.82 | 0.86 | 0.62 | 0.82 | 0.81 |
| WT7 | 0.85 | 0.87 | 0.83 | 0.83 | 0.84 | 0.85 | 1.00 | 0.84 | 0.89 | 0.85 | 0.87 | 0.89 | 0.84 | 0.86 | 0.85 |
| WT8 | 0.76 | 0.78 | 0.87 | 0.85 | 0.83 | 0.78 | 0.80 | 1.00 | 0.81 | 0.40 | 0.76 | 0.79 | 0.21 | 0.43 | 0.70 |
| WT9 | 0.81 | 0.84 | 0.84 | 0.83 | 0.84 | 0.82 | 0.87 | 0.83 | 1.00 | 0.83 | 0.84 | 0.86 | 0.82 | 0.84 | 0.84 |
| WT10 | 0.83 | 0.87 | 0.81 | 0.81 | 0.82 | 0.82 | 0.88 | 0.82 | 0.87 | 1.00 | 0.87 | 0.86 | 0.89 | 0.90 | 0.85 |
| WT11 | 0.87 | 0.88 | 0.82 | 0.82 | 0.81 | 0.84 | 0.88 | 0.82 | 0.86 | 0.86 | 1.00 | 0.86 | 0.88 | 0.86 | 0.85 |
| WT12 | 0.82 | 0.86 | 0.83 | 0.83 | 0.84 | 0.85 | 0.88 | 0.83 | 0.87 | 0.84 | 0.83 | 1.00 | 0.84 | 0.86 | 0.84 |
| WT13 | 0.88 | 0.90 | 0.83 | 0.64 | 0.83 | 0.87 | 0.88 | 0.15 | 0.88 | 0.90 | 0.90 | 0.88 | 1.00 | 0.91 | 0.80 |
| WT14 | 0.85 | 0.89 | 0.82 | 0.83 | 0.83 | 0.85 | 0.88 | 0.83 | 0.88 | 0.91 | 0.87 | 0.88 | 0.90 | 1.00 | 0.86 |

(B)

| Name | mt1 | mt2 | mt3 | mt4 | mt5 | mt6 | mt7 | mt8 | Average |
|---|---|---|---|---|---|---|---|---|---|
| mt1 | 1.00 | 0.81 | 0.73 | 0.79 | 0.84 | 0.80 | 0.63 | 0.70 | 0.76 |
| mt2 | 0.80 | 1.00 | 0.71 | 0.78 | 0.80 | 0.80 | 0.72 | 0.70 | 0.76 |
| mt3 | 0.75 | 0.79 | 1.00 | 0.79 | 0.80 | 0.78 | 0.68 | 0.61 | 0.74 |
| mt4 | 0.73 | 0.74 | 0.69 | 1.00 | 0.77 | 0.77 | 0.72 | 0.54 | 0.71 |
| mt5 | 0.83 | 0.81 | 0.77 | 0.82 | 1.00 | 0.81 | 0.66 | 0.66 | 0.77 |
| mt6 | 0.78 | 0.79 | 0.70 | 0.79 | 0.80 | 1.00 | 0.73 | 0.60 | 0.74 |
| mt7 | 0.52 | 0.65 | 0.54 | 0.71 | 0.55 | 0.70 | 1.00 | 0.39 | 0.58 |
| mt8 | 0.74 | 0.77 | 0.67 | 0.40 | 0.73 | 0.60 | 0.49 | 1.00 | 0.63 |

(C)

| Name | WT1 | WT2 | WT3 | WT4 | WT5 | WT6 | WT7 | WT8 | WT9 | WT10 | Average |
|---|---|---|---|---|---|---|---|---|---|---|---|
| WT1 | 1.00 | 0.66 | 0.70 | 0.71 | 0.71 | 0.72 | 0.65 | 0.69 | 0.71 | 0.71 | 0.70 |
| WT2 | 0.67 | 1.00 | 0.75 | 0.74 | 0.71 | 0.77 | 0.64 | 0.65 | 0.77 | 0.74 | 0.72 |
| WT3 | 0.70 | 0.75 | 1.00 | 0.77 | 0.78 | 0.80 | 0.70 | 0.77 | 0.82 | 0.77 | 0.76 |
| WT4 | 0.71 | 0.74 | 0.77 | 1.00 | 0.79 | 0.80 | 0.72 | 0.78 | 0.81 | 0.79 | 0.77 |
| WT5 | 0.68 | 0.70 | 0.77 | 0.77 | 1.00 | 0.77 | 0.69 | 0.76 | 0.77 | 0.75 | 0.74 |
| WT6 | 0.75 | 0.80 | 0.83 | 0.83 | 0.82 | 1.00 | 0.75 | 0.80 | 0.84 | 0.84 | 0.81 |
| WT7 | 0.73 | 0.68 | 0.79 | 0.80 | 0.81 | 0.79 | 1.00 | 0.81 | 0.80 | 0.80 | 0.78 |
| WT8 | 0.68 | 0.68 | 0.77 | 0.76 | 0.77 | 0.76 | 0.69 | 1.00 | 0.77 | 0.72 | 0.73 |
| WT9 | 0.73 | 0.78 | 0.83 | 0.83 | 0.81 | 0.83 | 0.73 | 0.79 | 1.00 | 0.81 | 0.79 |
| WT10 | 0.77 | 0.79 | 0.83 | 0.83 | 0.83 | 0.86 | 0.79 | 0.79 | 0.85 | 1.00 | 0.81 |

(D)

| Name | mt1 | mt2 | mt3 | mt4 | mt5 | mt6 | mt7 | mt8 | Average |
|---|---|---|---|---|---|---|---|---|---|
| mt1 | 1.00 | 0.52 | 0.63 | 0.73 | 0.40 | 0.46 | 0.62 | 0.70 | 0.58 |
| mt2 | 0.39 | 1.00 | 0.43 | 0.53 | 0.66 | 0.65 | 0.69 | 0.57 | 0.56 |
| mt3 | 0.54 | 0.47 | 1.00 | 0.59 | 0.60 | 0.61 | 0.59 | 0.64 | 0.58 |
| mt4 | 0.71 | 0.63 | 0.67 | 1.00 | 0.69 | 0.70 | 0.69 | 0.78 | 0.69 |
| mt5 | 0.22 | 0.62 | 0.52 | 0.39 | 1.00 | 0.70 | 0.75 | 0.55 | 0.54 |
| mt6 | 0.35 | 0.65 | 0.51 | 0.57 | 0.74 | 1.00 | 0.75 | 0.60 | 0.59 |
| mt7 | 0.40 | 0.63 | 0.47 | 0.49 | 0.71 | 0.67 | 1.00 | 0.54 | 0.56 |
| mt8 | 0.61 | 0.64 | 0.63 | 0.71 | 0.67 | 0.67 | 0.67 | 1.00 | 0.66 |

(E)

| Name | WT1 | WT2 | WT3 | WT4 | WT5 | WT6 | WT7 |
|---|---|---|---|---|---|---|---|
| CP | 8,760 | 16,337 | 2,882 | 6,871 | 7,084 | 3,209 | 4,270 |
| L | 135,025 | 142,996 | 134,903 | 119,334 | 109,794 | 105,918 | 147,203 |
| R | 87,190 | 83,045 | 103,064 | 67,291 | 90,122 | 65,931 | 104,227 |

| Name | WT8 | WT9 | WT10 | WT11 | WT12 | WT13 | WT14 |
|---|---|---|---|---|---|---|---|
| CP | 1,875 | 4,427 | 10,517 | 6,469 | 3,558 | 11,925 | 4,365 |
| L | 121,337 | 128,809 | 143,973 | 165,715 | 123,850 | 150,445 | 127,740 |
| R | 84,847 | 97,763 | 112,496 | 137,936 | 78,955 | 90,011 | 83,717 |

(F)

| Name | wls1 | wls2 | wls3 | wls4 | wls5 | wls6 | wls7 | wls8 |
|---|---|---|---|---|---|---|---|---|
| CP | 12,315 | 10,560 | 19,065 | 8,829 | 15,278 | 18,049 | 22,217 | 18,406 |
| L | 61,211 | 104,965 | 67,655 | 60,998 | 71,998 | 54,824 | 30,981 | 118,907 |
| R | 48,163 | 72,589 | 61,946 | 41,169 | 49,278 | 36,154 | 37,880 | 85,987 |

(G)

| Name | WT1 | WT2 | WT3 | WT4 | WT5 | WT6 | WT7 | WT8 | WT9 | WT10 |
|---|---|---|---|---|---|---|---|---|---|---|
| CP | 4,109 | 4,135 | 7,900 | 6,091 | 1,858 | 2,299 | 2,071 | 1,703 | 3,778 | 3,126 |
| L | 95,769 | 137,631 | 131,535 | 171,113 | 189,782 | 222,647 | 148,953 | 157,113 | 179,009 | 236,820 |
| R | 48,726 | 66,822 | 81,539 | 133,664 | 112,626 | 144,761 | 72,292 | 100,671 | 113,917 | 139,825 |

(H)

| Name | wls1 | wls2 | wls3 | wls4* | wls5 | wls6 | wls7 | wls8 | wls9 |
|---|---|---|---|---|---|---|---|---|---|
| CP | 48,788 | 4,187 | 16,193 | 9,959 | 25,000 | 6,113 | 7,838 | 8,408 | 21,336 |
| L | 21,096 | 125,189 | 14,677 | 84,497 | 83,013 | 89,354 | 113,537 | 89,212 | 141,797 |
| R | 6,150 | 65,240 | 16,060 | 149,222 | 36,216 | 68,132 | 71,081 | 91,641 | 88,221 |

**Supplemental Table 3. Computed correlation factors from reciprocal one-to-one registration foreach Z-stacks of *ngn1*.** (A-B) Correlation factors (CFs) for wildtype (A, NG1 to NG 21) or *wls* mutant (B, ng1 to ng14) groups. (C-D)Volumes of*ngn1* expression domain (µm$^3$) from samples of wildtype (C)or *wls* mutant (D) groups.L is left, R is right and M is middle domain.Intensity threshold (I) = 15 for vol. calculation.

(A)

| Name | NG1 | NG2 | NG3 | NG4 | NG5 | NG6 | NG7 | NG8 | NG9 | NG10 | NG11 | NG12 | NG13 | NG14 | NG15 | NG16 | NG17 | NG18 | NG19 | NG20 | NG21 | Average |
|---|---|---|---|---|---|---|---|---|---|---|---|---|---|---|---|---|---|---|---|---|---|---|
| NG1 | 1.00 | 0.33 | 0.29 | 0.25 | 0.27 | 0.25 | 0.29 | 0.28 | 0.30 | 0.29 | 0.27 | 0.30 | 0.28 | 0.27 | 0.27 | 0.24 | 0.24 | 0.24 | 0.27 | 0.30 | 0.26 | 0.27 |
| NG2 | 0.32 | 1.00 | 0.29 | 0.30 | 0.31 | 0.30 | 0.31 | 0.32 | 0.32 | 0.34 | 0.32 | 0.33 | 0.32 | 0.25 | 0.27 | 0.32 | 0.26 | 0.32 | 0.29 | 0.29 | 0.28 | 0.30 |
| NG3 | 0.27 | 0.30 | 1.00 | 0.37 | 0.40 | 0.35 | 0.36 | 0.44 | 0.37 | 0.41 | 0.40 | 0.38 | 0.34 | 0.35 | 0.28 | 0.38 | 0.28 | 0.37 | 0.31 | 0.35 | 0.29 | 0.35 |
| NG4 | 0.26 | 0.29 | 0.37 | 1.00 | 0.38 | 0.36 | 0.40 | 0.43 | 0.40 | 0.44 | 0.45 | 0.41 | 0.39 | 0.39 | 0.37 | 0.36 | 0.30 | 0.41 | 0.37 | 0.32 | 0.29 | 0.37 |
| NG5 | 0.29 | 0.31 | 0.41 | 0.39 | 1.00 | 0.40 | 0.41 | 0.49 | 0.41 | 0.44 | 0.44 | 0.41 | 0.34 | 0.37 | 0.29 | 0.38 | 0.29 | 0.41 | 0.34 | 0.35 | 0.31 | 0.37 |
| NG6 | 0.26 | 0.29 | 0.34 | 0.37 | 0.39 | 1.00 | 0.35 | 0.43 | 0.39 | 0.40 | 0.38 | 0.37 | 0.32 | 0.33 | 0.34 | 0.34 | 0.29 | 0.40 | 0.31 | 0.39 | 0.32 | 0.35 |
| NG7 | 0.30 | 0.32 | 0.37 | 0.38 | 0.39 | 0.36 | 1.00 | 0.42 | 0.39 | 0.43 | 0.35 | 0.39 | 0.36 | 0.33 | 0.35 | 0.38 | 0.30 | 0.35 | 0.34 | 0.36 | 0.28 | 0.36 |
| NG8 | 0.28 | 0.30 | 0.43 | 0.43 | 0.48 | 0.42 | 0.41 | 1.00 | 0.42 | 0.46 | 0.44 | 0.46 | 0.36 | 0.37 | 0.33 | 0.39 | 0.28 | 0.44 | 0.37 | 0.38 | 0.31 | 0.39 |
| NG9 | 0.25 | 0.31 | 0.36 | 0.40 | 0.39 | 0.39 | 0.38 | 0.42 | 1.00 | 0.41 | 0.40 | 0.43 | 0.35 | 0.34 | 0.33 | 0.35 | 0.28 | 0.39 | 0.32 | 0.35 | 0.30 | 0.36 |
| NG10 | 0.28 | 0.34 | 0.41 | 0.45 | 0.43 | 0.40 | 0.43 | 0.46 | 0.42 | 1.00 | 0.42 | 0.43 | 0.38 | 0.37 | 0.32 | 0.42 | 0.31 | 0.44 | 0.39 | 0.37 | 0.31 | 0.39 |
| NG11 | 0.27 | 0.31 | 0.40 | 0.45 | 0.43 | 0.38 | 0.37 | 0.44 | 0.40 | 0.42 | 1.00 | 0.42 | 0.37 | 0.39 | 0.36 | 0.38 | 0.32 | 0.44 | 0.33 | 0.25 | 0.30 | 0.37 |
| NG12 | 0.31 | 0.35 | 0.40 | 0.43 | 0.43 | 0.39 | 0.41 | 0.48 | 0.45 | 0.45 | 0.43 | 1.00 | 0.39 | 0.41 | 0.38 | 0.40 | 0.35 | 0.47 | 0.41 | 0.38 | 0.34 | 0.40 |
| NG13 | 0.28 | 0.32 | 0.35 | 0.41 | 0.34 | 0.33 | 0.37 | 0.37 | 0.36 | 0.38 | 0.38 | 0.39 | 1.00 | 0.40 | 0.33 | 0.39 | 0.32 | 0.42 | 0.37 | 0.36 | 0.37 | 0.36 |
| NG14 | 0.24 | 0.31 | 0.35 | 0.40 | 0.37 | 0.34 | 0.34 | 0.38 | 0.35 | 0.37 | 0.40 | 0.40 | 0.40 | 1.00 | 0.33 | 0.38 | 0.32 | 0.42 | 0.37 | 0.35 | 0.33 | 0.36 |
| NG15 | 0.24 | 0.28 | 0.29 | 0.38 | 0.30 | 0.35 | 0.34 | 0.34 | 0.35 | 0.33 | 0.37 | 0.37 | 0.33 | 0.34 | 1.00 | 0.32 | 0.34 | 0.39 | 0.34 | 0.35 | 0.31 | 0.33 |
| NG16 | 0.30 | 0.31 | 0.38 | 0.36 | 0.37 | 0.34 | 0.37 | 0.39 | 0.36 | 0.42 | 0.38 | 0.39 | 0.38 | 0.38 | 0.31 | 1.00 | 0.29 | 0.39 | 0.35 | 0.28 | 0.34 | 0.35 |
| NG17 | 0.26 | 0.26 | 0.29 | 0.31 | 0.31 | 0.29 | 0.33 | 0.32 | 0.29 | 0.30 | 0.33 | 0.34 | 0.33 | 0.32 | 0.33 | 0.30 | 1.00 | 0.36 | 0.31 | 0.33 | 0.31 | 0.31 |
| NG18 | 0.29 | 0.33 | 0.39 | 0.43 | 0.41 | 0.41 | 0.37 | 0.45 | 0.40 | 0.45 | 0.45 | 0.47 | 0.42 | 0.42 | 0.39 | 0.39 | 0.36 | 1.00 | 0.42 | 0.37 | 0.34 | 0.40 |
| NG19 | 0.26 | 0.30 | 0.32 | 0.38 | 0.34 | 0.32 | 0.34 | 0.38 | 0.33 | 0.40 | 0.34 | 0.40 | 0.36 | 0.37 | 0.34 | 0.35 | 0.32 | 0.42 | 1.00 | 0.31 | 0.31 | 0.34 |
| NG20 | 0.28 | 0.30 | 0.36 | 0.34 | 0.36 | 0.40 | 0.37 | 0.40 | 0.37 | 0.37 | 0.27 | 0.38 | 0.24 | 0.36 | 0.35 | 0.36 | 0.34 | 0.38 | 0.31 | 1.00 | 0.29 | 0.34 |
| NG21 | 0.27 | 0.29 | 0.30 | 0.31 | 0.32 | 0.33 | 0.30 | 0.33 | 0.32 | 0.32 | 0.32 | 0.34 | 0.38 | 0.34 | 0.31 | 0.35 | 0.32 | 0.34 | 0.31 | 0.29 | 1.00 | 0.32 |

(B)

| Name | ng1 | ng2 | ng3 | ng4 | ng5 | ng6 | ng7 | ng8 | ng9 | ng10 | ng11 | ng12 | ng13 | ng14 | Average |
|---|---|---|---|---|---|---|---|---|---|---|---|---|---|---|---|
| ng1 | 1.00 | 0.33 | 0.29 | 0.28 | 0.32 | 0.26 | 0.44 | 0.27 | 0.30 | 0.32 | 0.31 | 0.35 | 0.51 | 0.33 | 0.33 |
| ng2 | 0.33 | 1.00 | 0.29 | 0.28 | 0.30 | 0.23 | 0.25 | 0.28 | 0.29 | 0.31 | 0.36 | 0.34 | 0.36 | 0.35 | 0.30 |
| ng3 | 0.28 | 0.29 | 1.00 | 0.25 | 0.32 | 0.22 | 0.14 | 0.25 | 0.28 | 0.28 | 0.25 | 0.30 | 0.27 | 0.25 | 0.26 |
| ng4 | 0.23 | 0.26 | 0.25 | 1.00 | 0.32 | 0.29 | 0.17 | 0.25 | 0.23 | 0.27 | 0.24 | 0.29 | 0.26 | 0.22 | 0.25 |
| ng5 | 0.30 | 0.29 | 0.31 | 0.32 | 1.00 | 0.32 | 0.24 | 0.25 | 0.27 | 0.30 | 0.28 | 0.32 | 0.29 | 0.29 | 0.29 |
| ng6 | 0.25 | 0.22 | 0.23 | 0.30 | 0.32 | 1.00 | 0.24 | 0.25 | 0.25 | 0.28 | 0.26 | 0.31 | 0.24 | 0.16 | 0.25 |
| ng7 | 0.43 | 0.33 | 0.15 | 0.20 | 0.24 | 0.21 | 1.00 | 0.21 | 0.24 | 0.35 | 0.36 | 0.42 | 0.54 | 0.38 | 0.31 |
| ng8 | 0.25 | 0.27 | 0.26 | 0.26 | 0.24 | 0.25 | 0.22 | 1.00 | 0.26 | 0.28 | 0.30 | 0.32 | 0.23 | 0.19 | 0.26 |
| ng9 | 0.29 | 0.29 | 0.28 | 0.25 | 0.28 | 0.25 | 0.22 | 0.26 | 1.00 | 0.33 | 0.28 | 0.33 | 0.27 | 0.29 | 0.28 |
| ng10 | 0.32 | 0.31 | 0.28 | 0.28 | 0.23 | 0.28 | 0.35 | 0.29 | 0.33 | 1.00 | 0.35 | 0.41 | 0.36 | 0.33 | 0.32 |
| ng11 | 0.31 | 0.36 | 0.25 | 0.25 | 0.27 | 0.26 | 0.35 | 0.30 | 0.20 | 0.32 | 1.00 | 0.39 | 0.34 | 0.38 | 0.31 |
| ng12 | 0.35 | 0.34 | 0.29 | 0.31 | 0.32 | 0.32 | 0.41 | 0.32 | 0.32 | 0.40 | 0.39 | 1.00 | 0.37 | 0.40 | 0.35 |
| ng13 | 0.50 | 0.35 | 0.28 | 0.27 | 0.29 | 0.25 | 0.53 | 0.23 | 0.26 | 0.36 | 0.34 | 0.37 | 1.00 | 0.35 | 0.34 |
| ng14 | 0.33 | 0.34 | 0.25 | 0.25 | 0.28 | 0.16 | 0.37 | 0.18 | 0.29 | 0.33 | 0.37 | 0.40 | 0.35 | 1.00 | 0.30 |

(C)

| Name | NG1 | NG2 | NG3 | NG4 | NG5 | NG6 | NG7 | NG8 | NG9 | NG10 | NG11 |
|---|---|---|---|---|---|---|---|---|---|---|---|
| L | 47,927 | 49,036 | 35,400 | 62,397 | 41,468 | 73,293 | 57,276 | 65,918 | 36,103 | 76,355 | 94,292 |
| R | 53,034 | 51,295 | 42,703 | 58,157 | 59,112 | 85,280 | 73,378 | 75,464 | 41,231 | 86,785 | 88,125 |
| Middle | 61,861 | 80,736 | 38,636 | 76,948 | 44,664 | 142,000 | 55,712 | 72,802 | 42,630 | 107,933 | 109,019 |

| Name | NG12 | NG13 | NG14 | NG15 | NG16 | NG17 | NG18 | NG19 | NG20 | NG21 |
|---|---|---|---|---|---|---|---|---|---|---|
| L | 70,153 | 91,045 | 71,937 | 71,569 | 81,013 | 49,118 | 93,243 | 61,289 | 33,691 | 45,006 |
| R | 80,155 | 88,046 | 76,366 | 57,470 | 100,252 | 50,333 | 105,152 | 48,581 | 88,972 | 41,118 |
| Middle | 121,886 | 133,177 | 136,405 | 92,216 | 125,918 | 53,657 | 169,017 | 74,843 | 159,012 | 104,056 |

(D)

| Name | ng1 | ng2 | ng3 | ng4 | ng5 | ng6 | ng7 |
|---|---|---|---|---|---|---|---|
| L | 28,545 | 14,965 | 44,206 | 39,435 | 20,556 | 27,039 | 36,434 |
| R | 45,311 | 12,932 | 34,124 | 21,065 | 19,140 | 50,276 | 32,442 |
| Middle | 157,320 | 34,149 | 53,557 | 66,675 | 38,099 | 89,709 | 179,438 |

| Name | ng8 | ng9 | ng10 | ng11 | ng12 | ng13 | ng14 |
|---|---|---|---|---|---|---|---|
| L | 50,240 | 21,805 | 30,001 | 20,449 | 55,312 | 55,537 | 22,358 |
| R | 35,661 | 30,455 | 35,755 | 22,465 | 67,093 | 50,575 | 23,516 |
| Middle | 116,298 | 51,275 | 102,591 | 75,847 | 199,099 | 279,891 | 88,002 |

**Supplemental Table 4.Volumes of *ngn1*:GFP expressionsin habenula and their left-to-right (L/R) ratios.** (A-D) Volumes (vol. unit = μm$^3$) of *ngn1*:*GFP*expressing domains inhebanulaprogenitor zone inwildtype (WT, in A and C) or *wls* mutant (mt, in B and D) larva at 72 hours(h) (A-B) or 96h (C-D) post-fertilization. L is left, R is right habenula. Intensity threshold (I) = 20 for vol. calculation.

(A)

| Name | WT1 | WT2 | WT3 | WT4 | WT5 | WT6 |
|---|---|---|---|---|---|---|
| L | 48,739 | 48,847 | 47,768 | 30,735 | 42,791 | 35,161 |
| R | 26,652 | 34,341 | 21,227 | 19,077 | 31,652 | 20,473 |
| L/R % | 183 | 142 | 225 | 161 | 135 | 172 |

| Name | WT7 | WT8 | WT9 | WT10 | WT11 | WT12 |
|---|---|---|---|---|---|---|
| L | 71,110 | 25,882 | 18,709 | 46,782 | 101,721 | 39,887 |
| R | 42,070 | 7,406 | 14,470 | 47,384 | 53,932 | 32,922 |
| L/R % | 169 | 349 | 129 | 99 | 189 | 121 |

(B)

| Name | mt1 | mt2 | mt3 | mt4 | mt5 | mt6 | mt7 |
|---|---|---|---|---|---|---|---|
| L | 6,050 | 24,640 | 15,904 | 7,700 | 56,954 | 7,286 | 62,879 |
| R | 3,041 | 31,486 | 9,770 | 11,740 | 39,668 | 3,205 | 29,221 |
| L/R % | 199 | 78 | 163 | 66 | 144 | 227 | 215 |

(C)

| Name | WT1 | WT2 | WT3 | WT4 | WT5 | WT6 |
|---|---|---|---|---|---|---|
| L | 87,116 | 62,680 | 46,731 | 41,791 | 28,766 | 27,106 |
| R | 46,547 | 41,608 | 22,864 | 22,662 | 26,340 | 15,646 |
| L/R % | 187 | 151 | 204 | 184 | 109 | 173 |

| Name | WT7 | WT8 | WT9 | WT10 | WT11 | WT12 |
|---|---|---|---|---|---|---|
| L | 83,080 | 35,191 | 33,332 | 51,669 | 98,887 | 49,878 |
| R | 44,152 | 13,205 | 12,649 | 48,096 | 36,907 | 19,113 |
| L/R % | 188 | 267 | 264 | 107 | 268 | 261 |

(D)

| Name | mt1 | mt2 | mt3 | mt4 | mt5 | mt6 | mt7 |
|---|---|---|---|---|---|---|---|
| L | 6,626 | 23,509 | 18,689 | 7,451 | 53,360 | 20,593 | 41,094 |
| R | 1,086 | 11,906 | 1,012 | 6,974 | 23,881 | 4,867 | 39,222 |
| L/R % | 610 | 197 | 1,846 | 107 | 223 | 423 | 105 |

**Supplemental Table 5. Computed correlation factors from reciprocal one-to-one registration foreach Z-stacks of *her6*.** (A-B) Correlation factors (CFs) for wildtype (A, H1 to H17) or *wls* mutant (B, h1 to h16) groups. (C-D)Volumes (Vol.) of *her6* expression domains ($\mu m^3$)from samples of wildtype (C)or *wls* mutant (D) groups.L is left, R is right and M= is middle domain. Intensity threshold (I) = 15 for vol. calculation.

(A)

| Name | H1 | H2 | H3 | H4 | H5 | H6 | H7 | H8 | H9 | H10 | H11 | H12 | H13 | H14 | H15 | H16 | H17 | Average |
|---|---|---|---|---|---|---|---|---|---|---|---|---|---|---|---|---|---|---|
| H1 | 1.00 | 0.32 | 0.30 | 0.33 | 0.32 | 0.35 | 0.26 | 0.30 | 0.25 | 0.28 | 0.26 | 0.27 | 0.26 | 0.28 | 0.27 | 0.28 | 0.26 | 0.29 |
| H2 | 0.32 | 1.00 | 0.33 | 0.35 | 0.32 | 0.37 | 0.34 | 0.35 | 0.28 | 0.28 | 0.32 | 0.30 | 0.30 | 0.33 | 0.29 | 0.36 | 0.34 | 0.32 |
| H3 | 0.30 | 0.35 | 1.00 | 0.35 | 0.35 | 0.37 | 0.30 | 0.33 | 0.28 | 0.17 | 0.28 | 0.25 | 0.28 | 0.27 | 0.29 | 0.31 | 0.34 | 0.30 |
| H4 | 0.32 | 0.36 | 0.33 | 1.00 | 0.33 | 0.40 | 0.35 | 0.31 | 0.28 | 0.28 | 0.28 | 0.26 | 0.25 | 0.30 | 0.27 | 0.31 | 0.34 | 0.31 |
| H5 | 0.32 | 0.34 | 0.34 | 0.34 | 1.00 | 0.36 | 0.28 | 0.32 | 0.27 | 0.28 | 0.28 | 0.28 | 0.27 | 0.27 | 0.29 | 0.31 | 0.31 | 0.30 |
| H6 | 0.34 | 0.37 | 0.34 | 0.40 | 0.34 | 1.00 | 0.34 | 0.35 | 0.28 | 0.27 | 0.28 | 0.27 | 0.27 | 0.34 | 0.28 | 0.34 | 0.34 | 0.32 |
| H7 | 0.27 | 0.35 | 0.29 | 0.35 | 0.27 | 0.34 | 1.00 | 0.32 | 0.24 | 0.23 | 0.27 | 0.25 | 0.24 | 0.32 | 0.24 | 0.33 | 0.35 | 0.29 |
| H8 | 0.29 | 0.36 | 0.30 | 0.30 | 0.32 | 0.35 | 0.33 | 1.00 | 0.30 | 0.30 | 0.32 | 0.31 | 0.30 | 0.34 | 0.32 | 0.36 | 0.36 | 0.32 |
| H9 | 0.24 | 0.26 | 0.18 | 0.26 | 0.24 | 0.28 | 0.22 | 0.28 | 1.00 | 0.32 | 0.31 | 0.29 | 0.30 | 0.30 | 0.33 | 0.31 | 0.30 | 0.28 |
| H10 | 0.27 | 0.27 | 0.14 | 0.27 | 0.27 | 0.27 | 0.22 | 0.30 | 0.33 | 1.00 | 0.29 | 0.31 | 0.33 | 0.28 | 0.34 | 0.29 | 0.28 | 0.28 |
| H11 | 0.26 | 0.34 | 0.25 | 0.29 | 0.28 | 0.29 | 0.28 | 0.33 | 0.35 | 0.31 | 1.00 | 0.38 | 0.37 | 0.34 | 0.36 | 0.39 | 0.32 | 0.32 |
| H12 | 0.28 | 0.31 | 0.25 | 0.27 | 0.28 | 0.28 | 0.25 | 0.32 | 0.32 | 0.32 | 0.36 | 1.00 | 0.37 | 0.31 | 0.36 | 0.33 | 0.30 | 0.31 |
| H13 | 0.27 | 0.32 | 0.27 | 0.27 | 0.27 | 0.29 | 0.25 | 0.31 | 0.34 | 0.35 | 0.37 | 0.38 | 1.00 | 0.31 | 0.38 | 0.33 | 0.29 | 0.31 |
| H14 | 0.27 | 0.33 | 0.23 | 0.29 | 0.26 | 0.34 | 0.32 | 0.34 | 0.32 | 0.29 | 0.32 | 0.31 | 0.29 | 1.00 | 0.29 | 0.37 | 0.36 | 0.31 |
| H15 | 0.26 | 0.30 | 0.27 | 0.27 | 0.28 | 0.29 | 0.24 | 0.32 | 0.34 | 0.35 | 0.34 | 0.34 | 0.36 | 0.29 | 1.00 | 0.35 | 0.30 | 0.31 |
| H16 | 0.28 | 0.37 | 0.28 | 0.33 | 0.31 | 0.35 | 0.33 | 0.36 | 0.35 | 0.30 | 0.37 | 0.33 | 0.33 | 0.38 | 0.36 | 1.00 | 0.39 | 0.34 |
| H17 | 0.25 | 0.34 | 0.31 | 0.33 | 0.31 | 0.36 | 0.35 | 0.37 | 0.31 | 0.28 | 0.30 | 0.29 | 0.28 | 0.36 | 0.30 | 0.38 | 1.00 | 0.32 |

(B)

| Name | h1 | h2 | h3 | h4 | h5 | h6 | h7 | h8 | h9 | h10 | h11 | h12 | h13 | h14 | h15 | h16 | Average |
|---|---|---|---|---|---|---|---|---|---|---|---|---|---|---|---|---|---|
| h1 | 1.00 | 0.45 | 0.44 | 0.48 | 0.42 | 0.36 | 0.40 | 0.37 | 0.38 | 0.37 | 0.34 | 0.35 | 0.37 | 0.36 | 0.36 | 0.33 | 0.38 |
| h2 | 0.45 | 1.00 | 0.38 | 0.43 | 0.40 | 0.36 | 0.38 | 0.33 | 0.35 | 0.30 | 0.29 | 0.30 | 0.32 | 0.32 | 0.31 | 0.30 | 0.35 |
| h3 | 0.46 | 0.41 | 1.00 | 0.50 | 0.42 | 0.36 | 0.38 | 0.34 | 0.36 | 0.37 | 0.36 | 0.38 | 0.39 | 0.38 | 0.37 | 0.35 | 0.39 |
| h4 | 0.47 | 0.42 | 0.48 | 1.00 | 0.46 | 0.40 | 0.41 | 0.40 | 0.40 | 0.40 | 0.35 | 0.34 | 0.38 | 0.42 | 0.44 | 0.35 | 0.41 |
| h5 | 0.43 | 0.41 | 0.42 | 0.47 | 1.00 | 0.38 | 0.42 | 0.38 | 0.39 | 0.35 | 0.34 | 0.34 | 0.34 | 0.37 | 0.40 | 0.32 | 0.38 |
| h6 | 0.36 | 0.35 | 0.34 | 0.40 | 0.36 | 1.00 | 0.42 | 0.42 | 0.39 | 0.30 | 0.32 | 0.30 | 0.30 | 0.35 | 0.34 | 0.34 | 0.35 |
| h7 | 0.41 | 0.39 | 0.37 | 0.42 | 0.42 | 0.43 | 1.00 | 0.42 | 0.43 | 0.31 | 0.33 | 0.32 | 0.36 | 0.37 | 0.38 | 0.38 | 0.38 |
| h8 | 0.38 | 0.34 | 0.33 | 0.40 | 0.37 | 0.43 | 0.43 | 1.00 | 0.44 | 0.37 | 0.38 | 0.35 | 0.36 | 0.41 | 0.38 | 0.40 | 0.38 |
| h9 | 0.37 | 0.34 | 0.35 | 0.40 | 0.38 | 0.39 | 0.42 | 0.43 | 1.00 | 0.34 | 0.34 | 0.34 | 0.35 | 0.41 | 0.39 | 0.40 | 0.38 |
| h10 | 0.38 | 0.31 | 0.37 | 0.41 | 0.34 | 0.31 | 0.32 | 0.38 | 0.35 | 1.00 | 0.33 | 0.37 | 0.38 | 0.40 | 0.39 | 0.37 | 0.36 |
| h11 | 0.36 | 0.31 | 0.34 | 0.36 | 0.32 | 0.34 | 0.34 | 0.39 | 0.36 | 0.32 | 1.00 | 0.36 | 0.38 | 0.38 | 0.32 | 0.37 | 0.35 |
| h12 | 0.34 | 0.29 | 0.35 | 0.35 | 0.30 | 0.30 | 0.32 | 0.34 | 0.33 | 0.33 | 0.34 | 1.00 | 0.37 | 0.39 | 0.32 | 0.37 | 0.33 |
| h13 | 0.36 | 0.31 | 0.37 | 0.37 | 0.31 | 0.29 | 0.34 | 0.34 | 0.33 | 0.34 | 0.35 | 0.36 | 1.00 | 0.44 | 0.38 | 0.40 | 0.35 |
| h14 | 0.37 | 0.31 | 0.37 | 0.42 | 0.36 | 0.36 | 0.37 | 0.41 | 0.40 | 0.39 | 0.38 | 0.40 | 0.46 | 1.00 | 0.41 | 0.44 | 0.39 |
| h15 | 0.37 | 0.31 | 0.35 | 0.44 | 0.38 | 0.34 | 0.37 | 0.38 | 0.39 | 0.37 | 0.31 | 0.32 | 0.39 | 0.40 | 1.00 | 0.37 | 0.37 |
| h16 | 0.32 | 0.23 | 0.32 | 0.35 | 0.29 | 0.34 | 0.38 | 0.39 | 0.39 | 0.34 | 0.35 | 0.36 | 0.40 | 0.43 | 0.37 | 1.00 | 0.35 |

(C)

| Name | H1 | H2 | H3 | H4 | H5 | H6 | H7 | H8 | H9 |
|---|---|---|---|---|---|---|---|---|---|
| L | 34,203 | 28,902 | 33,934 | 33,640 | 30,593 | 34,232 | 10,512 | 26,384 | 34,967 |
| R | 41,020 | 54,019 | 30,040 | 30,592 | 11,076 | 34,100 | 12,859 | 23,508 | 40,397 |
| Middle | 54,038 | 89,893 | 68,006 | 97,914 | 37,590 | 138,578 | 57,288 | 64,473 | 58,912 |

| Name | H10 | H11 | H12 | H13 | H14 | H15 | H16 | H17 |
|---|---|---|---|---|---|---|---|---|
| L | 30,185 | 33,926 | 31,869 | 37,101 | 24,660 | 43,258 | 22,135 | 21,410 |
| R | 21,138 | 40,700 | 38,094 | 42,690 | 22,275 | 48,002 | 25,054 | 26,905 |
| Middle | 32,393 | 59,089 | 62,829 | 74,275 | 83,046 | 68,806 | 90,890 | 87,837 |

(D)

| Name | h1 | h2 | h3 | h4 | h5 | h6 | h7 | h8 |
|---|---|---|---|---|---|---|---|---|
| L | 130,392 | 84,439 | 146,017 | 132,979 | 63,623 | 37,697 | 74,302 | 53,085 |
| R | 124,404 | 91,778 | 115,275 | 144,523 | 66,198 | 47,809 | 94,935 | 71,482 |
| Middle | 129,877 | 99,118 | 85,090 | 114,196 | 53,906 | 37,557 | 57,631 | 74,205 |

| Name | h9 | h10 | h11 | h12 | h13 | h14 | h15 | h16 |
|---|---|---|---|---|---|---|---|---|
| L | 77,095 | 26,003 | 19,672 | 61,211 | 78,778 | 65,605 | 65,965 | 62,709 |
| R | 84,097 | 24,971 | 13,844 | 58,375 | 86,673 | 72,452 | 77,093 | 73,329 |
| Middle | 71,141 | 26,204 | 13,228 | 57,135 | 115,407 | 76,677 | 59,769 | 74,315 |

**Supplemental Table 6. Computed correlation factors from reciprocal one-to-one registration foreach Z-stacks of *ngn1* plus *her6 samples*.** (A) Correlation factors (CFs) of GFP (pineal) alignmentsfrom*ngn1*-labeled samples (NG1 to NG21) and *her6*-labeled samples (H1 to H17)in wild-type (WT) groups. (B) CFs of GFP (pineal) alignments from *ngn1*-labeled samples (NG1 to NG21) in WT group and *her6*-labeled samples (h1 to h16) in *wls* group.

(A)

| Name | NG1 | NG2 | NG3 | NG4 | NG5 | NG6 | NG7 | NG8 | NG9 | NG10 | NG11 | NG12 | NG13 | NG14 | NG15 | NG16 | NG17 | NG18 | NG19 | NG20 | NG21 | H1 | H2 | H3 | H4 | H5 | H6 | H7 | H8 | H9 | H10 | H11 | H12 | H13 | H14 | H15 | H16 | H17 | Average |
|---|---|---|---|---|---|---|---|---|---|---|---|---|---|---|---|---|---|---|---|---|---|---|---|---|---|---|---|---|---|---|---|---|---|---|---|---|---|---|---|
| NG1 | 1.00 | 0.65 | 0.63 | 0.59 | 0.58 | 0.62 | 0.58 | 0.61 | 0.62 | 0.63 | 0.62 | 0.70 | 0.67 | 0.58 | 0.55 | 0.62 | 0.56 | 0.62 | 0.64 | 0.59 | 0.58 | 0.62 | 0.66 | 0.66 | 0.66 | 0.67 | 0.65 | 0.61 | 0.57 | 0.55 | 0.59 | 0.54 | 0.49 | 0.53 | 0.59 | 0.59 | 0.60 | 0.59 | 0.60 |
| NG2 | 0.65 | 1.00 | 0.62 | 0.59 | 0.59 | 0.65 | 0.58 | 0.62 | 0.60 | 0.60 | 0.59 | 0.68 | 0.69 | 0.58 | 0.51 | 0.62 | 0.51 | 0.63 | 0.59 | 0.58 | 0.55 | 0.65 | 0.66 | 0.64 | 0.62 | 0.64 | 0.65 | 0.61 | 0.57 | 0.61 | 0.53 | 0.57 | 0.52 | 0.55 | 0.61 | 0.61 | 0.64 | 0.55 | 0.60 |
| NG3 | 0.62 | 0.61 | 1.00 | 0.63 | 0.64 | 0.61 | 0.56 | 0.64 | 0.61 | 0.61 | 0.61 | 0.65 | 0.68 | 0.58 | 0.55 | 0.62 | 0.56 | 0.64 | 0.62 | 0.60 | 0.63 | 0.60 | 0.65 | 0.66 | 0.64 | 0.65 | 0.64 | 0.59 | 0.63 | 0.65 | 0.59 | 0.61 | 0.55 | 0.62 | 0.64 | 0.65 | 0.63 | 0.64 | 0.62 |
| NG4 | 0.58 | 0.59 | 0.63 | 1.00 | 0.61 | 0.62 | 0.56 | 0.67 | 0.55 | 0.63 | 0.57 | 0.67 | 0.66 | 0.58 | 0.57 | 0.58 | 0.53 | 0.67 | 0.60 | 0.65 | 0.62 | 0.60 | 0.63 | 0.65 | 0.60 | 0.59 | 0.64 | 0.59 | 0.62 | 0.65 | 0.64 | 0.59 | 0.61 | 0.62 | 0.62 | 0.64 | 0.66 | 0.67 | 0.62 |
| NG5 | 0.58 | 0.60 | 0.65 | 0.61 | 1.00 | 0.63 | 0.63 | 0.59 | 0.59 | 0.58 | 0.65 | 0.67 | 0.65 | 0.62 | 0.60 | 0.51 | 0.61 | 0.61 | 0.60 | 0.67 | 0.64 | 0.59 | 0.66 | 0.62 | 0.64 | 0.63 | 0.64 | 0.66 | 0.62 | 0.63 | 0.58 | 0.58 | 0.53 | 0.59 | 0.62 | 0.55 | 0.53 | 0.60 | 0.61 |
| NG6 | 0.61 | 0.65 | 0.62 | 0.63 | 0.63 | 1.00 | 0.59 | 0.61 | 0.63 | 0.63 | 0.66 | 0.71 | 0.68 | 0.61 | 0.57 | 0.59 | 0.56 | 0.65 | 0.63 | 0.66 | 0.63 | 0.61 | 0.67 | 0.66 | 0.65 | 0.65 | 0.70 | 0.64 | 0.64 | 0.66 | 0.62 | 0.60 | 0.53 | 0.53 | 0.63 | 0.61 | 0.65 | 0.62 | 0.63 |
| NG7 | 0.56 | 0.57 | 0.56 | 0.55 | 0.61 | 0.58 | 1.00 | 0.55 | 0.53 | 0.56 | 0.62 | 0.61 | 0.59 | 0.55 | 0.60 | 0.50 | 0.62 | 0.61 | 0.58 | 0.60 | 0.56 | 0.50 | 0.62 | 0.55 | 0.58 | 0.61 | 0.61 | 0.61 | 0.52 | 0.48 | 0.50 | 0.53 | 0.47 | 0.51 | 0.57 | 0.48 | 0.57 | 0.56 | 0.56 |
| NG8 | 0.61 | 0.62 | 0.64 | 0.67 | 0.59 | 0.61 | 0.55 | 1.00 | 0.60 | 0.61 | 0.59 | 0.70 | 0.67 | 0.58 | 0.53 | 0.58 | 0.54 | 0.65 | 0.57 | 0.64 | 0.63 | 0.63 | 0.62 | 0.65 | 0.63 | 0.60 | 0.65 | 0.59 | 0.61 | 0.64 | 0.60 | 0.56 | 0.60 | 0.63 | 0.64 | 0.63 | 0.64 | 0.66 | 0.61 |
| NG9 | 0.59 | 0.59 | 0.60 | 0.56 | 0.58 | 0.61 | 0.53 | 0.59 | 1.00 | 0.58 | 0.57 | 0.64 | 0.64 | 0.57 | 0.54 | 0.62 | 0.50 | 0.62 | 0.61 | 0.54 | 0.54 | 0.61 | 0.65 | 0.63 | 0.60 | 0.60 | 0.64 | 0.61 | 0.60 | 0.61 | 0.59 | 0.59 | 0.57 | 0.58 | 0.63 | 0.63 | 0.64 | 0.61 | 0.59 |
| NG10 | 0.61 | 0.59 | 0.61 | 0.62 | 0.56 | 0.62 | 0.56 | 0.60 | 0.59 | 1.00 | 0.59 | 0.65 | 0.64 | 0.48 | 0.60 | 0.60 | 0.58 | 0.59 | 0.66 | 0.57 | 0.58 | 0.60 | 0.61 | 0.64 | 0.64 | 0.58 | 0.65 | 0.62 | 0.57 | 0.59 | 0.60 | 0.52 | 0.41 | 0.55 | 0.60 | 0.56 | 0.61 | 0.64 | 0.59 |
| NG11 | 0.61 | 0.59 | 0.62 | 0.57 | 0.64 | 0.66 | 0.64 | 0.59 | 0.58 | 0.60 | 1.00 | 0.68 | 0.64 | 0.61 | 0.58 | 0.57 | 0.62 | 0.66 | 0.62 | 0.59 | 0.59 | 0.56 | 0.67 | 0.60 | 0.61 | 0.65 | 0.62 | 0.61 | 0.59 | 0.59 | 0.55 | 0.53 | 0.48 | 0.48 | 0.60 | 0.56 | 0.59 | 0.60 | 0.60 |
| NG12 | 0.71 | 0.69 | 0.67 | 0.68 | 0.68 | 0.72 | 0.64 | 0.71 | 0.67 | 0.67 | 0.70 | 1.00 | 0.74 | 0.67 | 0.65 | 0.67 | 0.63 | 0.73 | 0.69 | 0.71 | 0.70 | 0.68 | 0.73 | 0.74 | 0.70 | 0.74 | 0.71 | 0.72 | 0.67 | 0.70 | 0.67 | 0.65 | 0.59 | 0.65 | 0.71 | 0.68 | 0.70 | 0.71 | 0.69 |
| NG13 | 0.68 | 0.70 | 0.70 | 0.67 | 0.66 | 0.69 | 0.61 | 0.68 | 0.66 | 0.67 | 0.65 | 0.74 | 1.00 | 0.64 | 0.55 | 0.67 | 0.59 | 0.68 | 0.67 | 0.66 | 0.64 | 0.65 | 0.70 | 0.69 | 0.68 | 0.70 | 0.74 | 0.62 | 0.66 | 0.67 | 0.66 | 0.63 | 0.58 | 0.62 | 0.66 | 0.65 | 0.65 | 0.66 | 0.66 |
| NG14 | 0.60 | 0.60 | 0.61 | 0.62 | 0.64 | 0.63 | 0.59 | 0.61 | 0.60 | 0.52 | 0.64 | 0.68 | 0.67 | 1.00 | 0.57 | 0.61 | 0.57 | 0.67 | 0.56 | 0.67 | 0.66 | 0.58 | 0.66 | 0.63 | 0.59 | 0.67 | 0.63 | 0.59 | 0.60 | 0.65 | 0.53 | 0.55 | 0.51 | 0.53 | 0.62 | 0.61 | 0.66 | 0.61 | 0.61 |
| NG15 | 0.56 | 0.52 | 0.51 | 0.59 | 0.62 | 0.58 | 0.63 | 0.54 | 0.57 | 0.63 | 0.59 | 0.65 | 0.56 | 0.57 | 1.00 | 0.53 | 0.65 | 0.63 | 0.69 | 0.62 | 0.63 | 0.57 | 0.65 | 0.54 | 0.60 | 0.64 | 0.65 | 0.66 | 0.56 | 0.60 | 0.60 | 0.57 | 0.50 | 0.52 | 0.58 | 0.55 | 0.57 | 0.66 | 0.59 |
| NG16 | 0.63 | 0.64 | 0.65 | 0.60 | 0.52 | 0.61 | 0.53 | 0.60 | 0.66 | 0.63 | 0.58 | 0.68 | 0.68 | 0.61 | 0.53 | 1.00 | 0.54 | 0.68 | 0.66 | 0.57 | 0.59 | 0.61 | 0.64 | 0.66 | 0.63 | 0.61 | 0.67 | 0.61 | 0.59 | 0.63 | 0.61 | 0.64 | 0.60 | 0.59 | 0.66 | 0.64 | 0.69 | 0.66 | 0.62 |
| NG17 | 0.57 | 0.54 | 0.58 | 0.54 | 0.62 | 0.57 | 0.65 | 0.56 | 0.52 | 0.61 | 0.63 | 0.63 | 0.59 | 0.56 | 0.64 | 0.53 | 1.00 | 0.52 | 0.63 | 0.57 | 0.56 | 0.52 | 0.66 | 0.56 | 0.63 | 0.64 | 0.62 | 0.62 | 0.53 | 0.55 | 0.54 | 0.51 | 0.43 | 0.49 | 0.60 | 0.52 | 0.56 | 0.56 | 0.57 |
| NG18 | 0.64 | 0.65 | 0.67 | 0.69 | 0.64 | 0.67 | 0.65 | 0.68 | 0.66 | 0.66 | 0.68 | 0.74 | 0.70 | 0.67 | 0.62 | 0.68 | 0.62 | 1.00 | 0.67 | 0.64 | 0.67 | 0.64 | 0.72 | 0.69 | 0.68 | 0.69 | 0.70 | 0.67 | 0.65 | 0.73 | 0.65 | 0.64 | 0.60 | 0.63 | 0.73 | 0.68 | 0.72 | 0.72 | 0.67 |
| NG19 | 0.65 | 0.61 | 0.65 | 0.62 | 0.62 | 0.65 | 0.61 | 0.59 | 0.64 | 0.69 | 0.64 | 0.70 | 0.68 | 0.55 | 0.69 | 0.66 | 0.63 | 0.67 | 1.00 | 0.62 | 0.61 | 0.66 | 0.68 | 0.63 | 0.68 | 0.67 | 0.73 | 0.67 | 0.62 | 0.62 | 0.62 | 0.56 | 0.51 | 0.60 | 0.65 | 0.62 | 0.66 | 0.67 | 0.64 |
| NG20 | 0.60 | 0.59 | 0.62 | 0.67 | 0.69 | 0.68 | 0.63 | 0.66 | 0.57 | 0.59 | 0.61 | 0.72 | 0.67 | 0.67 | 0.61 | 0.58 | 0.57 | 0.65 | 0.61 | 1.00 | 0.74 | 0.63 | 0.68 | 0.65 | 0.61 | 0.64 | 0.66 | 0.69 | 0.68 | 0.71 | 0.67 | 0.61 | 0.59 | 0.63 | 0.65 | 0.64 | 0.63 | 0.65 | 0.64 |
| NG21 | 0.59 | 0.57 | 0.66 | 0.64 | 0.65 | 0.64 | 0.59 | 0.65 | 0.61 | 0.61 | 0.61 | 0.71 | 0.65 | 0.66 | 0.63 | 0.59 | 0.58 | 0.67 | 0.61 | 0.74 | 1.00 | 0.61 | 0.62 | 0.68 | 0.62 | 0.65 | 0.64 | 0.68 | 0.68 | 0.71 | 0.68 | 0.62 | 0.58 | 0.64 | 0.67 | 0.65 | 0.66 | 0.72 | 0.64 |
| H1 | 0.62 | 0.64 | 0.60 | 0.61 | 0.58 | 0.61 | 0.51 | 0.63 | 0.62 | 0.61 | 0.56 | 0.67 | 0.64 | 0.56 | 0.55 | 0.59 | 0.51 | 0.62 | 0.64 | 0.62 | 0.59 | 1.00 | 0.64 | 0.66 | 0.62 | 0.63 | 0.68 | 0.61 | 0.61 | 0.63 | 0.60 | 0.57 | 0.55 | 0.61 | 0.62 | 0.60 | 0.66 | 0.64 | 0.61 |
| H2 | 0.67 | 0.66 | 0.66 | 0.61 | 0.67 | 0.68 | 0.63 | 0.62 | 0.67 | 0.62 | 0.67 | 0.73 | 0.70 | 0.64 | 0.63 | 0.62 | 0.65 | 0.70 | 0.66 | 0.67 | 0.60 | 0.65 | 1.00 | 0.70 | 0.70 | 0.75 | 0.70 | 0.68 | 0.64 | 0.65 | 0.64 | 0.58 | 0.51 | 0.58 | 0.65 | 0.64 | 0.67 | 0.65 | 0.65 |
| H3 | 0.66 | 0.64 | 0.68 | 0.66 | 0.62 | 0.66 | 0.57 | 0.66 | 0.65 | 0.66 | 0.60 | 0.74 | 0.69 | 0.62 | 0.54 | 0.64 | 0.55 | 0.68 | 0.62 | 0.64 | 0.65 | 0.66 | 0.70 | 1.00 | 0.68 | 0.66 | 0.68 | 0.65 | 0.68 | 0.69 | 0.61 | 0.59 | 0.62 | 0.64 | 0.68 | 0.64 | 0.68 | 0.68 | 0.65 |
| H4 | 0.66 | 0.62 | 0.65 | 0.60 | 0.64 | 0.66 | 0.59 | 0.63 | 0.63 | 0.65 | 0.62 | 0.69 | 0.67 | 0.57 | 0.59 | 0.62 | 0.62 | 0.66 | 0.67 | 0.61 | 0.61 | 0.63 | 0.70 | 0.68 | 1.00 | 0.68 | 0.70 | 0.67 | 0.58 | 0.61 | 0.62 | 0.60 | 0.53 | 0.59 | 0.64 | 0.64 | 0.63 | 0.66 | 0.63 |
| H5 | 0.66 | 0.63 | 0.66 | 0.59 | 0.62 | 0.65 | 0.61 | 0.60 | 0.61 | 0.59 | 0.65 | 0.73 | 0.68 | 0.64 | 0.62 | 0.58 | 0.62 | 0.67 | 0.65 | 0.62 | 0.63 | 0.63 | 0.75 | 0.65 | 0.67 | 1.00 | 0.68 | 0.67 | 0.59 | 0.62 | 0.60 | 0.53 | 0.55 | 0.57 | 0.62 | 0.63 | 0.63 | 0.62 | 0.63 |
| H6 | 0.65 | 0.65 | 0.65 | 0.65 | 0.64 | 0.71 | 0.61 | 0.65 | 0.66 | 0.66 | 0.62 | 0.70 | 0.73 | 0.61 | 0.64 | 0.65 | 0.61 | 0.68 | 0.72 | 0.65 | 0.63 | 0.68 | 0.70 | 0.68 | 0.70 | 0.68 | 1.00 | 0.67 | 0.64 | 0.66 | 0.65 | 0.63 | 0.59 | 0.63 | 0.68 | 0.67 | 0.69 | 0.68 | 0.66 |
| H7 | 0.60 | 0.60 | 0.60 | 0.59 | 0.65 | 0.64 | 0.62 | 0.59 | 0.63 | 0.63 | 0.61 | 0.71 | 0.62 | 0.57 | 0.64 | 0.59 | 0.61 | 0.64 | 0.65 | 0.67 | 0.66 | 0.61 | 0.68 | 0.64 | 0.67 | 0.67 | 0.67 | 1.00 | 0.62 | 0.64 | 0.62 | 0.58 | 0.55 | 0.56 | 0.64 | 0.62 | 0.60 | 0.67 | 0.63 |
| H8 | 0.56 | 0.56 | 0.63 | 0.61 | 0.60 | 0.62 | 0.51 | 0.60 | 0.60 | 0.57 | 0.57 | 0.65 | 0.64 | 0.55 | 0.53 | 0.56 | 0.50 | 0.61 | 0.59 | 0.64 | 0.64 | 0.61 | 0.63 | 0.65 | 0.57 | 0.59 | 0.63 | 0.61 | 1.00 | 0.64 | 0.63 | 0.61 | 0.58 | 0.63 | 0.60 | 0.59 | 0.60 | 0.62 | 0.60 |
| H9 | 0.55 | 0.61 | 0.67 | 0.67 | 0.63 | 0.68 | 0.53 | 0.65 | 0.63 | 0.60 | 0.60 | 0.70 | 0.67 | 0.64 | 0.59 | 0.63 | 0.54 | 0.72 | 0.62 | 0.70 | 0.70 | 0.65 | 0.66 | 0.69 | 0.62 | 0.64 | 0.67 | 0.65 | 0.66 | 1.00 | 0.72 | 0.67 | 0.70 | 0.70 | 0.67 | 0.69 | 0.68 | 0.68 | 0.65 |
| H10 | 0.60 | 0.54 | 0.60 | 0.65 | 0.59 | 0.62 | 0.52 | 0.61 | 0.61 | 0.61 | 0.56 | 0.66 | 0.65 | 0.52 | 0.54 | 0.60 | 0.53 | 0.64 | 0.62 | 0.65 | 0.66 | 0.62 | 0.65 | 0.61 | 0.63 | 0.62 | 0.67 | 0.63 | 0.65 | 0.72 | 1.00 | 0.67 | 0.66 | 0.70 | 0.65 | 0.63 | 0.63 | 0.66 | 0.62 |
| H11 | 0.55 | 0.59 | 0.63 | 0.61 | 0.59 | 0.61 | 0.56 | 0.58 | 0.62 | 0.55 | 0.55 | 0.65 | 0.64 | 0.54 | 0.57 | 0.64 | 0.51 | 0.64 | 0.56 | 0.62 | 0.63 | 0.58 | 0.59 | 0.64 | 0.61 | 0.58 | 0.65 | 0.60 | 0.64 | 0.68 | 0.68 | 1.00 | 0.65 | 0.69 | 0.62 | 0.65 | 0.64 | 0.65 | 0.61 |
| H12 | 0.51 | 0.53 | 0.61 | 0.63 | 0.54 | 0.56 | 0.49 | 0.62 | 0.60 | 0.54 | 0.49 | 0.59 | 0.58 | 0.51 | 0.50 | 0.60 | 0.43 | 0.60 | 0.52 | 0.59 | 0.58 | 0.61 | 0.52 | 0.63 | 0.54 | 0.57 | 0.61 | 0.56 | 0.60 | 0.70 | 0.67 | 0.65 | 1.00 | 0.74 | 0.62 | 0.66 | 0.66 | 0.64 | 0.58 |
| H13 | 0.54 | 0.56 | 0.64 | 0.64 | 0.60 | 0.54 | 0.53 | 0.65 | 0.63 | 0.58 | 0.49 | 0.65 | 0.63 | 0.52 | 0.52 | 0.58 | 0.49 | 0.67 | 0.61 | 0.62 | 0.64 | 0.63 | 0.59 | 0.65 | 0.60 | 0.59 | 0.65 | 0.58 | 0.65 | 0.70 | 0.71 | 0.66 | 0.74 | 1.00 | 0.65 | 0.67 | 0.67 | 0.70 | 0.61 |
| H14 | 0.60 | 0.62 | 0.66 | 0.61 | 0.63 | 0.64 | 0.59 | 0.66 | 0.66 | 0.62 | 0.61 | 0.71 | 0.67 | 0.62 | 0.57 | 0.66 | 0.60 | 0.73 | 0.65 | 0.64 | 0.66 | 0.64 | 0.67 | 0.69 | 0.65 | 0.64 | 0.69 | 0.65 | 0.62 | 0.67 | 0.66 | 0.62 | 0.61 | 0.64 | 1.00 | 0.67 | 0.73 | 0.72 | 0.65 |
| H15 | 0.60 | 0.62 | 0.67 | 0.66 | 0.56 | 0.62 | 0.50 | 0.65 | 0.65 | 0.58 | 0.57 | 0.69 | 0.65 | 0.61 | 0.54 | 0.64 | 0.53 | 0.67 | 0.61 | 0.63 | 0.64 | 0.62 | 0.65 | 0.65 | 0.63 | 0.64 | 0.68 | 0.64 | 0.64 | 0.69 | 0.63 | 0.65 | 0.66 | 0.66 | 0.67 | 1.00 | 0.70 | 0.67 | 0.63 |
| H16 | 0.61 | 0.66 | 0.65 | 0.67 | 0.54 | 0.67 | 0.59 | 0.66 | 0.67 | 0.64 | 0.60 | 0.70 | 0.66 | 0.66 | 0.57 | 0.69 | 0.56 | 0.72 | 0.67 | 0.63 | 0.65 | 0.68 | 0.68 | 0.68 | 0.64 | 0.65 | 0.70 | 0.61 | 0.62 | 0.68 | 0.63 | 0.64 | 0.65 | 0.66 | 0.73 | 0.70 | 1.00 | 0.67 | 0.65 |
| H17 | 0.60 | 0.57 | 0.66 | 0.69 | 0.62 | 0.64 | 0.58 | 0.68 | 0.65 | 0.66 | 0.62 | 0.72 | 0.66 | 0.59 | 0.66 | 0.66 | 0.57 | 0.72 | 0.68 | 0.64 | 0.71 | 0.66 | 0.66 | 0.70 | 0.67 | 0.64 | 0.70 | 0.69 | 0.64 | 0.69 | 0.67 | 0.65 | 0.64 | 0.70 | 0.72 | 0.67 | 0.67 | 1.00 | 0.66 |

(B)

| Name | NG1 | NG2 | NG3 | NG4 | NG5 | NG6 | NG7 | NG8 | NG9 | NG10 | NG11 | NG12 | NG13 | NG14 | NG15 | NG16 | NG17 | NG18 | NG19 | NG20 | NG21 | h1 | h2 | h3 | h4 | h5 | h6 | h7 | h8 | h9 | h10 | h11 | h12 | h13 | h14 | h15 | h16 | Average |
|---|---|---|---|---|---|---|---|---|---|---|---|---|---|---|---|---|---|---|---|---|---|---|---|---|---|---|---|---|---|---|---|---|---|---|---|---|---|---|
| NG1 | 1.00 | 0.52 | 0.63 | 0.57 | 0.56 | 0.57 | 0.62 | 0.53 | 0.54 | 0.59 | 0.62 | 0.60 | 0.56 | 0.55 | 0.47 | 0.56 | 0.54 | 0.50 | 0.53 | 0.49 | 0.53 | 0.57 | 0.54 | 0.58 | 0.54 | 0.62 | 0.57 | 0.65 | 0.56 | 0.52 | 0.61 | 0.63 | 0.63 | 0.59 | 0.56 | 0.64 | 0.53 | 0.57 |
| NG2 | 0.62 | 1.00 | 0.67 | 0.64 | 0.67 | 0.62 | 0.63 | 0.63 | 0.64 | 0.64 | 0.63 | 0.64 | 0.67 | 0.64 | 0.63 | 0.61 | 0.62 | 0.65 | 0.61 | 0.66 | 0.66 | 0.64 | 0.65 | 0.67 | 0.69 | 0.64 | 0.67 | 0.65 | 0.68 | 0.66 | 0.66 | 0.68 | 0.74 | 0.70 | 0.67 | 0.62 | 0.68 | 0.65 |
| NG3 | 0.63 | 0.67 | 1.00 | 0.62 | 0.61 | 0.60 | 0.65 | 0.56 | 0.60 | 0.64 | 0.63 | 0.62 | 0.65 | 0.66 | 0.56 | 0.62 | 0.57 | 0.55 | 0.57 | 0.59 | 0.61 | 0.65 | 0.61 | 0.65 | 0.62 | 0.62 | 0.65 | 0.61 | 0.59 | 0.64 | 0.69 | 0.64 | 0.70 | 0.68 | 0.55 | 0.69 | 0.66 | 0.62 |
| NG4 | 0.57 | 0.65 | 0.61 | 1.00 | 0.74 | 0.62 | 0.57 | 0.64 | 0.62 | 0.63 | 0.63 | 0.67 | 0.64 | 0.62 | 0.62 | 0.68 | 0.70 | 0.66 | 0.61 | 0.63 | 0.63 | 0.60 | 0.59 | 0.62 | 0.67 | 0.69 | 0.68 | 0.63 | 0.66 | 0.57 | 0.59 | 0.61 | 0.72 | 0.67 | 0.67 | 0.61 | 0.58 | 0.64 |
| NG5 | 0.58 | 0.67 | 0.61 | 0.74 | 1.00 | 0.55 | 0.56 | 0.64 | 0.64 | 0.64 | 0.65 | 0.67 | 0.64 | 0.63 | 0.62 | 0.68 | 0.71 | 0.65 | 0.64 | 0.63 | 0.63 | 0.59 | 0.57 | 0.66 | 0.64 | 0.65 | 0.64 | 0.59 | 0.65 | 0.61 | 0.61 | 0.61 | 0.71 | 0.65 | 0.66 | 0.63 | 0.59 | 0.63 |
| NG6 | 0.54 | 0.59 | 0.57 | 0.59 | 0.55 | 1.00 | 0.62 | 0.58 | 0.59 | 0.56 | 0.55 | 0.57 | 0.55 | 0.54 | 0.52 | 0.52 | 0.57 | 0.54 | 0.53 | 0.55 | 0.54 | 0.62 | 0.62 | 0.59 | 0.58 | 0.57 | 0.61 | 0.54 | 0.55 | 0.58 | 0.56 | 0.60 | 0.66 | 0.64 | 0.56 | 0.50 | 0.56 | 0.57 |
| NG7 | 0.61 | 0.60 | 0.63 | 0.56 | 0.54 | 0.63 | 1.00 | 0.57 | 0.53 | 0.61 | 0.63 | 0.63 | 0.62 | 0.55 | 0.50 | 0.55 | 0.55 | 0.48 | 0.55 | 0.54 | 0.54 | 0.65 | 0.62 | 0.61 | 0.59 | 0.58 | 0.62 | 0.61 | 0.59 | 0.59 | 0.58 | 0.64 | 0.65 | 0.66 | 0.59 | 0.60 | 0.56 | 0.59 |
| NG8 | 0.51 | 0.59 | 0.54 | 0.63 | 0.62 | 0.59 | 0.57 | 1.00 | 0.62 | 0.61 | 0.61 | 0.60 | 0.57 | 0.53 | 0.55 | 0.59 | 0.62 | 0.57 | 0.56 | 0.57 | 0.58 | 0.61 | 0.59 | 0.59 | 0.58 | 0.58 | 0.61 | 0.54 | 0.59 | 0.57 | 0.52 | 0.56 | 0.66 | 0.62 | 0.57 | 0.53 | 0.51 | 0.58 |
| NG9 | 0.52 | 0.61 | 0.57 | 0.61 | 0.63 | 0.60 | 0.53 | 0.62 | 1.00 | 0.58 | 0.60 | 0.56 | 0.60 | 0.51 | 0.55 | 0.60 | 0.65 | 0.57 | 0.52 | 0.58 | 0.54 | 0.61 | 0.56 | 0.61 | 0.59 | 0.59 | 0.65 | 0.52 | 0.64 | 0.59 | 0.59 | 0.55 | 0.65 | 0.63 | 0.60 | 0.51 | 0.58 | 0.58 |
| NG10 | 0.58 | 0.62 | 0.62 | 0.61 | 0.63 | 0.57 | 0.61 | 0.61 | 0.58 | 1.00 | 0.67 | 0.61 | 0.61 | 0.58 | 0.58 | 0.61 | 0.64 | 0.56 | 0.59 | 0.55 | 0.59 | 0.57 | 0.59 | 0.56 | 0.60 | 0.59 | 0.62 | 0.56 | 0.56 | 0.54 | 0.59 | 0.56 | 0.66 | 0.61 | 0.55 | 0.61 | 0.54 | 0.59 |
| NG11 | 0.61 | 0.61 | 0.62 | 0.61 | 0.63 | 0.57 | 0.63 | 0.62 | 0.61 | 0.67 | 1.00 | 0.67 | 0.62 | 0.59 | 0.60 | 0.65 | 0.65 | 0.56 | 0.59 | 0.57 | 0.61 | 0.60 | 0.59 | 0.61 | 0.61 | 0.63 | 0.62 | 0.59 | 0.61 | 0.59 | 0.59 | 0.62 | 0.65 | 0.62 | 0.58 | 0.62 | 0.52 | 0.61 |
| NG12 | 0.57 | 0.61 | 0.59 | 0.64 | 0.64 | 0.57 | 0.62 | 0.60 | 0.56 | 0.61 | 0.66 | 1.00 | 0.61 | 0.57 | 0.57 | 0.60 | 0.64 | 0.57 | 0.54 | 0.55 | 0.59 | 0.59 | 0.60 | 0.62 | 0.61 | 0.62 | 0.62 | 0.61 | 0.59 | 0.62 | 0.57 | 0.60 | 0.66 | 0.63 | 0.59 | 0.60 | 0.55 | 0.60 |
| NG13 | 0.56 | 0.67 | 0.65 | 0.64 | 0.64 | 0.59 | 0.63 | 0.59 | 0.62 | 0.62 | 0.64 | 0.63 | 1.00 | 0.70 | 0.71 | 0.70 | 0.69 | 0.66 | 0.69 | 0.74 | 0.66 | 0.59 | 0.62 | 0.68 | 0.66 | 0.63 | 0.63 | 0.55 | 0.64 | 0.65 | 0.58 | 0.59 | 0.69 | 0.67 | 0.62 | 0.61 | 0.62 | 0.64 |
| NG14 | 0.55 | 0.64 | 0.66 | 0.62 | 0.63 | 0.56 | 0.56 | 0.55 | 0.53 | 0.59 | 0.61 | 0.60 | 0.70 | 1.00 | 0.69 | 0.71 | 0.63 | 0.65 | 0.63 | 0.67 | 0.72 | 0.59 | 0.59 | 0.61 | 0.62 | 0.58 | 0.60 | 0.57 | 0.61 | 0.62 | 0.61 | 0.60 | 0.65 | 0.63 | 0.48 | 0.62 | 0.61 | 0.61 |
| NG15 | 0.46 | 0.62 | 0.55 | 0.62 | 0.61 | 0.55 | 0.51 | 0.56 | 0.56 | 0.59 | 0.61 | 0.59 | 0.71 | 0.68 | 1.00 | 0.71 | 0.71 | 0.68 | 0.67 | 0.74 | 0.69 | 0.54 | 0.57 | 0.64 | 0.64 | 0.58 | 0.57 | 0.52 | 0.65 | 0.60 | 0.59 | 0.53 | 0.63 | 0.63 | 0.54 | 0.54 | 0.55 | 0.60 |
| NG16 | 0.57 | 0.61 | 0.62 | 0.68 | 0.68 | 0.55 | 0.57 | 0.62 | 0.62 | 0.63 | 0.67 | 0.64 | 0.71 | 0.71 | 0.72 | 1.00 | 0.77 | 0.71 | 0.70 | 0.69 | 0.74 | 0.57 | 0.60 | 0.63 | 0.65 | 0.64 | 0.62 | 0.58 | 0.65 | 0.60 | 0.65 | 0.53 | 0.65 | 0.64 | 0.58 | 0.64 | 0.56 | 0.64 |
| NG17 | 0.54 | 0.65 | 0.58 | 0.70 | 0.71 | 0.60 | 0.56 | 0.64 | 0.68 | 0.65 | 0.66 | 0.67 | 0.69 | 0.63 | 0.72 | 0.77 | 1.00 | 0.76 | 0.72 | 0.68 | 0.73 | 0.60 | 0.62 | 0.62 | 0.68 | 0.61 | 0.64 | 0.54 | 0.65 | 0.66 | 0.62 | 0.59 | 0.69 | 0.68 | 0.59 | 0.62 | 0.62 | 0.65 |
| NG18 | 0.50 | 0.65 | 0.54 | 0.66 | 0.65 | 0.56 | 0.50 | 0.58 | 0.60 | 0.58 | 0.57 | 0.59 | 0.66 | 0.65 | 0.68 | 0.70 | 0.75 | 1.00 | 0.67 | 0.69 | 0.69 | 0.55 | 0.54 | 0.61 | 0.64 | 0.56 | 0.62 | 0.51 | 0.61 | 0.63 | 0.61 | 0.55 | 0.63 | 0.62 | 0.53 | 0.54 | 0.62 | 0.61 |
| NG19 | 0.53 | 0.61 | 0.58 | 0.60 | 0.64 | 0.56 | 0.57 | 0.58 | 0.53 | 0.61 | 0.60 | 0.57 | 0.69 | 0.63 | 0.68 | 0.69 | 0.72 | 0.67 | 1.00 | 0.68 | 0.72 | 0.56 | 0.53 | 0.60 | 0.63 | 0.59 | 0.57 | 0.52 | 0.61 | 0.60 | 0.57 | 0.48 | 0.61 | 0.61 | 0.55 | 0.55 | 0.58 | 0.60 |
| NG20 | 0.49 | 0.67 | 0.59 | 0.64 | 0.63 | 0.58 | 0.55 | 0.60 | 0.59 | 0.56 | 0.59 | 0.57 | 0.74 | 0.67 | 0.75 | 0.69 | 0.69 | 0.69 | 0.68 | 1.00 | 0.68 | 0.54 | 0.59 | 0.63 | 0.66 | 0.60 | 0.59 | 0.56 | 0.65 | 0.62 | 0.61 | 0.54 | 0.64 | 0.60 | 0.60 | 0.53 | 0.62 | 0.62 |
| NG21 | 0.53 | 0.66 | 0.61 | 0.63 | 0.63 | 0.56 | 0.56 | 0.58 | 0.56 | 0.60 | 0.62 | 0.62 | 0.66 | 0.72 | 0.69 | 0.73 | 0.72 | 0.70 | 0.72 | 0.68 | 1.00 | 0.56 | 0.57 | 0.57 | 0.63 | 0.58 | 0.58 | 0.55 | 0.61 | 0.63 | 0.61 | 0.54 | 0.67 | 0.63 | 0.58 | 0.63 | 0.63 | 0.62 |
| h1 | 0.56 | 0.62 | 0.64 | 0.59 | 0.58 | 0.64 | 0.66 | 0.62 | 0.61 | 0.57 | 0.60 | 0.62 | 0.58 | 0.57 | 0.53 | 0.55 | 0.59 | 0.54 | 0.55 | 0.53 | 0.55 | 1.00 | 0.65 | 0.63 | 0.59 | 0.58 | 0.62 | 0.58 | 0.61 | 0.62 | 0.63 | 0.62 | 0.70 | 0.67 | 0.58 | 0.55 | 0.62 | 0.60 |
| h2 | 0.51 | 0.63 | 0.59 | 0.58 | 0.55 | 0.63 | 0.62 | 0.59 | 0.57 | 0.59 | 0.59 | 0.61 | 0.61 | 0.58 | 0.55 | 0.58 | 0.61 | 0.56 | 0.52 | 0.58 | 0.56 | 0.65 | 1.00 | 0.62 | 0.59 | 0.59 | 0.65 | 0.58 | 0.62 | 0.60 | 0.60 | 0.59 | 0.68 | 0.69 | 0.58 | 0.51 | 0.62 | 0.59 |
| h3 | 0.56 | 0.64 | 0.62 | 0.60 | 0.63 | 0.59 | 0.60 | 0.59 | 0.60 | 0.55 | 0.60 | 0.62 | 0.66 | 0.59 | 0.62 | 0.60 | 0.62 | 0.59 | 0.57 | 0.60 | 0.55 | 0.62 | 0.61 | 1.00 | 0.63 | 0.64 | 0.61 | 0.56 | 0.64 | 0.61 | 0.61 | 0.61 | 0.65 | 0.68 | 0.58 | 0.55 | 0.62 | 0.61 |
| h4 | 0.53 | 0.67 | 0.60 | 0.65 | 0.62 | 0.59 | 0.58 | 0.58 | 0.59 | 0.59 | 0.61 | 0.62 | 0.62 | 0.60 | 0.63 | 0.62 | 0.66 | 0.61 | 0.60 | 0.64 | 0.62 | 0.58 | 0.59 | 0.63 | 1.00 | 0.61 | 0.62 | 0.56 | 0.67 | 0.55 | 0.63 | 0.57 | 0.67 | 0.66 | 0.58 | 0.57 | 0.58 | 0.61 |
| h5 | 0.61 | 0.61 | 0.60 | 0.67 | 0.64 | 0.57 | 0.58 | 0.59 | 0.60 | 0.59 | 0.63 | 0.64 | 0.62 | 0.57 | 0.58 | 0.62 | 0.59 | 0.55 | 0.57 | 0.58 | 0.56 | 0.58 | 0.60 | 0.65 | 0.61 | 1.00 | 0.63 | 0.63 | 0.59 | 0.59 | 0.58 | 0.65 | 0.67 | 0.65 | 0.62 | 0.60 | 0.51 | 0.60 |
| h6 | 0.56 | 0.65 | 0.63 | 0.66 | 0.63 | 0.61 | 0.62 | 0.62 | 0.65 | 0.62 | 0.62 | 0.63 | 0.62 | 0.58 | 0.56 | 0.60 | 0.63 | 0.61 | 0.55 | 0.58 | 0.56 | 0.61 | 0.65 | 0.62 | 0.63 | 0.63 | 1.00 | 0.59 | 0.61 | 0.63 | 0.63 | 0.66 | 0.71 | 0.68 | 0.61 | 0.57 | 0.59 | 0.62 |
| h7 | 0.62 | 0.61 | 0.58 | 0.60 | 0.56 | 0.54 | 0.60 | 0.54 | 0.51 | 0.55 | 0.58 | 0.61 | 0.52 | 0.55 | 0.50 | 0.50 | 0.51 | 0.50 | 0.49 | 0.54 | 0.53 | 0.56 | 0.57 | 0.56 | 0.55 | 0.61 | 0.58 | 1.00 | 0.55 | 0.53 | 0.56 | 0.62 | 0.61 | 0.59 | 0.55 | 0.60 | 0.50 | 0.56 |
| h8 | 0.54 | 0.65 | 0.57 | 0.64 | 0.63 | 0.57 | 0.59 | 0.59 | 0.64 | 0.56 | 0.60 | 0.59 | 0.62 | 0.59 | 0.63 | 0.62 | 0.63 | 0.59 | 0.59 | 0.63 | 0.59 | 0.61 | 0.62 | 0.64 | 0.67 | 0.59 | 0.61 | 0.55 | 1.00 | 0.60 | 0.61 | 0.59 | 0.70 | 0.67 | 0.58 | 0.53 | 0.58 | 0.60 |
| h9 | 0.50 | 0.62 | 0.61 | 0.54 | 0.54 | 0.57 | 0.57 | 0.56 | 0.58 | 0.52 | 0.57 | 0.62 | 0.61 | 0.58 | 0.58 | 0.56 | 0.62 | 0.59 | 0.57 | 0.59 | 0.60 | 0.59 | 0.59 | 0.60 | 0.56 | 0.58 | 0.61 | 0.53 | 0.59 | 1.00 | 0.58 | 0.57 | 0.64 | 0.64 | 0.57 | 0.54 | 0.62 | 0.58 |
| h10 | 0.58 | 0.59 | 0.66 | 0.57 | 0.58 | 0.57 | 0.57 | 0.52 | 0.59 | 0.58 | 0.59 | 0.56 | 0.55 | 0.58 | 0.56 | 0.61 | 0.59 | 0.58 | 0.54 | 0.58 | 0.58 | 0.61 | 0.59 | 0.61 | 0.62 | 0.56 | 0.62 | 0.56 | 0.60 | 0.59 | 1.00 | 0.59 | 0.65 | 0.64 | 0.48 | 0.60 | 0.60 | 0.58 |
| h11 | 0.62 | 0.66 | 0.62 | 0.59 | 0.59 | 0.60 | 0.64 | 0.57 | 0.55 | 0.58 | 0.60 | 0.62 | 0.58 | 0.58 | 0.45 | 0.51 | 0.57 | 0.49 | 0.46 | 0.51 | 0.53 | 0.61 | 0.59 | 0.62 | 0.57 | 0.64 | 0.66 | 0.64 | 0.59 | 0.58 | 0.60 | 1.00 | 0.68 | 0.64 | 0.61 | 0.58 | 0.57 | 0.59 |
| h12 | 0.63 | 0.73 | 0.69 | 0.71 | 0.70 | 0.69 | 0.66 | 0.67 | 0.67 | 0.67 | 0.66 | 0.69 | 0.68 | 0.65 | 0.63 | 0.64 | 0.69 | 0.63 | 0.61 | 0.64 | 0.66 | 0.71 | 0.69 | 0.67 | 0.68 | 0.68 | 0.72 | 0.64 | 0.71 | 0.67 | 0.67 | 0.70 | 1.00 | 0.74 | 0.67 | 0.65 | 0.67 | 0.67 |
| h13 | 0.59 | 0.68 | 0.67 | 0.66 | 0.64 | 0.67 | 0.67 | 0.63 | 0.64 | 0.62 | 0.63 | 0.66 | 0.66 | 0.62 | 0.62 | 0.63 | 0.67 | 0.62 | 0.60 | 0.60 | 0.62 | 0.68 | 0.70 | 0.70 | 0.67 | 0.66 | 0.69 | 0.61 | 0.68 | 0.66 | 0.67 | 0.65 | 0.74 | 1.00 | 0.64 | 0.55 | 0.67 | 0.65 |
| h14 | 0.57 | 0.67 | 0.56 | 0.67 | 0.66 | 0.60 | 0.61 | 0.60 | 0.62 | 0.58 | 0.61 | 0.63 | 0.62 | 0.49 | 0.54 | 0.58 | 0.59 | 0.58 | 0.55 | 0.60 | 0.54 | 0.60 | 0.60 | 0.61 | 0.62 | 0.64 | 0.63 | 0.59 | 0.61 | 0.60 | 0.52 | 0.64 | 0.68 | 0.67 | 1.00 | 0.57 | 0.61 | 0.60 |
| h15 | 0.65 | 0.63 | 0.69 | 0.62 | 0.63 | 0.53 | 0.62 | 0.55 | 0.53 | 0.62 | 0.63 | 0.63 | 0.61 | 0.62 | 0.55 | 0.64 | 0.62 | 0.54 | 0.55 | 0.53 | 0.63 | 0.56 | 0.52 | 0.51 | 0.59 | 0.62 | 0.58 | 0.63 | 0.54 | 0.57 | 0.63 | 0.59 | 0.65 | 0.56 | 0.57 | 1.00 | 0.53 | 0.59 |
| h16 | 0.54 | 0.68 | 0.66 | 0.57 | 0.59 | 0.59 | 0.58 | 0.56 | 0.61 | 0.55 | 0.54 | 0.58 | 0.62 | 0.61 | 0.56 | 0.57 | 0.62 | 0.61 | 0.57 | 0.62 | 0.63 | 0.63 | 0.64 | 0.65 | 0.60 | 0.52 | 0.61 | 0.53 | 0.60 | 0.66 | 0.63 | 0.58 | 0.68 | 0.68 | 0.61 | 0.53 | 1.00 | 0.60 |

**Supplemental Table 7. Computed correlation factors from reciprocal one-to-one registration foreach Z-stacks of *foxD3:*GFP signals**. (A) Correlation factors (CFs) fromimage registrationresults. NC1 to NC11 or NJ1 to NJ10 are names for samples from the injection controlor the *ngn1*:*her6* plasmid-injected groups, respectively. (B-C) Volumes (Vol.) of*ngn1* expressing domain (μm$^{3}$) from samples of injectioncontrol (B)or *ngn1*:*her6* plasmid-injected (C) groups. Intensity threshold (I) = 18 for vol. calculation.

(A)

| Name | NC1 | NC2 | NC3 | NC4 | NC5 | NC6 | NC7 | NC8 | NC9 | NC10 | NC11 | NJ1 | NJ2 | NJ3 | NJ4 | NJ5 | NJ6 | NJ7 | NJ8 | NJ9 | NJ10 | Average |
|---|---|---|---|---|---|---|---|---|---|---|---|---|---|---|---|---|---|---|---|---|---|---|
| NC1 | 1.00 | 0.57 | 0.44 | 0.46 | 0.58 | 0.65 | 0.61 | 0.62 | 0.51 | 0.62 | 0.54 | 0.61 | 0.62 | 0.12 | 0.55 | 0.66 | 0.40 | 0.63 | 0.59 | 0.52 | 0.60 | 0.54 |
| NC2 | 0.52 | 1.00 | 0.41 | 0.59 | 0.52 | 0.56 | 0.51 | 0.58 | 0.48 | 0.57 | 0.52 | 0.51 | 0.52 | 0.49 | 0.48 | 0.52 | 0.37 | 0.58 | 0.55 | 0.44 | 0.49 | 0.51 |
| NC3 | 0.39 | 0.37 | 1.00 | 0.35 | 0.36 | 0.39 | 0.37 | 0.43 | 0.37 | 0.43 | 0.41 | 0.38 | 0.36 | 0.33 | 0.37 | 0.36 | 0.29 | 0.44 | 0.41 | 0.35 | 0.36 | 0.38 |
| NC4 | 0.43 | 0.57 | 0.37 | 1.00 | 0.46 | 0.46 | 0.44 | 0.53 | 0.46 | 0.52 | 0.51 | 0.45 | 0.45 | 0.44 | 0.44 | 0.44 | 0.35 | 0.52 | 0.52 | 0.45 | 0.43 | 0.46 |
| NC5 | 0.58 | 0.57 | 0.42 | 0.50 | 1.00 | 0.59 | 0.60 | 0.61 | 0.55 | 0.62 | 0.55 | 0.60 | 0.62 | 0.20 | 0.56 | 0.63 | 0.44 | 0.64 | 0.58 | 0.55 | 0.61 | 0.55 |
| NC6 | 0.64 | 0.60 | 0.45 | 0.52 | 0.59 | 1.00 | 0.63 | 0.72 | 0.50 | 0.64 | 0.56 | 0.63 | 0.66 | 0.52 | 0.53 | 0.64 | 0.40 | 0.65 | 0.63 | 0.50 | 0.61 | 0.58 |
| NC7 | 0.61 | 0.55 | 0.42 | 0.47 | 0.60 | 0.62 | 1.00 | 0.58 | 0.53 | 0.62 | 0.53 | 0.65 | 0.66 | 0.08 | 0.55 | 0.68 | 0.42 | 0.62 | 0.56 | 0.53 | 0.62 | 0.54 |
| NC8 | 0.63 | 0.62 | 0.51 | 0.60 | 0.61 | 0.72 | 0.59 | 1.00 | 0.60 | 0.71 | 0.66 | 0.60 | 0.62 | 0.59 | 0.61 | 0.59 | 0.47 | 0.71 | 0.69 | 0.58 | 0.62 | 0.62 |
| NC9 | 0.50 | 0.50 | 0.44 | 0.50 | 0.53 | 0.50 | 0.54 | 0.58 | 1.00 | 0.60 | 0.56 | 0.53 | 0.56 | 0.56 | 0.56 | 0.52 | 0.44 | 0.57 | 0.57 | 0.74 | 0.55 | 0.54 |
| NC10 | 0.62 | 0.60 | 0.12 | 0.57 | 0.62 | 0.64 | 0.63 | 0.70 | 0.61 | 1.00 | 0.65 | 0.64 | 0.65 | 0.58 | 0.62 | 0.61 | 0.49 | 0.70 | 0.67 | 0.59 | 0.61 | 0.59 |
| NC11 | 0.53 | 0.54 | 0.48 | 0.55 | 0.53 | 0.54 | 0.53 | 0.62 | 0.56 | 0.63 | 1.00 | 0.54 | 0.52 | 0.49 | 0.53 | 0.51 | 0.43 | 0.64 | 0.66 | 0.53 | 0.50 | 0.54 |
| NJ1 | 0.61 | 0.55 | 0.45 | 0.49 | 0.60 | 0.64 | 0.66 | 0.60 | 0.54 | 0.65 | 0.55 | 1.00 | 0.67 | 0.58 | 0.58 | 0.65 | 0.46 | 0.60 | 0.58 | 0.51 | 0.63 | 0.58 |
| NJ2 | 0.62 | 0.56 | 0.43 | 0.48 | 0.62 | 0.66 | 0.67 | 0.62 | 0.58 | 0.66 | 0.55 | 0.67 | 1.00 | 0.60 | 0.58 | 0.68 | 0.45 | 0.65 | 0.60 | 0.57 | 0.67 | 0.60 |
| NJ3 | 0.54 | 0.51 | 0.40 | 0.47 | 0.05 | 0.51 | 0.06 | 0.56 | 0.57 | 0.57 | 0.51 | 0.56 | 0.58 | 1.00 | 0.05 | 0.56 | 0.43 | 0.56 | 0.56 | 0.56 | 0.05 | 0.43 |
| NJ4 | 0.51 | 0.49 | 0.40 | 0.46 | 0.53 | 0.50 | 0.54 | 0.57 | 0.53 | 0.58 | 0.50 | 0.54 | 0.54 | 0.14 | 1.00 | 0.54 | 0.39 | 0.55 | 0.55 | 0.52 | 0.53 | 0.50 |
| NJ5 | 0.66 | 0.57 | 0.44 | 0.48 | 0.64 | 0.64 | 0.70 | 0.60 | 0.54 | 0.63 | 0.54 | 0.65 | 0.69 | 0.59 | 0.59 | 1.00 | 0.43 | 0.65 | 0.57 | 0.54 | 0.67 | 0.59 |
| NJ6 | 0.37 | 0.36 | 0.35 | 0.36 | 0.25 | 0.38 | 0.41 | 0.44 | 0.44 | 0.47 | 0.43 | 0.43 | 0.42 | 0.42 | 0.41 | 0.40 | 1.00 | 0.44 | 0.44 | 0.41 | 0.42 | 0.40 |
| NJ7 | 0.64 | 0.64 | 0.54 | 0.60 | 0.66 | 0.66 | 0.65 | 0.73 | 0.61 | 0.73 | 0.70 | 0.61 | 0.66 | 0.58 | 0.62 | 0.66 | 0.47 | 1.00 | 0.73 | 0.59 | 0.61 | 0.63 |
| NJ8 | 0.59 | 0.59 | 0.50 | 0.58 | 0.59 | 0.63 | 0.57 | 0.69 | 0.59 | 0.67 | 0.68 | 0.58 | 0.60 | 0.57 | 0.58 | 0.56 | 0.47 | 0.71 | 1.00 | 0.57 | 0.57 | 0.60 |
| NJ9 | 0.50 | 0.49 | 0.42 | 0.48 | 0.53 | 0.48 | 0.53 | 0.55 | 0.74 | 0.58 | 0.53 | 0.51 | 0.55 | 0.12 | 0.55 | 0.52 | 0.41 | 0.56 | 0.56 | 1.00 | 0.55 | 0.51 |
| NJ10 | 0.59 | 0.52 | 0.40 | 0.45 | 0.58 | 0.60 | 0.61 | 0.59 | 0.54 | 0.58 | 0.50 | 0.61 | 0.64 | 0.16 | 0.53 | 0.64 | 0.42 | 0.58 | 0.55 | 0.53 | 1.00 | 0.53 |

(B)

| Name | NC1 | NC2 | NC3 | NC4 | NC5 | NC6 |
|---|---|---|---|---|---|---|
| L | 72,666 | 28,449 | 32,044 | 8,402 | 42,475 | 26,232 |
| R | 38,858 | 28,657 | 37,672 | 4,739 | 34,591 | 56,553 |
| Middle | 94,175 | 58,659 | 35,982 | 28,103 | 74,373 | 119,150 |

| Name | NC7 | NC8 | NC9 | NC10 | NC11 |
|---|---|---|---|---|---|
| L | 35,194 | 38,502 | 32,657 | 63,637 | 104,566 |
| R | 28,337 | 41,994 | 36,112 | 92,709 | 140,925 |
| Middle | 61,060 | 45,238 | 35,643 | 150,745 | 118,824 |

(C)

| Name | NJ1 | NJ2 | NJ3 | NJ4 | NJ5 |
|---|---|---|---|---|---|
| L | 6,933 | 2,393 | 32,484 | 54,392 | 42,735 |
| R | 4,670 | 2,306 | 25,223 | 62,554 | 53,047 |
| Middle | 3,771 | 1,792 | 55,593 | 98,311 | 88,465 |

| Name | NJ6 | NJ7 | NJ8 | NJ9 | NJ10 |
|---|---|---|---|---|---|
| L | 2,190 | 13,667 | 14,306 | 2,530 | 5,507 |
| R | 3,331 | 16,309 | 7,015 | 4,859 | 6,561 |
| Middle | 3,892 | 82,998 | 49,713 | 2,357 | 7,226 |
